# EWSR1’s visual modalities are defined by its association with nucleic acids and RNA polymerase II

**DOI:** 10.1101/2023.08.16.553246

**Authors:** Soumya Sundara Rajan, Vernon J. Ebegboni, Patricio Pichling, Katelyn R. Ludwig, Tamara L. Jones, Raj Chari, Andy Tran, Michael J. Kruhlak, Jadranka Loncarek, Natasha J. Caplen

## Abstract

We report systematic analysis of endogenous EWSR1’s cellular organization. We demonstrate that EWSR1, which contains low complexity and nucleic acid binding domains, is present in cells in faster and slower-recovering fractions, indicative of a protein undergoing both rapid exchange and longer-term interactions. The employment of complementary high-resolution imaging approaches shows EWSR1 exists in in two visual modalities, a distributed state which is present throughout the nucleoplasm, and a concentrated state consistent with the formation of foci. Both EWSR1 visual modalities localize with nascent RNA. EWSR1 foci concentrate in regions of euchromatin, adjacent to protein markers of transcriptional activation, and significantly colocalize with phosphorylated RNA polymerase II. Interestingly, EWSR1 and FUS, another FET protein, exhibit distinct spatial organizations. Our results contribute to bridging the gap between our understanding of the biophysical and biochemical properties of FET proteins, including EWSR1, their functions as transcriptional regulators, and the participation of these proteins in tumorigenesis and neurodegenerative disease.

**SUMMARY:** Rajan et al. report the visualization of endogenous EWSR1. EWSR1 exists in two visual modalities in the nucleoplasm, one distributed and one as foci. Both EWSR1 modalities localize with nascent RNA. EWSR1 foci concentrate in regions of euchromatin and colocalize with phosphorylated RNA polymerase II.

## INTRODUCTION

EWSR1 (Ewing sarcoma breakpoint region) is a member of the FET family of nucleic acid proteins that includes FUS (Fused in sarcoma) and TAF15 (TATA-box binding protein associated factor-15) (reviewed in Lee et al., 2019; Schwartz et al., 2015). Functional protein domains common to the FET proteins include a zinc finger domain, a highly conserved RNA recognition motif (RRM), multiple low complexity domains (LCDs), and a C-terminal nuclear localization sequence (Ohno et al., 1994; Zakaryan and Gehring, 2006) (**Figure 1A**). These structural features contribute to EWSR1’s functional interactions with nucleic acids and proteins. For example, examination of EWSR1’s DNA binding activity employing chromatin-based immunoprecipitation (IP) have shown that EWSR1 preferentially binds at the transcriptional start sites of actively transcribed genes and downstream of the poly-A signal (Luo et al., 2015). Furthermore, examination of EWSR1’s protein-protein interactions have shown it binds RNA polymerase II (RNA pol II) (Bertolotti et al., 1998) specifically, the C-terminal domain (CTD), and like FUS, can regulate the phosphorylation of RNA pol II (Gorthi et al., 2018; Schwartz et al., 2012). Transcriptome-wide analysis of EWSR1 has also shown its binding of thousands of transcripts (Hoell et al., 2011); however, these and similar studies of FUS’s binding of RNA (Hoell et al., 2011; Rogelj et al., 2012; Wang et al., 2015) indicate that EWSR1 binds RNA irrespective of sequence or structure, suggesting that it can interact with multiple sites within an RNA and can function at various stages in the processing of RNA. Interestingly, while these studies suggested EWSR1 binds RNA irrespective of structure, more recent *in vitro* analysis of the interaction of EWSR1 with RNA and DNA in different structural conformations have shown that EWSR1’s RRM domain, and the low complexity Arg/Gly/Gly domains adjacent to the RRM, along with the zinc finger domain promote binding of RNA stem loops and G-rich DNA sequences folded in a quadruplex state (Selig et al., 2023). Furthermore, the entire EWSR1 protein (Pan et al., 2020) and EWSR1’s RRM and Arg/Gly/Gly domains (Selig et al., 2023) can bind the hybrid RNA:DNA structures that typify R-loops and that these interactions may have functional consequences (Gorthi et al., 2018).

**Figure 1:**
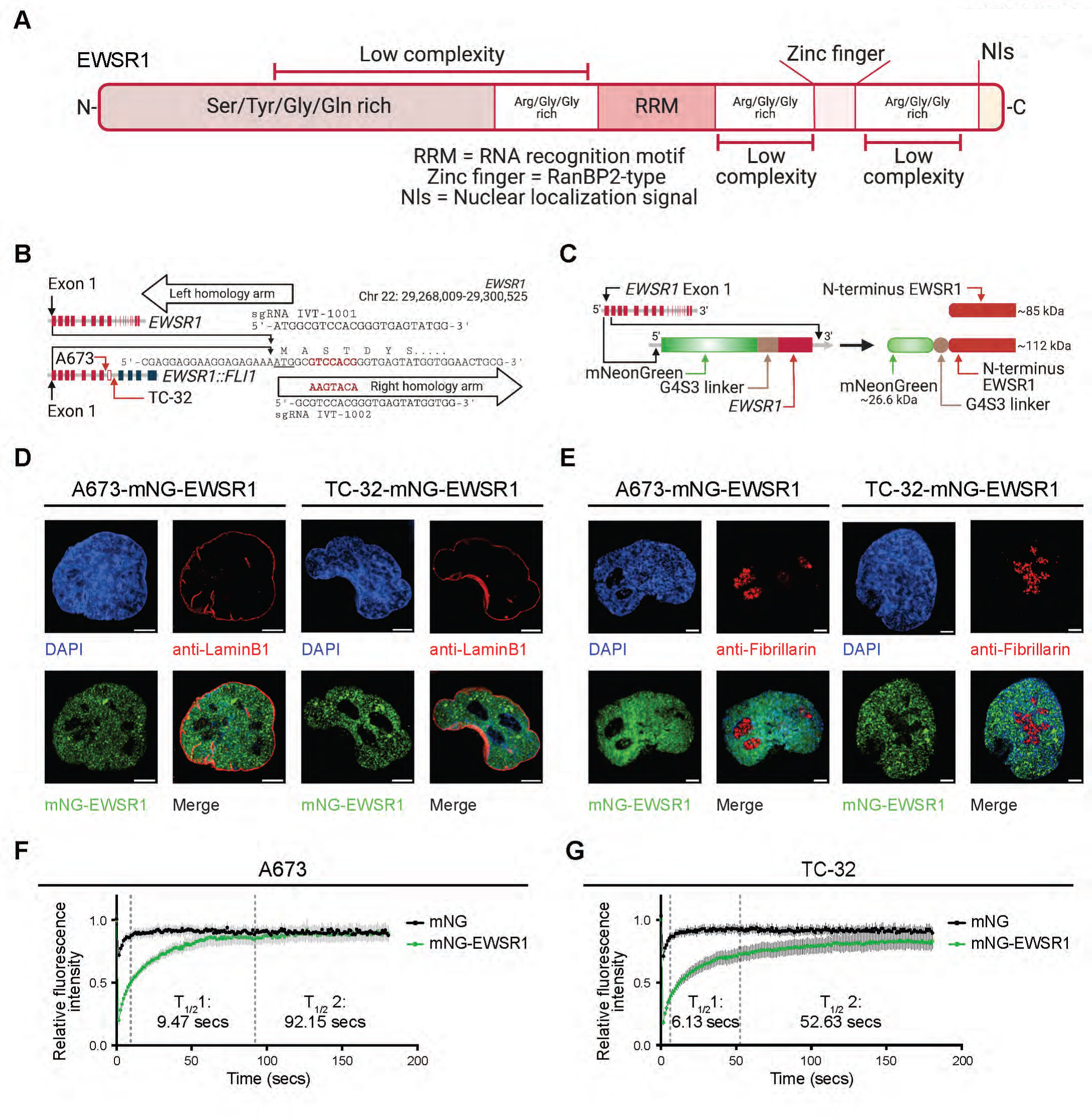
Endogenous EWSR1 exists in two kinetic fractions. (**A**) A schematic of EWSR1’s protein domains. (**B**) Schematic of the CRISPR-Cas9 strategy used to insert DNA elements into the *EWSR1* genomic locus of EWS cells immediately upstream of *EWSR1* exon 1. Indicated in red text is the minor sequence change within *EWSR1* exon 1 generated through the employment of the right homology arm donor sequences. Red arrows indicate the A673 and TC-32 *EWSR1* intron-7 and intron-8 breakpoints present in the rearranged *EWSR1* alleles, respectively. (**C**) Schematic representation of the introduction of the mNeonGreen (mNG) fluorescent reporter with the G4S3 linker donor cassette at the 5’ end of the *EWSR1* gene and the N-terminus of the expected modified protein. (**D**) Confocal microscopy images of A673-mNG-EWSR1 and TC-32-mNG-EWSR1 nuclei: Blue – DAPI, green – mNG-EWSR1, red – anti-Lamin B1, a nuclear membrane marker, and merged. Scale bar = 5 µm. Representative of 30 images across three independent experiments. (**E**) Confocal microscopy images of A673-mNG-EWSR1 and TC-32-mNG-EWSR1 nuclei: Blue – DAPI, green – mNG-EWSR1, red – anti-Fibrillarin, a nucleoli marker, and merged. Scale bar = 5 µm. Representative of 30 images across three independent experiments. (**F**) Relative fluorescence intensity after photo-bleaching of A673-mNG-EWSR1 cells and A673-mNG cells (unfused mNG) (n=10, mean ± SEM) and the derived dissociation coefficients. (**G**) Relative fluorescence intensity after photo-bleaching of TC-32-mNG-EWSR1 cells and TC-32-mNG cells (unfused mNG) (n=10, mean ± SEM) and derived dissociation coefficients. **A, B**, and **C** generated using BioRender.

Study of EWSR1’s function also needs to consider the biophysical properties of its multiple LCDs, particularly its N-terminus region. For example, recent *in vitro* analysis of EWSR1’s N-terminus of (residues 1-264) coupled with predictive modeling has shown that EWSR1’s Ser/Tyr/Gly/Gln-rich LCD is unstructured, adopting polyproline II helix and random coil conformations (Johnson et al., 2022). This finding is functionally relevant as many studies have demonstrated how dynamic nucleic-acid-protein assemblies that form membrane-less biomolecular condensates involve proteins containing LCDs, such as those in the FET proteins (reviewed in Kilgore and Young, 2022; McSwiggen et al., 2019; Schwartz et al., 2015). In this context, the investigation of the biophysical properties of the FET proteins’ LCDs have proven highly informative in establishing the parameters of biomolecular condensate formation and has informed the study of the concept that dynamic nucleic acid-protein interactions can promote the assembly of functional compartments within cells (Burke et al., 2015; Han et al., 2012; Kato et al., 2012; Maharana et al., 2018; Murthy et al., 2021; Shimobayashi et al., 2021). For example, the binding of FUS to RNA promotes its interaction with the CTD of RNA pol II (Burke et al., 2015; Schwartz et al., 2013).

The studies discussed above have, in many cases by necessity, predominantly used biochemical assays or ectopically expressed tagged versions of the FET proteins at non-physiological concentrations to study their function (Altmeyer et al., 2015; Gorthi et al., 2018; Johnson et al., 2022; Maharana et al., 2018; Patel et al., 2015; Schwartz et al., 2012; Schwartz et al., 2013; Shin et al., 2017). Studies using these experimental approaches have proven critical to assessing the biochemical and biophysical properties of the many interactions the FET proteins undergo (Burke et al., 2015; Selig et al., 2023; Shimobayashi et al., 2021); however, it remains to be determined to what extent these findings reflect the function of these proteins within the nucleus of a transcriptionally active cell. In this study, we have generated two EWSR1-fluorescent EWS reporter lines and have conducted a systematic high-resolution microscopy-based localization study of endogenous EWSR1 using cell line models of the EWSR1::FLI1 fusion-driven tumor Ewing sarcoma (EWS) that harbor one non-rearranged *EWSR1* allele (Giard et al., 1973; Martinez-Ramirez et al., 2003; Whang-Peng et al., 1986). Using 3D-structured illumination microscopy (3D-SIM) and stimulated emission depletion (STED) microscopy, we demonstrate the distribution of a large proportion of EWSR1 throughout the nucleoplasm, with the remaining fraction of EWSR1 protein detected as foci. We also report the spatial organization of EWSR1 in nuclear space relative to nucleic acids and exemplar proteins that function in genome organization or transcription. Our results show that EWSR1 foci overlap significantly with phosphorylated RNA polymerase II, demonstrating in cells the critical association of EWSR1 with sites of active transcription. Furthermore, we provide visual evidence that FUS and EWSR1 do not share the same nucleoplasmic configuration suggesting that these proteins may have distinct functions based on their specific spatial organization within nuclei.

## RESULTS

### The generation of mNG-EWSR1 reporter cell line

To generate reporter cell lines of endogenous EWSR1, we used CRISPR-Cas9 gene editing to insert the mNeonGreen (mNG) reporter gene (Shaner et al., 2013) into a site at the 5’ end of *EWSR1* exon 1 (**Figure 1B**) of two EWS cell lines - A673 and TC-32. **Figure 1C** shows a schematic of the DNA cassette inserted into the *EWSR1* locus, the targeting guide RNAs, and the N-terminus of the predicted fusion protein. Following single cell cloning, we confirmed mNG protein expression using flow cytometry. Representative results for two A673 and two TC-32 clones demonstrated similar fluorescent signals in the clones derived from each cell line, with the TC-32-mNG positive clones expressing a higher median fluorescence signal than the A673-mNG positive clones (**Figure S1A**). We anticipated that the dependency of EWS cells on EWSR1::FLI1 function would preclude identification of any clones in which we had modified the re-arranged allele, but as one approach to confirm the fusion of the mNG reporter to EWSR1, we transfected single cell clones expressing >90% green fluorescence with either a siRNA targeting the full-length *EWSR1* transcript (siEWSR1; s4866) or one targeting the *FLI1* region of the *EWSR1::FLI1* fusion mRNA (siFLI1; s5266) (**Figure S1B**) and performed flow cytometry (**Figure S1C**). As expected, we identified no clones expressing an mNG-modified EWSR1::FLI1 protein; however, based on the shift in fluorescence following transfection of siEWSR1, but not siFLI1, we identified several EWSR1-modified clones, and **Figure S1C** shows representative analysis of two A673 (4 and 9) clones and two TC-32 (6 and 17) clones. Image-based assessment of A673-mNG clone 4 and TC-32-mNG clone 6 demonstrated nuclear localization of the mNG-EWSR1 protein, and we also observed a significant reduction in nuclear fluorescent signal following transfection of the *EWSR1* siRNA (**Figure S1D**). Sequence analysis of A673-mNG clone 4 and TC32-mNG clone 6 (**Supplementary file 1**) confirmed the correct insertion of the mNG containing DNA cassette in-frame with *EWSR1*. The parental and modified cells exhibited similar growth kinetics (**Figure S1E**). For our subsequent studies, we used A673-mNG clone 4 and TC32-mNG clone 6, henceforth designated as A673-mNG-EWSR1 and TC-32-mNG-EWSR1, respectively. We next used super-resolution microscopy to assess the sub-cellular localization of the mNG-EWSR1 protein expressed in the modified EWS cell lines (**Figures 1D** and **E**). We observed mNG-EWSR1’s predominate localization to the nucleoplasm of non-dividing cells (**Figure 1D**), with little or no localization in nucleoli (indicated by Fibrillarin immunofluorescence (IF), **Figure 1E**).

### Endogenous EWSR1 exists in two kinetic fractions

Determination of the rate of recovery upon photo-bleaching can offer insights into the kinetics of a protein within a cellular compartment and, thus, the dynamics of its interactions with other molecules (McNally, 2008; Stavreva and McNally, 2004; Thompson et al., 1981). Therefore, we next assessed the mobility of full length EWSR1 in our reporter cell lines. For this analysis, we used the mNG-EWSR1 expressing cells and isogenic control cell lines (A673-mNG-P2A-HiBiT-EWSR1 and TC-32-mNG-P2A-HiBiT-EWSR1) expressing cleaved, unfused mNG (**Figures S1F - I)**. We generated the isogenic control cell lines using the same gene editing approach as employed for the mNG-EWSR1 expressing cell lines but used a DNA cassette that includes a high-efficiency P2A self-cleavage peptide present between the mNG reporter gene and a HiBiT peptide tag (**Figure S1F**). Flow cytometry (**Figure S1G**), immunoblot (**Figure S1H**) and image-based analyses (**Figure S1I**) demonstrate collectively that these cell lines express unfused mNG (**Figure S1I**) (Kim et al., 2011) Analysis of the fluorescence recovery after photobleaching (FRAP) (Zeiss Airyscan) recovery curves demonstrated that EWSR1 is present in both a faster-recovering fraction, as indicated by T_1/2_ values of less than 10 secs, and in a slower recovering fraction, as indicated by T_1/2_ values of over 50 secs (**Figures 1F** and **1G**). These values are relative to the unfused mNG control that is freely diffusing and fits a mono-exponential recovery curve. Our results are consistent with the endogenous EWSR1 protein existing in two kinetic states within cells, one in which the protein is undergoing long term interactions and one consisting of rapidly dynamic interactions. We hypothesize that these two kinds of interactions reflect EWSR1’s varied binding abilities with different partners such as longer-term interactions with another macromolecule (e.g., itself or another protein) and dynamic transient interactions, with nucleic acids or proteins.

### Endogenous EWSR1 organizes as two visual modalities

Super-resolution microscopy-based visualization of mNG-EWSR1 in the modified A673 and TC-32 (**Figure S2A**) cells (Nikon SoRa spinning disk super-resolution microscope) show a range of fluorescence intensities throughout the nucleoplasm. To assess mNG-EWSR1’s nucleoplasmic organization at a higher resolution, we next employed 3D-SIM. At this resolution, we confirmed the presence of a range of mNG-EWSR1 intensities across the nucleoplasm in both modified cell lines (**Figures 2A** and **B**). As a means of depicting the intra-nucleoplasmic distribution of mNG-EWSR1 fluorescence, we plotted fluorescence intensity across a defined region of a representative nuclei and the indicated insets. These line plots reveal the varied distribution of mNG-EWSR1 intensities across distance. Critically, the Y-axis of these intensity plots reflect the cell line-specific differences in protein expression levels detected by FACS analysis **(Figure S1A)**. As represented in the line plots, we observe the presence of two patterns of mNG-EWSR1 signal – a lower intensity “distributed” signal and peaks of high intensity signal consistent with the concentration of EWSR1 protein as foci. These two visual modalities of mNG-EWSR1 occur throughout the volume of nucleus as depicted via representative z-steps (**Figure 2C)**.

**Figure 2:**
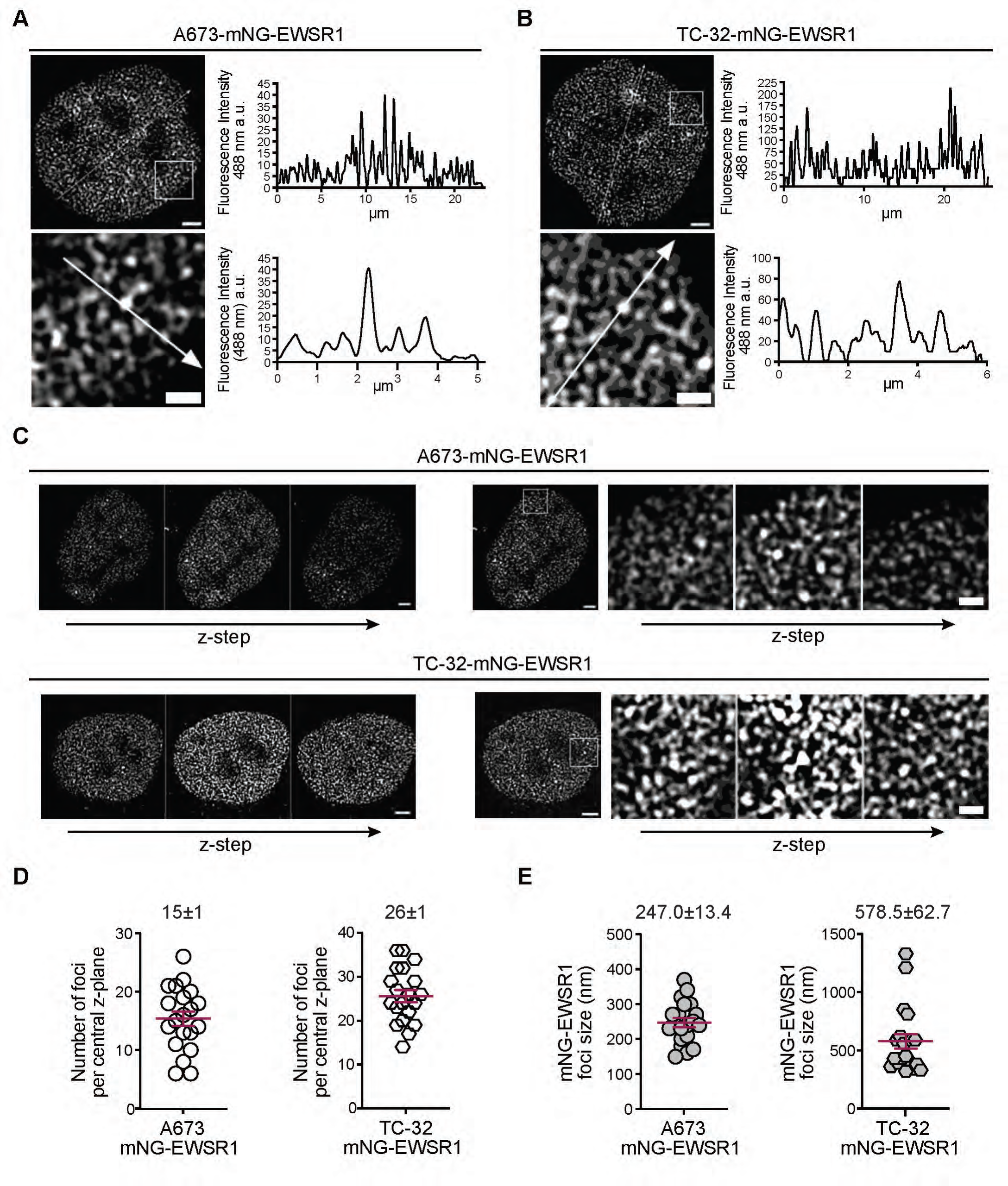
Endogenous EWSR1 organizes as two visual modalities. (**A**) A 3D-SIM image of an A673-mNG-EWSR1 nucleus, with the indicated line and corresponding line plot and the indicated inset. Inset shows line and the corresponding intensity plot. Representative of 100 images across ten independent experiments. (**B**) A 3D-SIM image of a TC-32-mNG-EWSR1 nucleus, with the indicated line and corresponding line plot and the indicated inset. Inset shows line and the corresponding intensity plot. Representative of 100 images across ten independent experiments. (**C**) Representative z-steps of a 3D-SIM reconstructed images obtained using A673-mNG-EWSR1 cells (upper panels) and TC-32 mNG-EWSR1 cells (lower panels). The panels on the left show z-steps of a 3D-SIM reconstructed image of a whole nuclei. The middle panels, the central plane of a different representative nuclei indicating the inset used for the representative z-stacks shown on the right. Representative of 20 images across ten independent experiments. **(D)** Number of mNG-EWSR1 foci per central z-plane of (n=20) in A673- and TC-32-mNG-EWSR1 nuclei **(E)** mNG-EWSR1 foci size (n=20) obtained using A673-mNG-EWSR1 cells and TC-32-mNG-EWSR1 cells. (**D** and **E)** Thresholds applied to define mNG-EWSR1 foci: three times greater intensity than background and a size restriction of greater than 20 nm; red lines indicate the mean ± SEM. Nucleus: scale bar = 3 µm; Inset: scale bar = 1 µm.

The detection of a proportion of the mNG-EWSR1 protein as foci could represent protein aggregates or detection of increased EWSR1 protein at locations of functional relevance. To understand the nucleoplasmic distribution of mNG-EWSR1 foci further, we defined analytic criteria we could apply across both modified cell lines under different experimental and imaging conditions. Specifically, we applied a signal threshold of at least three times the intensity above the background subtracted signal and a size threshold (20 nm) that disregards signal intensities below the pixel size (see Methods). Post-application of these thresholds, we quantified about 15 foci per central z-stack of an A673 mNG-EWSR1 nuclei and ∼25 foci per central z-stack of TC-32 mNG-EWSR1 nuclei **(Figure 2D)**. Foci size varies from about 200 to 1000 nm, with TC-32 mNG-EWSR1 showing more variation in foci size than A673 mNG-EWSR1 cells **(Figure 2E).**

To substantiate the results generated using the mNG-fused EWSR1, we assessed EWSR1 expression and organization in unmodified A673 and TC-32 cells using an antibody against EWSR1 **(Figure S2B**) and 3D-SIM (**Figures S2C** and **D**). Consistent with our analysis of the relative expression of mNG-EWSR1 in A673 and TC-32 measured by flow cytometry (**Figure S1A**), immunoblot analysis demonstrated that TC-32 cells express more EWSR1 protein than A673 cells (**Figure S2B**). Detection of EWSR1 using IF and 3D-SIM confirmed its nucleoplasmic distribution and corresponding line plots demonstrate variations in signal intensities across representative nuclei and regions within nuclei (**Figures S2C** and **D**). However, unlike our assessment of mNG-EWSR1 fluorescence, the antibody-based fluorescence intensities did not reflect the cell line-specific differences in the expression levels of EWSR1 protein in A673 and TC-32 (Y-axis). Quantification of the number of foci per central z-plane showed concordance with that observed for mNG-EWSR1 (**Figure S2E**). The size of EWSR1 foci detected by IF in A673 cells also accorded with that determined using the mNG-EWSR1 protein; however, analysis of TC-32 cells using the EWSR1 antibody did not detect the foci over 400 nm (**Figure S2F**). One explanation for this discrepancy is the inability of the antibody to penetrate larger foci. Nonetheless, our comparison of the detection of EWSR1 by IF with the visualization of EWSR1 using a fluorescent reporter supports using the A673 and TC-32 mNG-EWSR1 cell lines as resources for the study of endogenous EWSR1.

### Endogenous EWSR1’s localization relative to DNA and nascent RNA

EWSR1’s structural domains (**Figure 1A**) and previous studies have highlighted its potential to interact with nucleic acids (Luo et al., 2015; Schwartz et al., 2013; Selig et al., 2023; Takahama et al., 2011; Wang et al., 2015); however, we lack a detailed examination of the spatial organization of these interactions within cells. We first assessed the localization of mNG-EWSR1 and DNA using the DAPI fluorescent dye at the resolution limits of 3D-SIM (∼120 nm); see Figure S3 for representative images demonstrating the use of TetraSpeck microspheres that ensured correct channel alignments after image reconstruction of all multi-channel images presented in this study. **Figure 3A** shows merged images of mNG-EWSR1 (green) and DNA (blue) of a representative A673 nucleus, and **Figure 3B** shows images of a representative TC-32 nucleus. These images, including the indicated region within each nucleus, show mNG-EWSR1 and DAPI as discrete, non-overlapping signals. Further examination of mNG-EWSR1’s organization relative to DNA show its exclusion from condensed regions of the DNA as marked by intense DAPI staining, throughout the volume of the nucleus in both A673 mNG-EWSR1 cells (**Figure 3C**) and TC-32 mNG-EWSR1 cells (**Figure 3D**).

**Figure 3:**
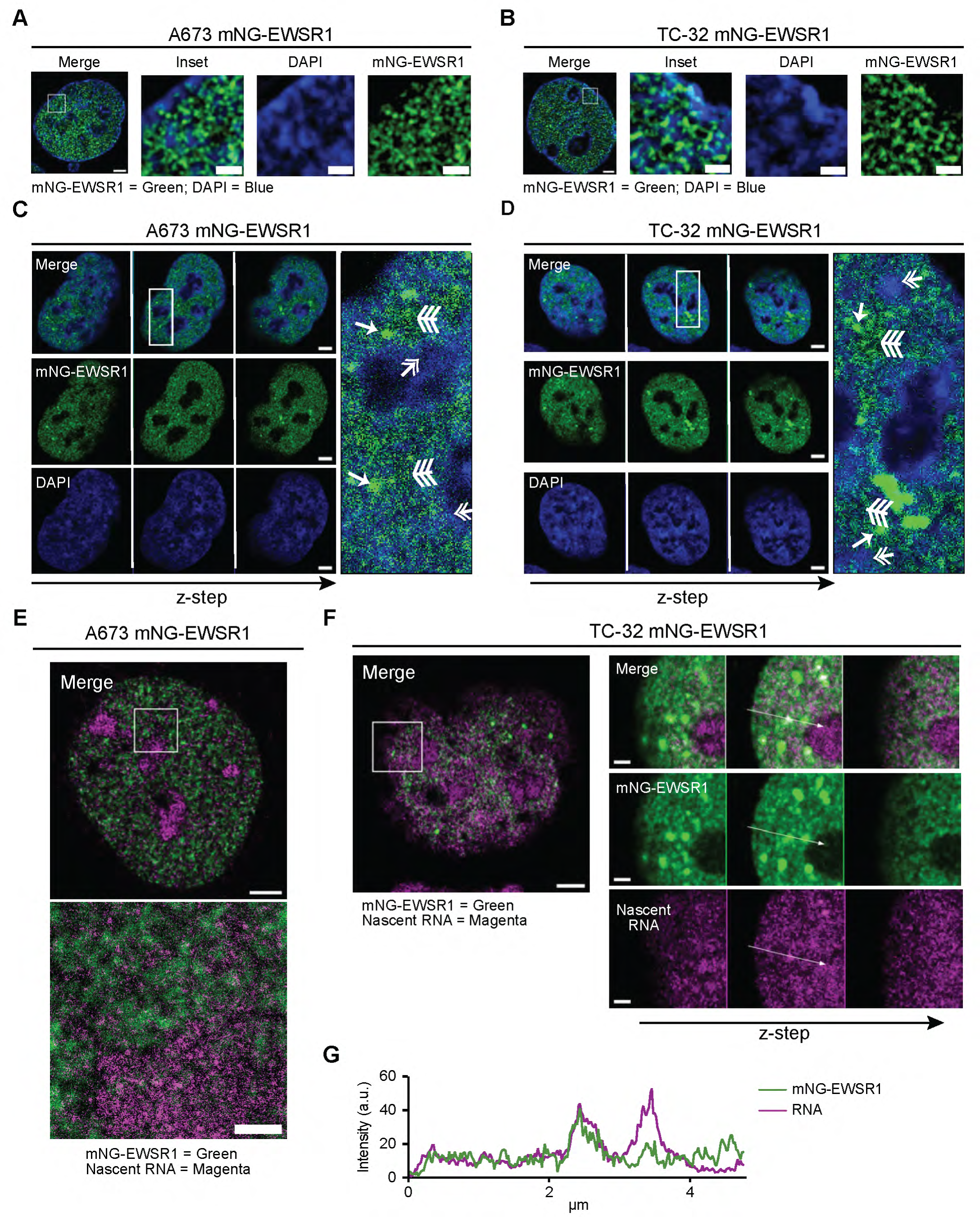
Endogenous EWSR1’s localization relative to DNA and nascent RNA. (**A**) Merged 3D-SIM images of an A673 mNG-EWSR1 nucleus showing mNG-EWSR1 and DNA marked by DAPI, with the inset indicated. (Scale bar = 3 µm). The inset shows an enlarged view of the indicated region, merged and single channels, respectively (Scale bar = 1 µm). Representative of 20 nuclei across three independent experiments. (**B**) Merged 3D-SIM images of a TC-32 mNG-EWSR1 nucleus showing mNG-EWSR1 and DNA marked by DAPI, with the inset indicated. (Scale bar = 3 µm). The inset shows an enlarged view of the indicated region, merged and single channel, respectively. Representative of 20 nuclei across three independent experiments. (Scale bar = 1 µm). **(C)** Representative z-plane images of a A673-mNG-EWSR1 nucleus showing mNG-EWSR1 and DAPI (merged and single channel). (Scale bar = 3 µm). Representative of 20 nuclei across three independent experiments. Inset marked on the central plane shows an enlarged merge image with single arrow head depicting mNG-EWSR1 foci, double arrow head depicting condensed DNA marked by excess DAPI staining and three arrowheads marking distributed mNG-EWSR1. **(D)** Representative z-plane images for a TC-32-mNG-EWSR1 nucleus showing mNG-EWSR1 and DAPI (merged and single channel). (Scale bar = 3 µm). Representative of 20 nuclei across three independent experiments. Inset marked on the central plane shows an enlarged merge image with single arrow head depicting mNG-EWSR1 foci, double arrow head depicting condensed DNA marked by excess DAPI staining and three arrowheads marking distributed mNG-EWSR1. (**E)** A 3D-SIM image (central z-plane) of a A673-mNG-EWSR1 nucleus showing images of mNG-EWSR1 and nascent RNA, with the inset indicated (Scale bar = 3 µm). The inset shows a merged STED-resolved view of the indicated region (mNG-EWSR1 (488 nm) green, fluorescently labeled nascent RNA (594 nm) magenta). **(F)** A merged image (central z-plane) of a TC-32-mNG-EWSR1 nucleus showing mNG-EWSR1 (488 nm, green) and (fluorescently labeled nascent RNA (594 nm, magenta with the indicated inset (Scale bar = 3 µm). The right-hand panels show representative z-steps for the indicated STED-resolved inset (merged and single channel) (Scale bar = 1 µm). Representative of 10 nuclei across two independent experiments. **(G)** Fluorescence intensity plots of the line profile indicate in Figure 3F.

To explore endogenous EWSR1’s association with RNA, we labeled nascent RNA by treating cells with 5-ethynyl uridine (EU) for 40 minutes and employed the Click-iT® RNA Alexa Fluor® 594 fluorochrome for its visualization. For these studies we used 2D-STED microscopy as it enables visualization of mNG-EWSR1 at a resolution of ∼220 nm and analysis of the Alexa Fluor® 594 incorporated RNA at a lateral resolution of ∼60 nm. Images of a representative A673-mNG-EWSR1 nucleus and the indicated inset (**Figure 3E**) show nascent RNA and mNG-EWSR1 occupy the same nucleoplasmic space excluding the nucleolus. We observe comparable results in TC-32 cells and show that this organization extends throughout the nucleoplasmic volume of the cell as represented by the representative z-steps (**Figure 3F**). As a complementary means of representing the spatial distributions of nascent RNA and mNG-EWSR1, **Figure 3G**, shows a line plot drawn across the indicated region of the inset presented in **Figure 3F**. This representative line plot demonstrates that the nascent RNA and mNG-EWSR1 maxima fluorescent signals at the central plane of the nucleus overlap. Interestingly, the overlap in signals shows close alignments of nascent RNA and both distributed EWSR1 and EWSR1 foci.

### EWSR1 foci coincide with euchromatic regions

Analysis of mNG-EWSR1’s locations relative to DNA and RNA are consistent with biochemical studies reporting EWSR1’s function in the regulation of gene expression, but to confirm that EWSR1 preferentially localizes to transcriptionally active regions of the genome, we next incorporated into our analysis detection of Histone H3, which, forms part of the nucleosome structure responsible for the higher-order organization of DNA. In brief, we visualized histone H3 using an antibody that detects the H3.1, H3.2, H3.3, and CENPA histone variants. We defined euchromatic regions based on the detection of antibody-accessible Histone H3 and low intensity DAPI (Ahmad and Henikoff, 2002; Jin and Felsenfeld, 2007). **Figure 4A** shows images of a representative nucleus and the indicated inset from A673-mNG-EWSR1 cells, and **Figure 4B**, the same for a representative TC-32-mNG-EWSR1 nucleus (merged and dual-colored images: DAPI – blue, Histone H3 – red, mNG-EWSR1 – green). The inset images and representative line plots indicate a region of euchromatin based on detection of elevated levels of antibody-accessible Histone H3 and low intensity DAPI and show multiple mNG-EWSR1 foci within these nucleoplasmic spaces. Our analysis of EWSR1’s localization relative to the nucleic acids suggest that both visual modalities of EWSR1 – distributed and foci – are components of the dynamic interactions required for transcriptional regulation, so we next examined EWSR1’s localization to relative well studied protein regulators of gene expression, specifically BRD4 and MED1.

**Figure 4:**
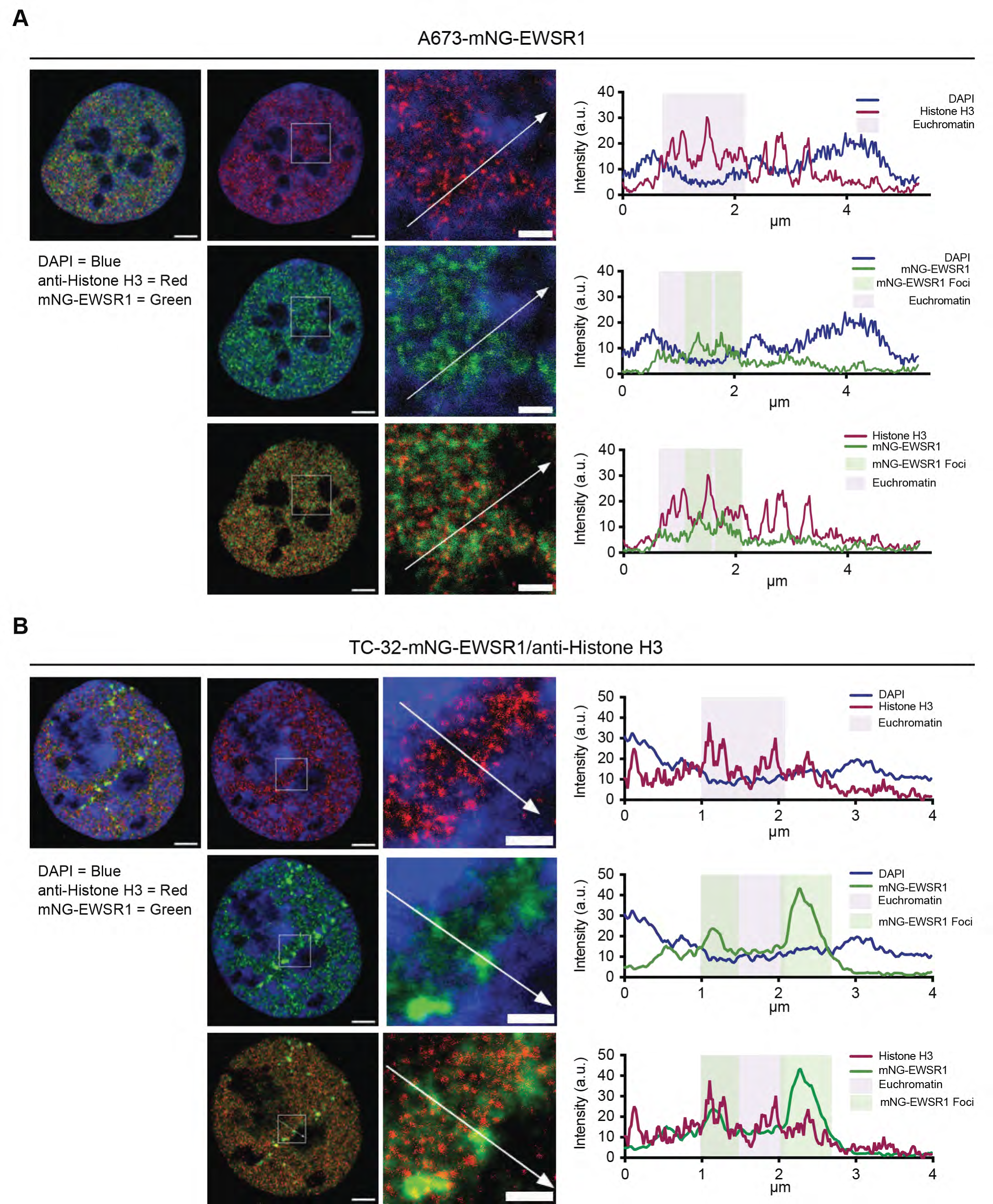
Regions of euchromatin and visualization of EWSR1 foci coincide. (**A**) Merged three- or two-color 3D-SIM images (central z-plane) of an A673-mNG-EWSR1 nucleus showing mNG-EWSR1 (green), Histone H3-IF (magenta) and DAPI (blue). The dual channel images show the indicated inset used for the enlarged views (Histone H3-IF resolved to 60 nm), and the lines used to generate the line profile shown on the right. (**B**) Merged three- or two-color 3D-SIM images (central z-plane) of a TC-32-mNG-EWSR1 nucleus showing mNG-EWSR1 (green), Histone H3-IF (magenta) and DAPI (blue). The dual channel images show the indicated inset used for the enlarged views (Histone H3-IF resolved to 60 nm), and the lines used to generate the line profile shown on the right. Nuclear scale bar = 3 µm and inset scale bar = 1 µm.

### EWSR1 foci are adjacent to transcriptional coactivators in nucleoplasmic space

Coactivators of transcription, including BRD4, a member of the bromodomain and extra-terminal (BET) family of proteins, and the mediator complex, are essential regulators of RNA polymerase II-driven transcription (Cho et al., 2018; Sabari et al., 2018; Zamudio et al., 2019). We, therefore, next examined EWSR1’s localization relative to BRD4, and a component of the mediator complex, MED1, using IF. **Figures 5A** and **B** show representative images of A673-mNG-EWSR1 and TC-32-mNG-EWSR1 nuclei, respectively, in which we detected BRD4 and Histone H3 by IF. Accompanying these images are STED-resolved images of the indicated regions and line plots generated using the indicated lines. Because STED microscopy can resolve Histone H3-IF, we used this as a marker of genomic organization rather than DAPI. The images and line plots show BRD4 exists in proximity to nuclear regions that are Histone H3 antibody accessible. Merged images of mNG-EWSR1 and BRD4 demonstrate the proximity of these two proteins in nucleoplasmic space with mNG-EWSR1 foci located adjacent to BRD4 signals as shown in the insets and corresponding line plots. We next resolved MED1 detected by IF up to 60 nm (**Figures 5C** and **5D**) and observed MED1 signal maxima flank mNG-EWSR1 foci. Collectively, these results suggest that mNG-EWSR1, BRD4, and MED1 are present in proximal 3D nuclear space, and provide evidence for the presence of EWSR1 foci at sites poised for the initiation of transcription.

**Figure 5:**
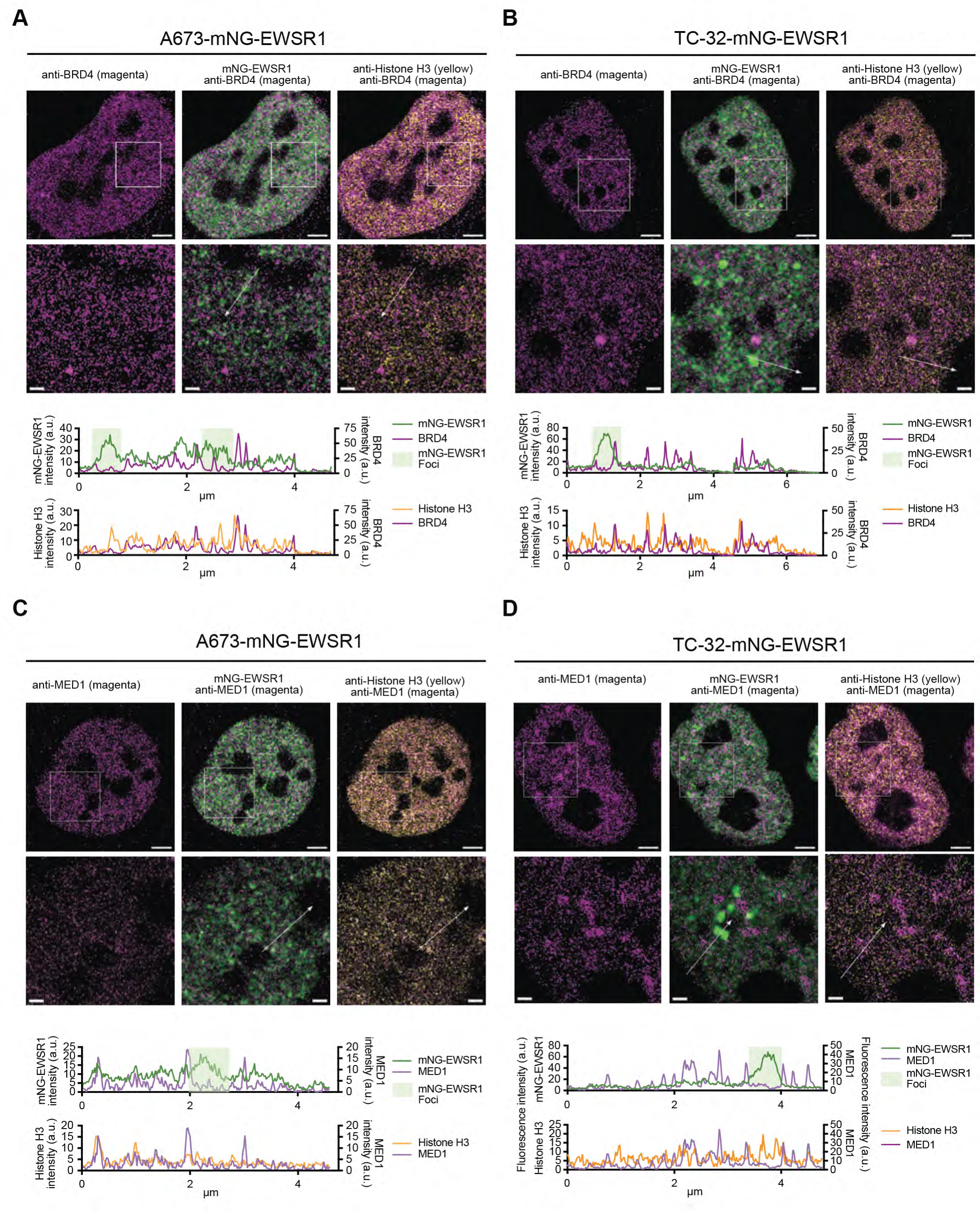
EWSR1 foci are present adjacent to transcriptional coactivators in nucleoplasmic space. (**A**) Single and dual-color 3D-SIM images of an A673-mNG-EWSR1 nucleus (BRD4-IF – magenta, mNG-EWSR1 – green, Histone H3-IF – yellow) (scale bar = 3 µm) and the indicated STED-resolved insets (scale bar = 1 µm), including the line used to generate the dual channel line profiles shown below. Representative of 20 images across two independent experiments. (**B**) Single and dual-color 3D-SIM images of a TC-32-mNG-EWSR1 nucleus (BRD4-IF – magenta, mNG-EWSR1 – green, Histone H3-IF – yellow) (scale bar = 3 µm) and the indicated STED-resolved insets (scale bar = 1 µm), including the line used to generate dual channel line profiles shown below. **(C)** Single and dual-color 3D-SIM images of an A673-mNG-EWSR1 nucleus (MED1-IF – magenta, mNG-EWSR1 – green, Histone H3-IF – yellow) (scale bar = 3 µm) and the indicated STED-resolved insets (scale bar = 1 µm), including the line used to generate dual channel line profiles shown below. Representative of 20 images across two independent experiments. **(D)** Single and dual-color 3D-SIM images of a TC-32-mNG-EWSR1 nucleus (MED1-IF – magenta, mNG-EWSR1 – green, Histone H3-IF – yellow) (scale bar = 3 µm) and the indicated STED-resolved insets (scale bar = 1 µm), including the line used to generate the dual channel line profiles shown below. Representative of 20 images across two independent experiments.

### EWSR1 concentrates at regions of active transcription

The regulation of transcription includes post-translational modification (PTM) of the CTD of RNA pol II (Harlen and Churchman, 2017). For example, phosphorylation of serine 5 (pS5-RNA pol II) is associated with transcriptional initiation, and transcriptional elongation is associated with the concurrent dephosphorylation of serine 5 and the phosphorylation of serine 2 (pS2-RNA pol II). One of the FET proteins’ best-characterized functions has focused on their interactions with RNA pol II and their contributions to the regulation of the serine 2 phosphorylation event (Schwartz et al., 2012). To extend these findings to a cellular context in the modified EWS cells, we first confirmed that the mNG-EWSR1 protein retained its interaction with RNA pol II (total and phosphorylated) using protein lysates prepared from wild type and modified A673 and TC-32 cells and immunoprecipitation (**Figure S4A**). Next, we visualized the localization of mNG-EWSR1 and phosphorylated RNA polymerase II (pS5 and pS2) by 3D-SIM, followed by STED microscopy. Consistent with the biochemical analysis of EWSR1’s interaction with RNA pol II, 3D-SIM fluorescent images of mNG-EWSR1 and the immunofluorescent detection of pS5-RNA pol II or pS2-RNA pol II demonstrate the proximity of these two proteins in A673-mNG-EWSR1 (**Figure 6A**) and TC-32 mNG-EWSR1 cells (**Figure 6B**), with significant overlap occurring at mNG-EWSR1 foci in both cell lines. We next aimed to quantify this colocalization. To do so, we used the previously established thresholds that define mNG-EWSR1 foci and performed a pixel-level analysis of the relative localizations of mNG-EWSR1 foci and either pS5-RNA pol II or pS2-RNA pol II. For this analysis, we calculated two statistical parameters, the overlap in fluorescent signals indicated by the determination of a Mander’s Overlap Coefficient (MOC) and the intersection of fluorescent signals by the calculation of a Pearson Correlation Coefficient (PCC) (Aaron et al., 2018; Dunn et al., 2011; Zinchuk and Grossenbacher-Zinchuk, 2014). Both coefficients resulted in mean values of >0.75, indicating that within the nucleoplasm, there is a significant association of mNG-EWSR1 foci with RNA pol II in states associated with either transcriptional initiation or transcriptional elongation. **(Figures 6C** and **6D)**. We next aimed to substantiate these findings using STED microscopy. Visualization of mNG-EWSR1 and pS5-RNA pol II or pS2-RNA pol II in modified A673 (**Figures S4C and S4D**) and TC-32 (**Figure 6E** and **F**) cells at this increased resolution confirmed the colocalization of mNG-EWSR1 foci with phosphorylated RNA pol II. This association is particularly clear when examined as line plots across the indicated inset regions and provide further evidence of the colocalization of EWSR1 foci with transcriptionally active RNA pol II.

**Figure 6:**
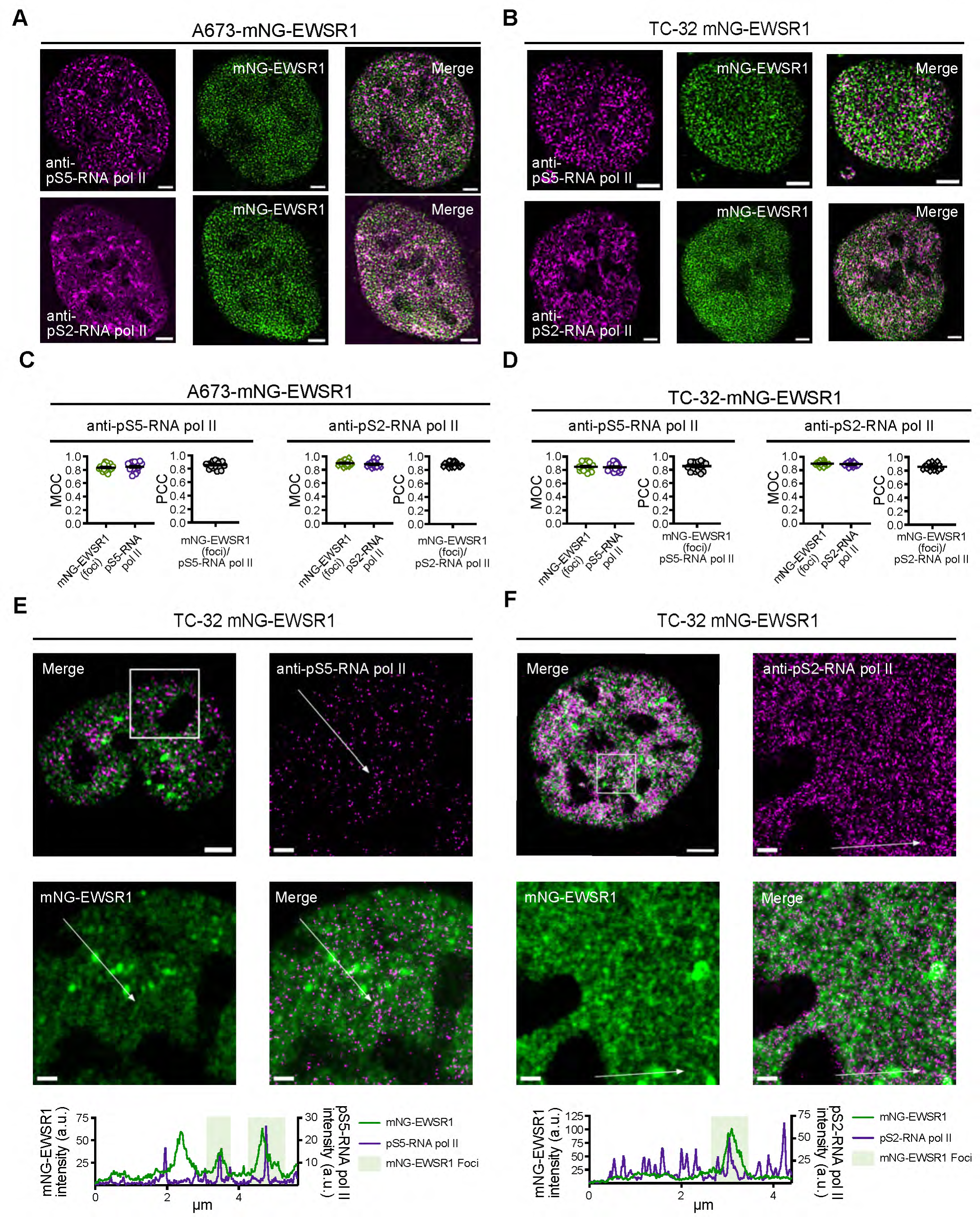
EWSR1 concentrates at regions of active transcription. (**A**) Single and merged 3D-SIM images of an A673 mNG-EWSR1 nucleus visualizing mNG-EWSR1and pS5-RNA pol II (IF) or mNG-EWSR1 and pS2-RNA pol II (IF) (Scale bar = 3 µm). Representative of 20 images across two independent experiments. (**B**) Single and merged 3D-SIM images of a TC-32 mNG-EWSR1 nucleus visualizing mNG-EWSR1and pS5-RNA pol II (IF) or mNG-EWSR1 and pS2-RNA pol II (IF) (scale bar = 3 µm). Representative of 20 images across two independent experiments. **(C)** The MOC and PCC analysis of mNG-EWSR1 foci and pS5-RNA pol II or pS2-RNA pol II (20 A673-mNG-EWSR1 nuclei, black lines indicate the mean ± SEM). **(D)** The MOC and PCC analysis of mNG-EWSR1 foci and pS5-RNA pol II or pS2-RNA pol II (20 TC-32-mNG-EWSR1 nuclei, black lines indicate the mean ± SEM). (**E**) Merged 3D-SIM image of a TC-32-mNG-EWSR1 nucleus (pS5-RNA pol II-IF – magenta, mNG-EWSR1 – green) (scale bar = 3 µm) and the indicated STED-resolved insets (scale bar = 1 µm), including the line used to the generate the dual channel line profile shown below. Representative of 20 nuclei across two independent replicates. **(F)** Merged 3D-SIM image of a TC-32-mNG-EWSR1 nucleus (pS2-RNA pol II-IF – magenta, mNG-EWSR1 – green) (scale bar = 3 µm) and the indicated STED-resolved insets (scale bar = 1 µm), including the line used to the generate the dual channel line profile shown below. Representative of 20 nuclei across two independent replicates.

### Dual CDC7/CDK9 inhibition changes the localization of EWSR1 associated with phosphorylated RNA polymerase II

The importance of the cellular localization of FET proteins, especially FUS, and their function, is evidenced by their redistribution to other cellular compartments in response to external stimuli, such as DNA damage (Mastrocola et al., 2013; Nogami et al., 2022; Rulten et al., 2014), or their aberrant localization in specific disease states (reviewed in Baradaran-Heravi et al., 2020; Harrison and Shorter, 2017; Mackenzie and Neumann, 2012; Rhoads et al., 2018). To assess if endogenous EWSR1 responds in a similar manner, we treated mNG-EWSR1 cells with PHA-767941, a dual CDC7/CDK9 inhibitor, that targets kinases that function in the regulation of DNA replication (CDC7) and the regulation of transcription (CDK9) (Montagnoli et al., 2008). Previously, Flores and colleagues reported that 2 µM PHA767941 shows EWS cell line-specific effects on transcriptional blockade (Flores et al., 2020). We therefore aimed to visualize the effects of this compound on EWSR1’s localization in our reporter cell lines. Using the same concentration of compound as the previous study, we first assessed PHA-767491’s effect on the cell cycle and the phosphorylation of RNA pol II (serine 2) post-treatment of unmodified A673 and TC-32 cells (**Figure S5A** and **B)**. At the time points assessed (1, 6 and 16 hours), we observed minimal effect on cell cycle following addition of PHA-767491, but we did see decreased detection of pS2-RNA pol II. Visualization of mNG-EWSR1 in A673 cells treated with PHA-767491 (2 µM) shows a substantial shift of the mNG-EWSR1 signal, particularly foci, from the nucleoplasm to nucleoli (marked by Fibrillarin) at all time points compared to representative DMSO-treated cells **(Figure 7A)**. Analysis of the distribution of mNG-EWSR1, pS2-RNA pol II, and Fibrillarin in TC-32 reporter cells following exposure to PHA-767491 showed comparable results (**Figure S5C**). To quantify the changes in the localization of mNG-EWSR1, we determined colocalization coefficients for mNG-EWSR1 foci and pS5- or pS2-RNA pol II and mNG-EWSR1 and Fibrillarin in control and pS2-RNA pol II treated cells. For these analyses, we calculated MOC colocalization coefficients only because Fibrillarin and mNG-EWSR1’s locations in different sub-nuclear compartments preclude assessment of their intersection under non-treatment conditions. Following treatment with PHA-767491, we observed a substantial decrease in the colocalization of the mNG-EWSR1 foci and pS2-RNA pol II signals and a concomitant increase in the MOC of the mNG-EWSR1foci and Fibrillarin signals (**Figure 7B** and **Figure S5D**). These results are consistent with a selective effect on the location of the endogenous EWSR1 associated with transcriptionally active RNA polymerase II following the inhibition of kinases that regulate DNA replication or transcription.

**Figure 7:**
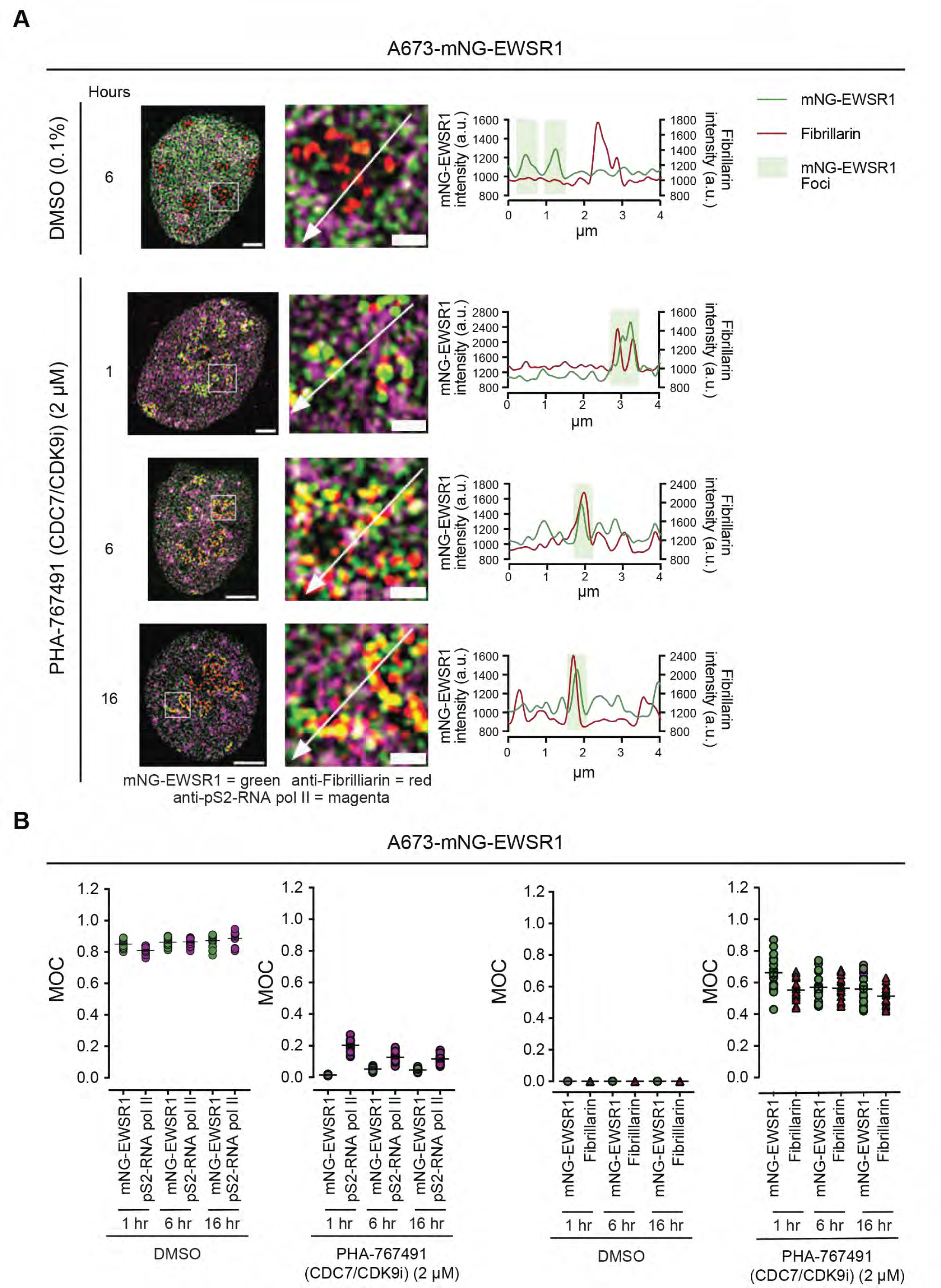
A dual CDC7/CDK9 inhibitor induces changes in the localization of EWSR1 associated with phosphorylated RNA polymerase II. **(A)** Representative merged SIM images and indicated insets showing mNG-EWSR1 (green), pS2-RNA pol II-IF (magenta), and Fibrillarin-IF (red) in A673-mNG-EWSR1 cells treated with either DMSO (0.1%) for 6 hours or PHA-767941 (2 µM) for 1, 6 or 16 hours (whole nucleus scale bar = 3 µm, inset scale bar = 1 µm). The inset images show the lines used to generate the lines plots shown on the right. Representative of 20 images across two independent experiments. (**B**) MOC analysis of mNG-EWSR1 foci and pS2-RNA-pol II and mNG-EWSR1 foci and Fibrillarin following the treatment of modified A673 cells with either DMSO or PHA767941 for 1, 6, or 16 hours.

### FUS localizes in nucleoplasm adjacent to EWSR1

Given the close homology of the FET proteins (**Figure 8A**), we were interested in comparing the distribution of EWSR1 and FUS in our reporter systems. Multiple *in vitro* assays, including immunoprecipitation, two-hybrid analysis, and proximity label-mass spectrometry, have demonstrated the physical association of EWSR1 and FUS (Havugimana et al., 2022; Marcon et al., 2014; Pahlich et al., 2009; Wang et al., 2015). However, we lack evidence for this interaction within cells, and thus we next visualized FUS in our EWS reporter cells. We first assessed the expression of FUS in unmodified A673 and TC-32 cells using immunoblotting and observed that, like EWSR1 (**Figure S2A**), TC-32 cells express slightly more FUS than A673 cells (**Figure 8B**). Next, using IF, we visualized FUS in TC-32 mNG-EWSR1 cells (**Figure 8C**). Unlike mNG-EWSR1, based on DAPI staining, we observed incomplete exclusion of FUS from regions of condensed DNA as well as nucleolar signal. We also observed minimal concurrence in the localization of maximal FUS and mNG-EWSR1 signals suggesting that EWSR1 and FUS exist close to each other but occupy distinct nucleoplasmic spaces, indicating that while homologous, these proteins fulfill related, but different cellular functions.

**Figure 8:**
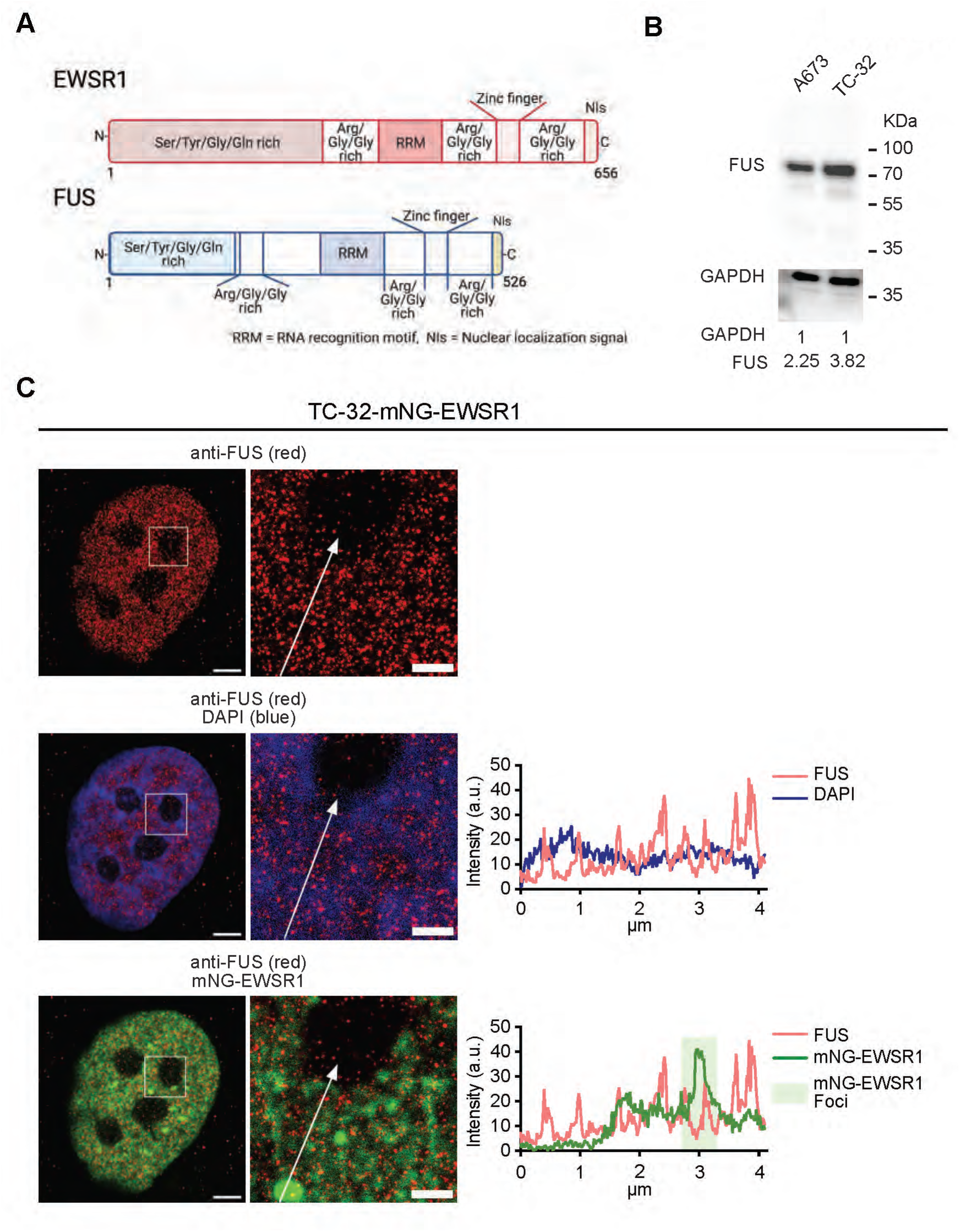
EWSR1 and FUS exhibit close but distinct nucleoplasmic organizations. (**A**) Schematic depicting the similarities between EWSR1 and FUS protein domains. **(B)** Immunoblot of whole cell lysates prepared from unmodified A673 and TC-32 cells and analyzed using antibodies against the indicated proteins. (**C**) Single and dual-color 3D-SIM images of an A673-mNG-EWSR1 nucleus (FUS-IF – magenta, mNG-EWSR1 – green, DAPI – blue; scale bar = 3 µm) and the indicated STED-resolved insets (scale bar = 1 µm), including the line used to the generate the dual channel line profiles shown on the right. In the indicated insets. Representative of 20 images across two independent experiments. **A** generated using BioRender.

## Discussion

In this study, we report that endogenous EWSR1 exists in two visual modalities – distributed and foci, and we define its organization relative to DNA, RNA, and proteins that regulate genome organization and transcription (**Figure 9**). We demonstrate the exclusion of EWSR1 from condensed regions of DNA as indicated by intense DAPI staining, and its presence in euchromatic regions identified by less intense DAPI staining and the accessibility of an antibody against Histone H3. EWSR1 in both visual modalities, distributed and foci, is present in proximity to nascent RNA. The transcriptional regulators, BRD4 and MED1 flank EWSR1 foci. In contrast, EWSR1 foci colocalize significantly with phosphorylated RNA pol II. Our results confirm previous biochemical studies of the FET proteins and extend them to show that within the nucleoplasm, EWSR1’s associations with RNA and RNA pol II are critical determinants of its organization.

**Figure 9:**
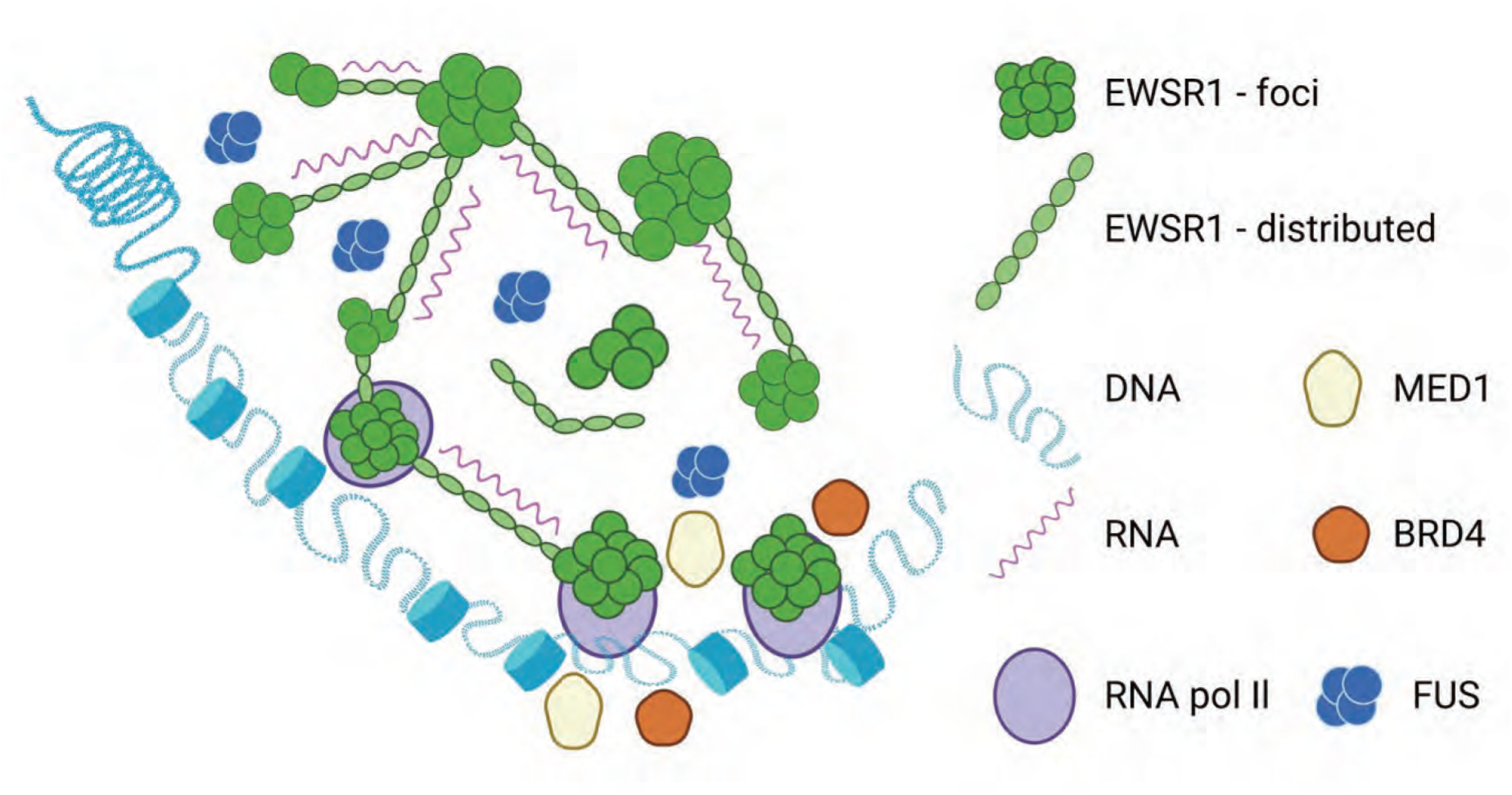
The nucleoplasmic organization of endogenous EWSR1. A schematic model of the nucleoplasmic organization of EWSR1. Figure generated using BioRender.

Previous studies of EWSR1, and FUS, its closely related family member, have shown these proteins contain structured domains that interact with nucleic acids and multiple low complexity domains that increase their binding affinities (Ozdilek et al., 2017) and can mediate concentration-dependent homogenous and heterogenous multivalent protein-protein interactions. Determining the functional consequences of these types of interactions requires quantifying relevant parameters *in vivo* using endogenous protein (Liu and Tjian, 2018; McSwiggen et al., 2019). In this study, we report that endogenous full length EWSR1 exists in two kinetic fractions – one that undergoes rapid exchanges and another that reflects longer-term interactions. These results suggest that depending on the macromolecules with which EWSR1 colocalizes, the kinetics of the interaction may differ significantly and that this has functional consequences. To explore the diversity of EWSR1’s interactions, we used high-resolution visualization of EWSR1’s spatial configuration.

Our study supports the model that EWSR1 binds broadly to pre-mRNA (Schwartz et al., 2013; Wang et al., 2015) as we observe almost complete association of mNG-EWSR1 signals, distributed and foci, with nascent RNA. Examining the mNG-EWSR1 foci in more detail, we show that these colocalize significantly with phosphorylated RNA polymerase, providing visual evidence that within cells endogenous EWSR1 forms part of RNA polymerase assemblies. We speculate that the fluorescent signals used to define the mNG-EWSR1 foci represent an unknown number of individual EWSR1 molecules and that foci could encompass a variable number of EWSR1 molecules. Future studies to address this question will need to assess alternative means of visualizing endogenous EWSR1 that are compatible with higher resolution microscopy as the limitations of resolving the 488 nm fluorescence signals means that we are unable to resolve mNG-EWSR1 beyond 120 nm. Nevertheless, the resolution offered by 3D-SIM imaging revealed that irrespective of signal intensity, fluorescently tagged EWSR1 and EWSR1 imaged using IF shows it exists in a defined nucleoplasmic configuration when compared to that of unfused mNG. Critically, this finding was consistent across the two cell lines used in this study. It is unclear as to which of EWSR1’s interactions are essential to generating this configuration though we hypothesize that it is its significant colocalization with RNA and RNA polymerase II that is principally responsible for the organization we observe; however, its proximity to other nuclear components could also contribute to its arrangement within cells. However, our data suggests that limited penetration of the antibody into EWSR1 foci may not reflect the entirety of this visual modality providing credence for the generation of CRISPR-modified resources to better understand the functions of FET protein family members.

To aid in establishing parameters for the analysis of EWSR1’s localization relative to other nuclear components that may contribute to its organization, in this study, we visualized its colocalization with DNA using the DAPI fluorochrome and Histone H3 and BRD4 and MED1, which other studies have examined as part of the investigation of biomolecular condensates involved in the activation of transcription (Cho et al., 2018; Sabari et al., 2018; Zamudio et al., 2019). Our studies shows that EWSR1 is present only in euchromatic regions of DNA marked by low DAPI staining and anti-Histone H3 IF signals. These results suggest that EWSR1 and Histone H3 exist in proximity to each other, but it will require further study to determine their exact relative organization. We also observed a clear separation of the mNG-EWSR1 foci and the IF signals corresponding to either BRD4 or MED1. However, when interpreting these findings, it is critical to note the limitations of comparing the fluorescence from an endogenously expressed protein and that generated using immunofluorescence. It is also critical to consider that BRD4 and MED1 are much larger proteins than mNG-EWSR1 (>200 kDa versus ∼100 kDa), and MED1 is part of a large multi-component complex. However, the establishment of the mNG-EWSR1 expressing cells used in this study offers the opportunity to modify these lines further to study EWSR1’s colocalization with other endogenous proteins using complementary fluorescent reporter genes.

Differentiating the respective functions of the FUS and EWSR1 proteins has proven challenging because of their homology. Both proteins bind the CTD of RNA polymerase and contribute to the regulation of transcription, but it is unclear whether they engage in the selective regulation of specific transcriptional events or if they can substitute for each other. Interestingly, our visualization of FUS using IF demonstrated that its nucleoplasmic distribution though comparable to that of EWSR1, exhibits a slightly less uniform configuration; however, this observation will need confirmation using endogenously tagged FUS. Critically, and in contrast to proteomic analysis of FUS and EWSR1 (Havugimana et al., 2022; Marcon et al., 2014), we noted minimal colocalization of the two proteins, suggesting that they function separately, and future studies will need to explore these potential differences further.

We also observed the redistribution of EWSR1 foci to the nucleoli of cells following dual inhibition of CDC7/CDK9, consistent with reports of FUS and TAF15 redistribution following exposure to the RNA polymerase inhibitor THZ1 (Yasuhara et al., 2022). Studies assessing FET protein localization under different biological conditions are critical as alterations in their nuclear-cytoplasmic distribution is a pathological feature of diseases associated with mutations in *FET* genes (Harrison and Shorter, 2017). Mutations in FET genes, particularly *FUS*, are associated with cases of amyotrophic lateral sclerosis (ALS) or frontotemporal dementia (FTD) (Da Cruz and Cleveland, 2011; Mackenzie et al., 2010). A critical pathological characteristic of these mutant FET proteins is the disruption of their nuclear localization and the formation of cytoplasmic aggregates (Mackenzie et al, 2010, Kwiatkowski et al 2009, Vance et al 2009, Da Cruz 2011).

While outside the focus of our study, we confirm that the modified cell lines used in this study should serve as appropriate models for the visualization of endogenous EWSR1 at high resolution in the context of stimuli that disrupt FET protein localization and the investigation of the mechanisms that regulate these responses.

In this study, we selected to use two EWS cell lines to generate our model systems for the analysis of EWSR1’s cellular localization because these cells are uni-allelic for *EWSR1*, harbor few other mutations and have stable genomes (Brohl et al., 2014; Crompton et al., 2014; Tirode et al., 2014). There is also minimal evidence for the disruption of EWSR1’s function in tumors, though studies have suggested it may contribute to R-loop accumulation in EWS cells (Gorthi et al., 2018) or more recently in maintaining centromere identity (Kitagawa et al., 2023). Future experimentation employing the modified cell lines described in this study have the potential to aid investigation of the response of EWS cells to external stimuli such as DNA damage, the effect of altering the post-translational modifications of EWSR1, which studies have linked to its propensity to aggregate (Nosella et al., 2021), and the opportunity to study EWSR1’s interaction with the EWSR1::FLI1 fusion oncoprotein (Embree et al., 2009; Spahn et al., 2003) in further depth. Nevertheless, our findings using the EWS cell lines suggest these are appropriate models for the study of EWSR1, as we have validated previously described *in vitro* findings and the results across the two cell lines proved highly reproducible. It will, however, be essential for future studies to confirm our findings using additional cell line model systems, and our study offers validated resources, including the reagents needed for gene targeting, to accelerate such efforts.

---

**Figure S1:**
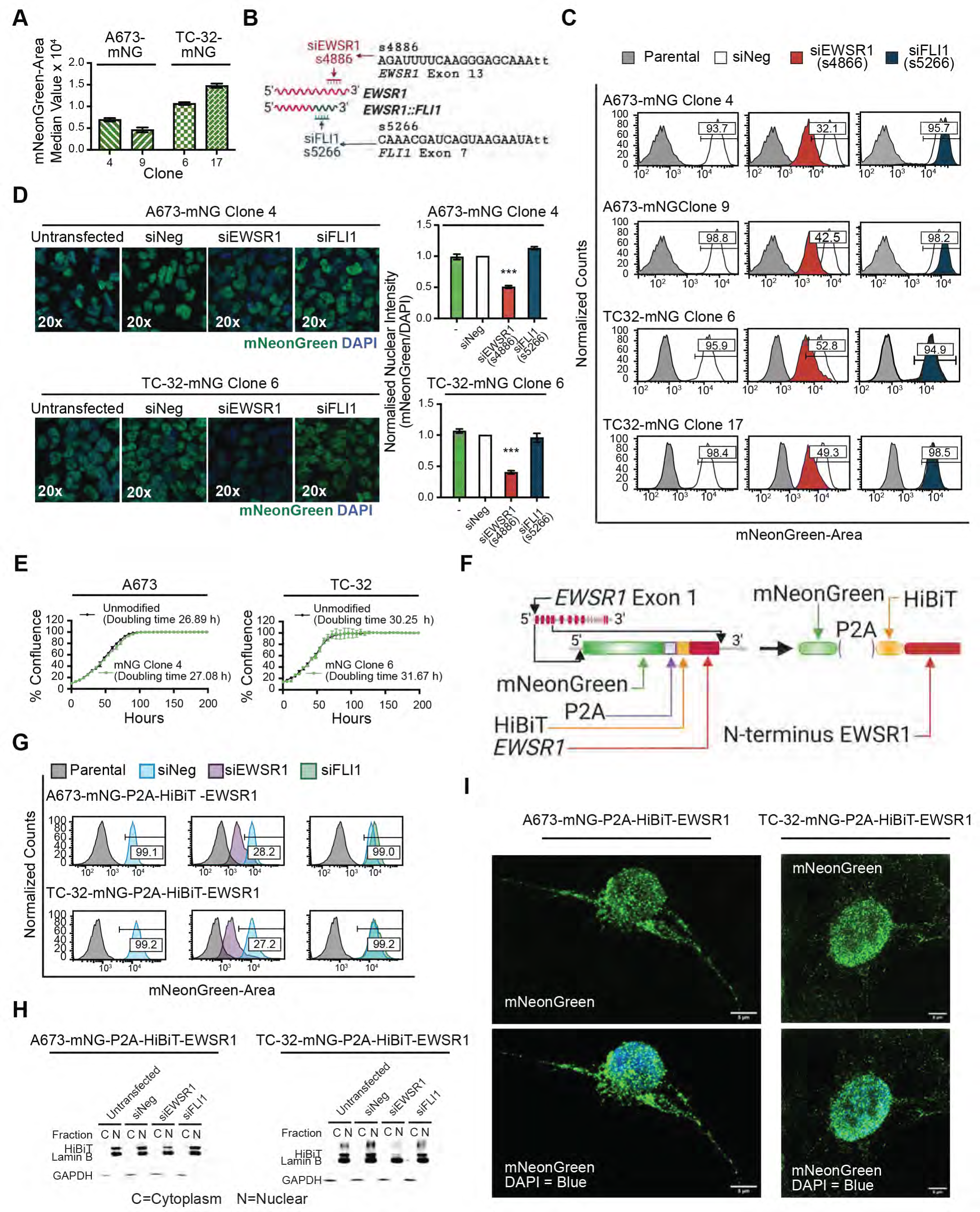
The generation of EWS cell lines expressing mNeonGreen-EWSR1 and isogenic control cell lines expressing unfused mNeonGreen. (A) Median fluorescent value for two A673-mNG EWSR1 clones and two TC-32 mNG-EWSR1 clones (three independent analyses of each clone; mean ± SEM). (B) Schematic depicting the siRNAs targeting *EWSR*1 or *EWSR1::FLI1* (see Grohar et al., 2016 for validation) used to characterize modified clones. (C) Representative flow cytometry analysis of A673-mNG-EWSR1 and TC-32-mNG-EWSR1 modified clones. The shift in mNG fluorescence following transfection of the indicated siRNAs (48 hours for TC32, 72 hours for A673) suggests insertion of the reporter gene into the non-rearranged *EWSR1* allele (middle panels) for the representative clones shown here. Representative data of three independent experiments. (D) Images of mNG fluorescence 48 hours post-transfection of the indicated siRNAs in the A673-mNG and TC-32-mNG clones and quantification of the mNG signal normalized to DAPI (n=9 nuclei per transfection; mean ± SEM). (E) Proliferation curves (cell counts normalized to their respective day 0 counts) of A673-mNG clone 4 (n=3) and TC-32-mNG clone 6 (n=3) compared to respective unmodified A673 (n=3) and TC-32 parental cells (n=3). (F) Schematics of the insertion of the indicated DNA elements into the *EWSR1* genomic locus of EWS cells and expression of the cleaved mNG protein and one part of a split luciferase (HiBiT) fused to the N-terminus of EWSR1. (G) Representative flow cytometry analysis of A673 and TC-32 modified clones. The shift in mNG fluorescence following transfection of the indicated siRNAs suggests the insertion of the reporter gene into the non-rearranged *EWSR1* allele. (H) Immunoblots of fractionated protein lysates prepared from mNG-EWSR1 modified A673 or TC-32 cells transfected with indicated siRNAs and analyzed using the indicated antibodies. C=cytoplasm; N=Nuclear. (I) Representative super-resolution confocal microscopy images of modified A673 and TC-32 cells expressing unfused mNG. Scale bar = 6 µm. Representative of 30 images across three independent experiments. B and I generated using BioRender.

**Figure S2:**
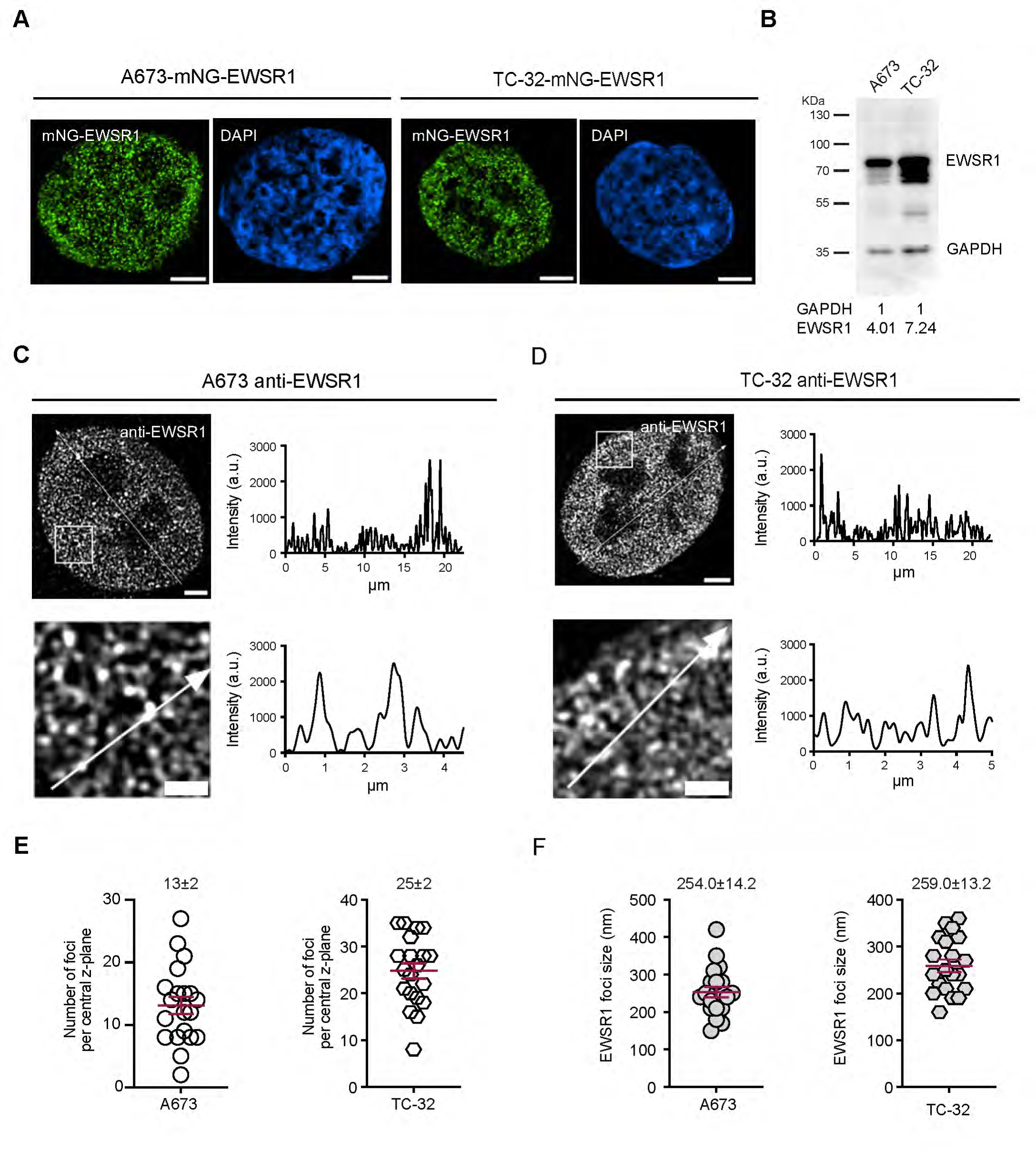
Structured illumination microscopy of endogenous EWSR1 using immunofluorescence. (**A**) Representative, deconvolved images of an A673-mNG-EWSR1 nucleus and a TC-32-mNG-EWSR1 nucleus (mNG-EWSR1, green; DAPI, blue (∼35 z-steps of 0.15 µm covering an average of 5 µm nuclear thickness). Scale bar = 3 µm. Representative of >100 images across >15 independent experiments. (**B**) Immunoblot of whole cell lysates prepared from unmodified A673 and TC-32 cells and analyzed using antibodies against the indicated proteins. Indicated are the relative intensities normalized to loading controls. (**C**) A 3D-SIM image of an A673 nucleus (scale bar = 3 µm) and the indicated inset (scale bar = 1 µm) showing immunofluorescent detection of EWSR1. Each images shows the indicated lines used to generate the corresponding intensity plot. Representative of 20 images across three independent experiments. (**D**) A 3D-SIM image of a TC-32 nucleus (scale bar = 3 µm) and the indicated inset (scale bar = 1 µm) showing immunofluorescent detection of EWSR1. Each images shows the indicated lines used to generate the corresponding intensity plot. Representative of 20 images across three independent experiments. (**E**) Number of EWSR1 foci per central z-plane of an A673 or TC-32 nucleus (n=20) in (applied thresholds for EWSR1 foci = 3X greater intensity and size restriction of 20nm). **(F)** EWSR1 foci size (n=20) in A673 and TC-32 cells. The red lines in D and E indicate the mean ± SEM.

**Figure S3:**
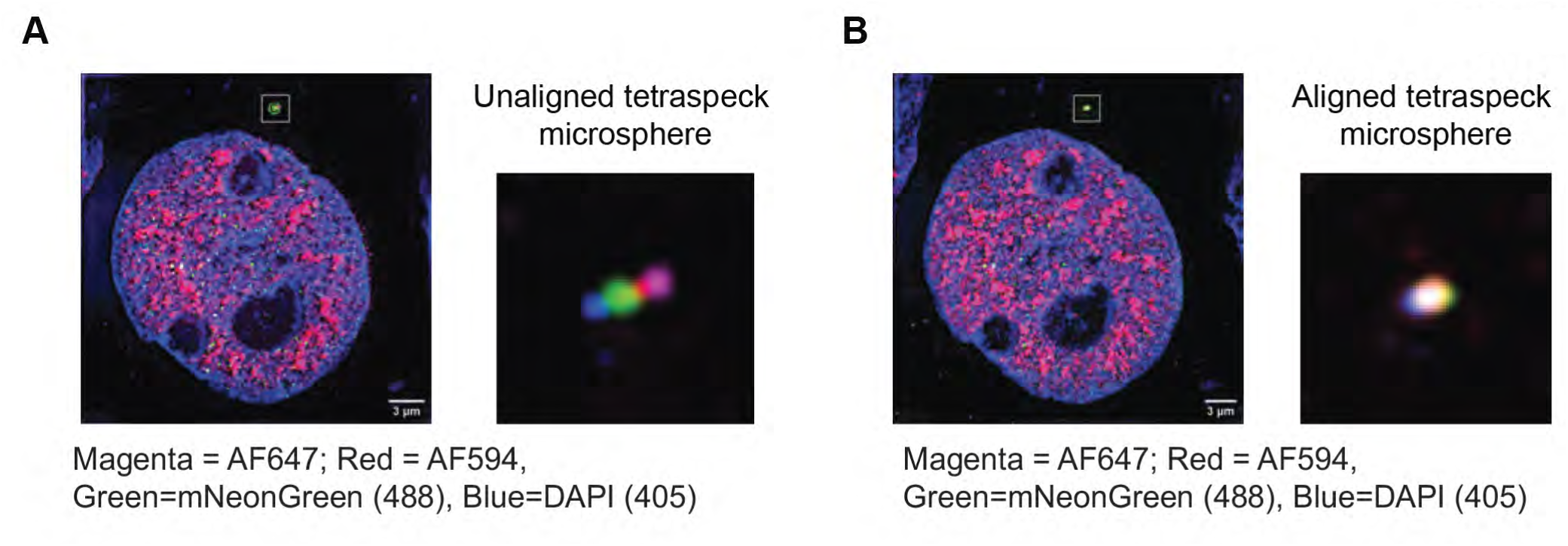
Laser alignments during image acquisition. **(A)** Representative nucleus imaged in 405, 488, 594 and 647 channels using 3D-SIM showing a marked TetraSpeck microsphere in the same plane of view. Inset shows enlarged merged view of the unaligned microsphere. **(B)** Same representative nucleus shown in (A) using 3D-SIM showing a marked TetraSpeck microsphere in the same plane of view. Inset shows enlarged merged view of the now aligned microsphere.

**Figure S4:**
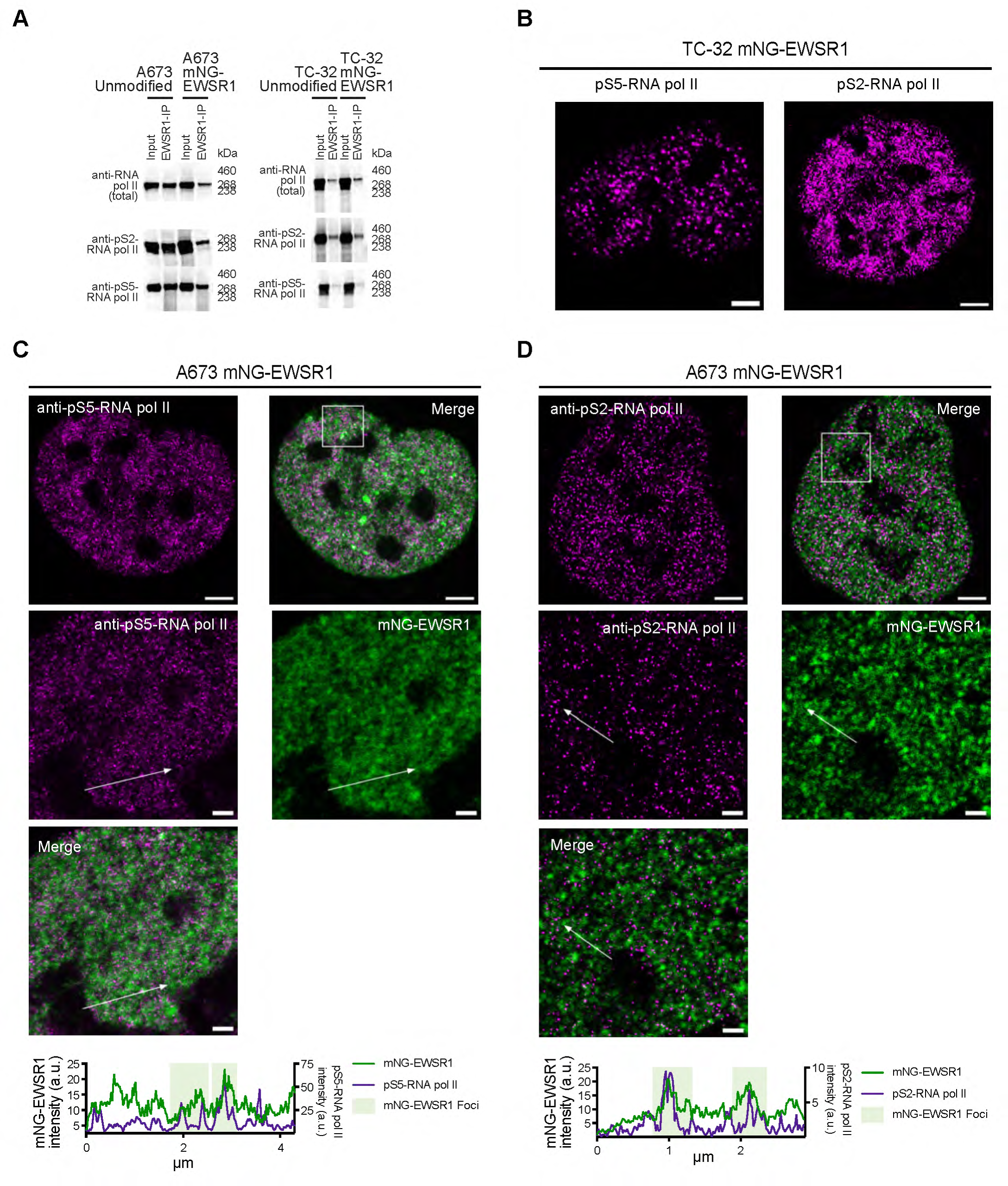
EWSR1 concentrates at regions of active transcription. **(A)** EWSR1-immunoprecipitation (IP) of protein lysates prepared from unmodified (A673 and TC-32) and modified (A673-mNG-EWSR1 and TC-32-mNG-EWSR1) cells and immunoblotted using antibodies against the indicated antibodies. **(B)** Single-channel pS5-RNA pol II and pS25-RNA pol II images of the TC-32-mNG-EWSR1 nucleus presented in Figures 7E and **7F** (scale bar = 3 µm). Representative of 20 nuclei across two independent experiments. (**C**) Merged 3D-SIM image of an A673-mNG-EWSR1 nucleus (pS5-RNA pol II-IF – magenta, mNG-EWSR1 – green) (scale bar = 3 µm) and the indicated STED-resolved insets (scale bar = 1 µm), including the line used to generate the dual channel line profile shown below. Representative of 20 nuclei across two independent replicates. **(D)** Merged 3D-SIM image of an A673-mNG-EWSR1 nucleus (pS2-RNA pol II-IF – magenta, mNG-EWSR1 – green) (scale bar = 3 µm) and the indicated STED-resolved insets (scale bar = 1 µm), including the line used to generate the dual channel line profile shown below. Representative of 20 nuclei across two independent replicates.

**Figure S5:**
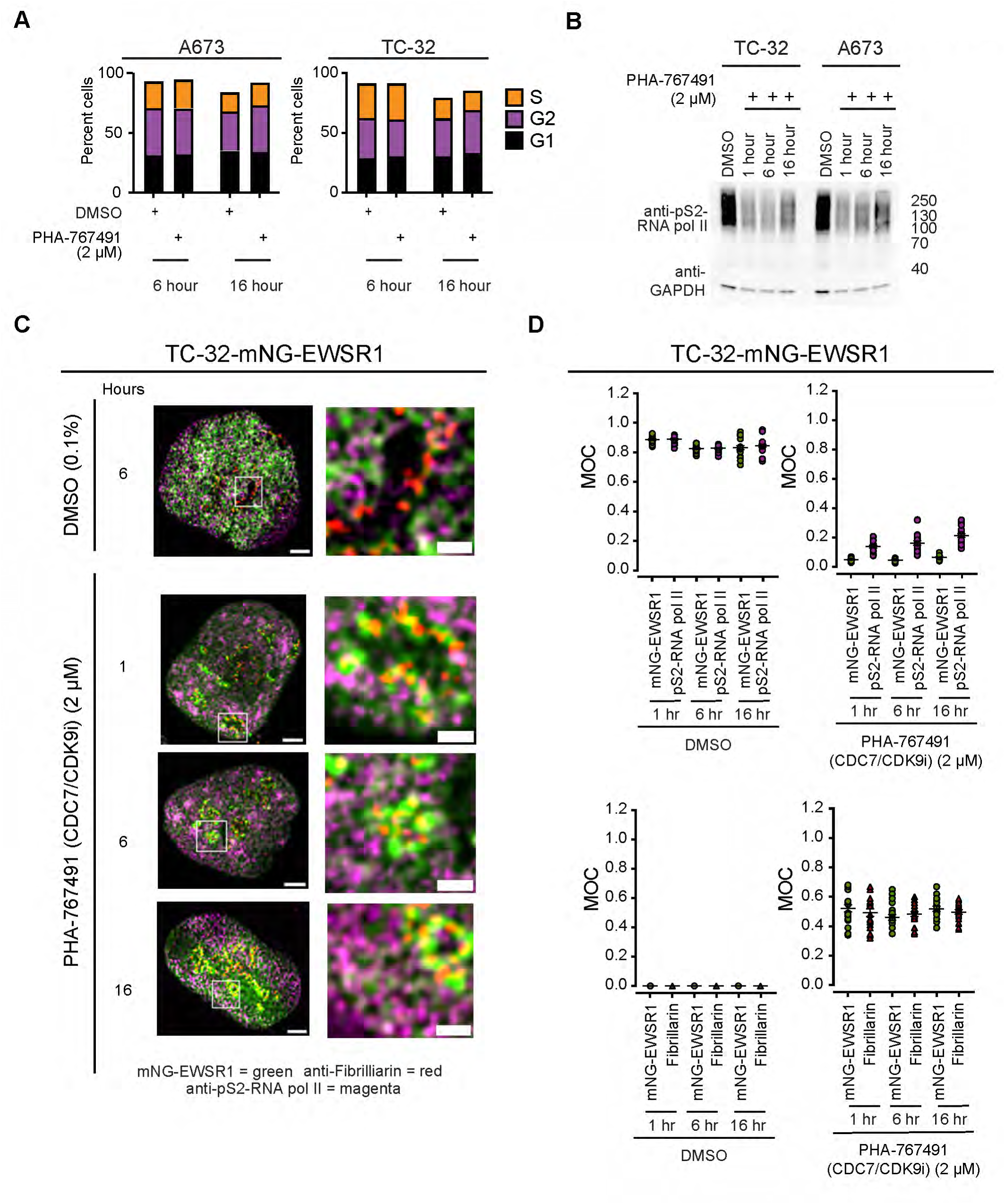
A dual CDC7/CDK9 inhibitor induces changes in the localization of EWSR1 associated with RNA pol II. **(A)** Cell cycle analysis of (DMSO) or PHA767941-treated A673 or TC-32 cells. (**B**) Immunoblot of whole cell lysates prepared from control (DMSO) or PHA767941-treated A673 or TC-32 cells using antibodies against the indicated proteins. (**C**) Representative merged images obtained using 3D-SIM showing mNG-EWSR1, pS2-RNA pol II, and Fibrillarin in TC-32-mNG-EWSR1 cells treated with either DMSO (0.1%) for 6 hours or PHA-767941 (2 µM) for 1, 6 or 16 hours (Scale bar = 3 µm). Representative of 20 images across two independent experiments with the indicated insets shown (Scale bar = 1 µm). **(D)** MOC analysis of mNG-EWSR1 foci and pS2-RNA-pol II and mNG-EWSR1 foci and Fibrillarin following the treatment of modified TC-32 cells following treatment with either DMSO or PHA767941 for 1, 6, or 16 hours.

**Table S1.**
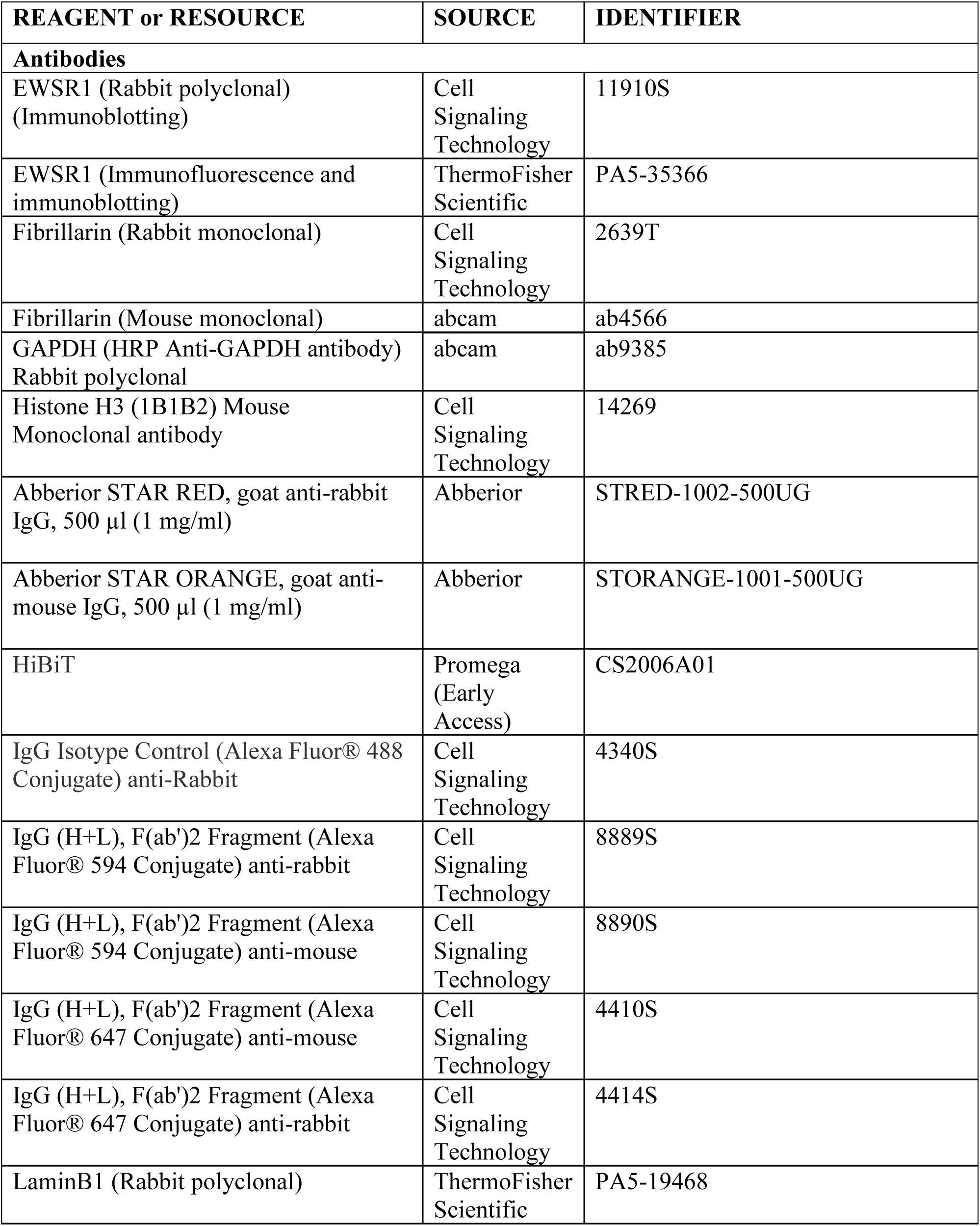
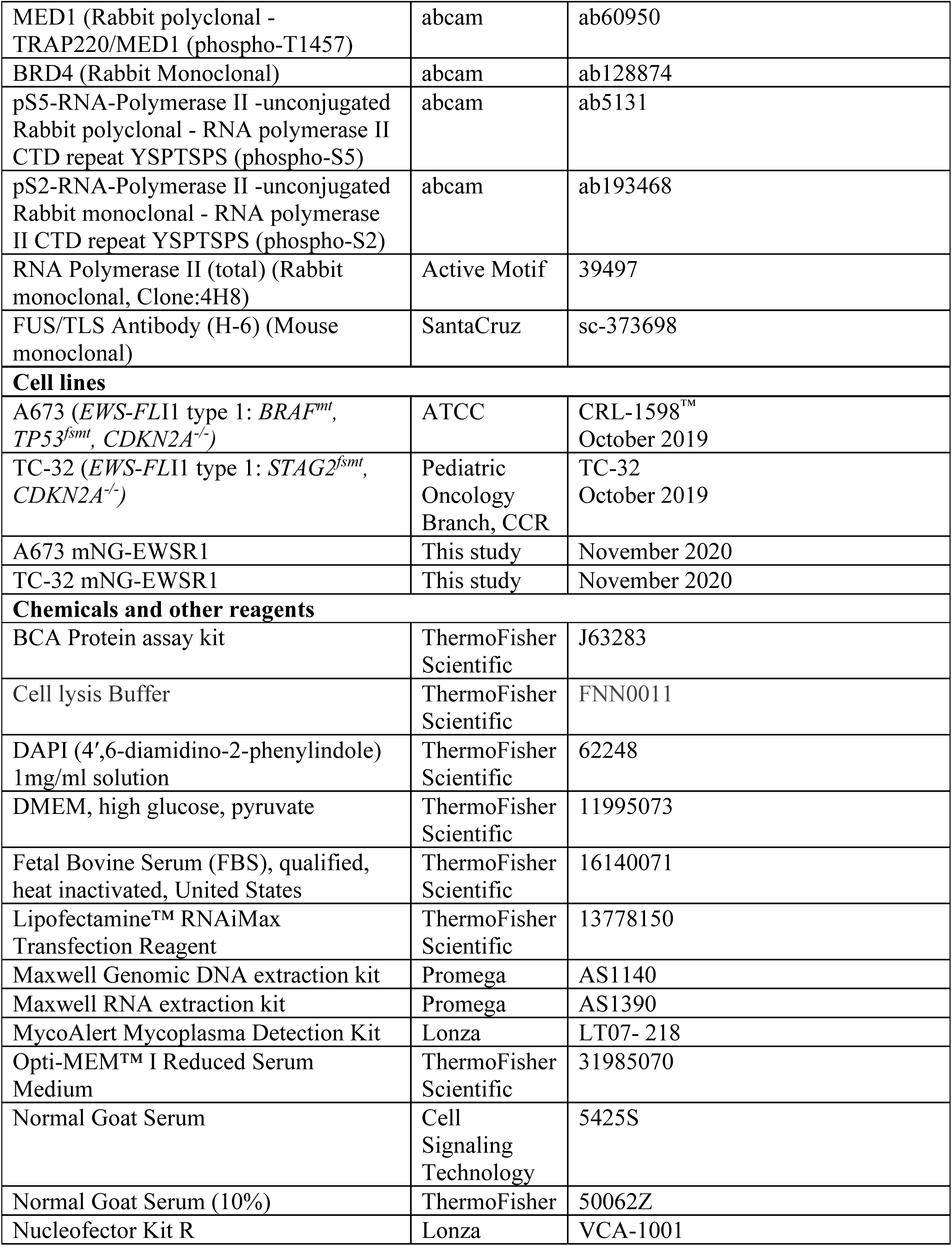
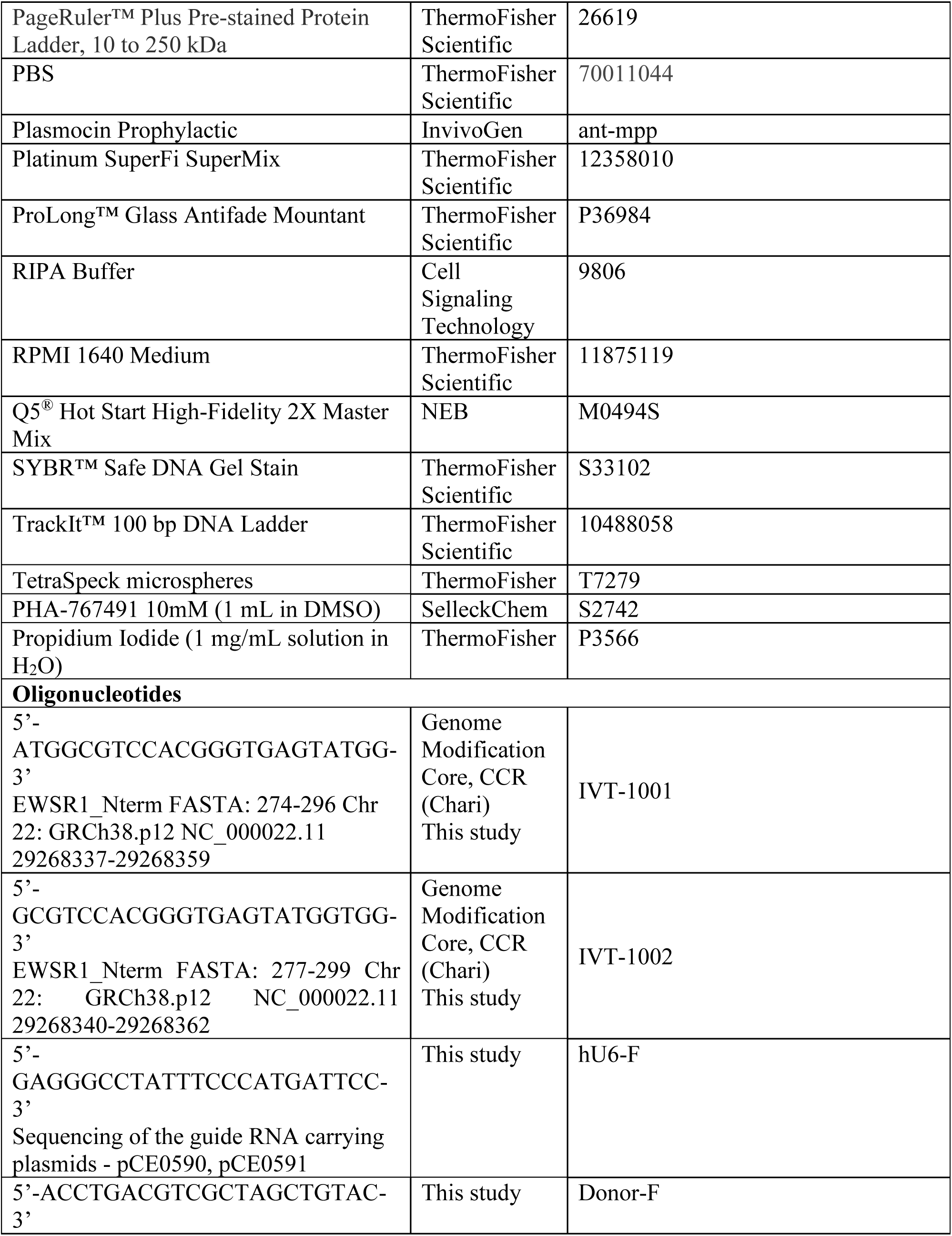
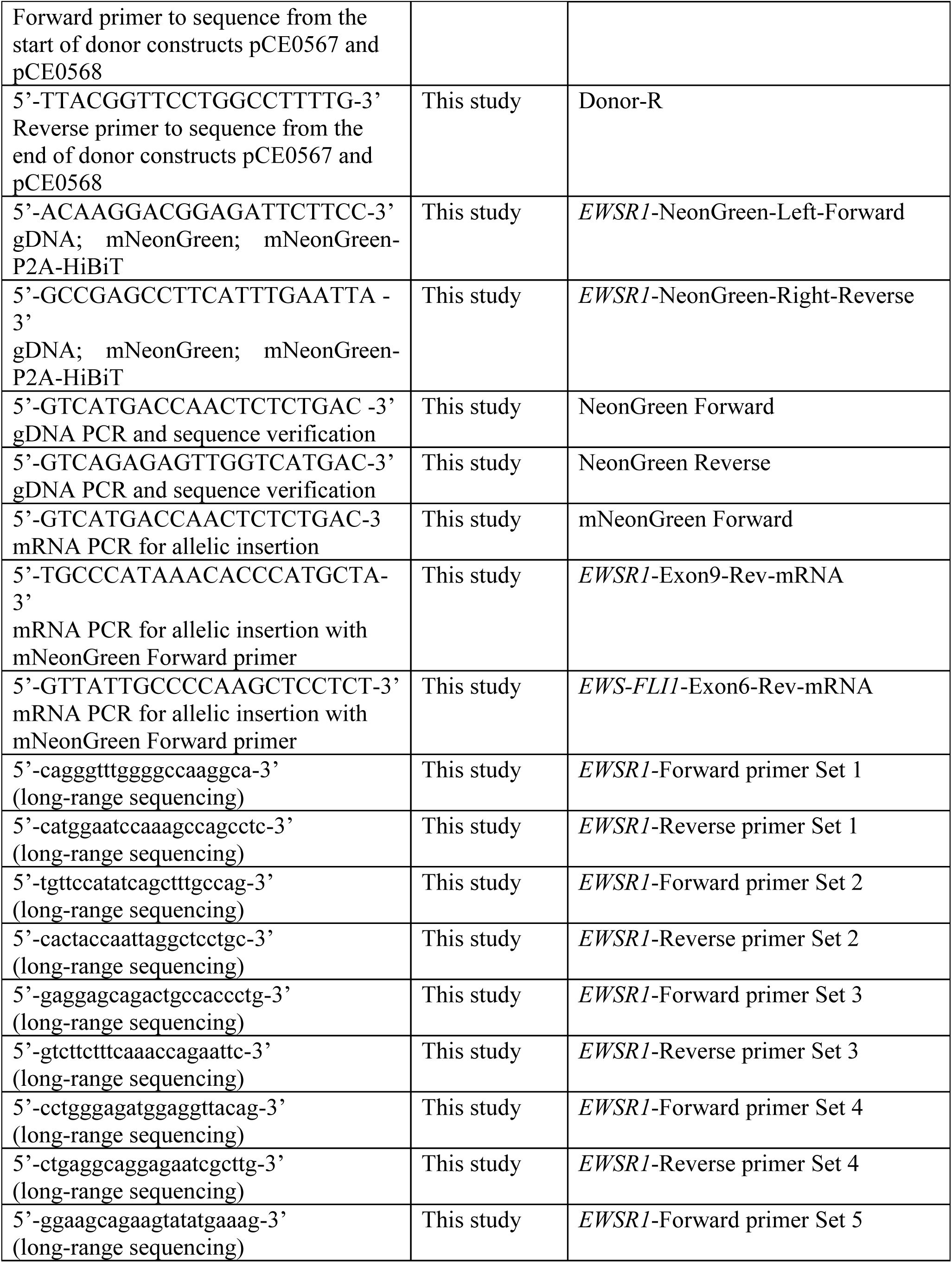
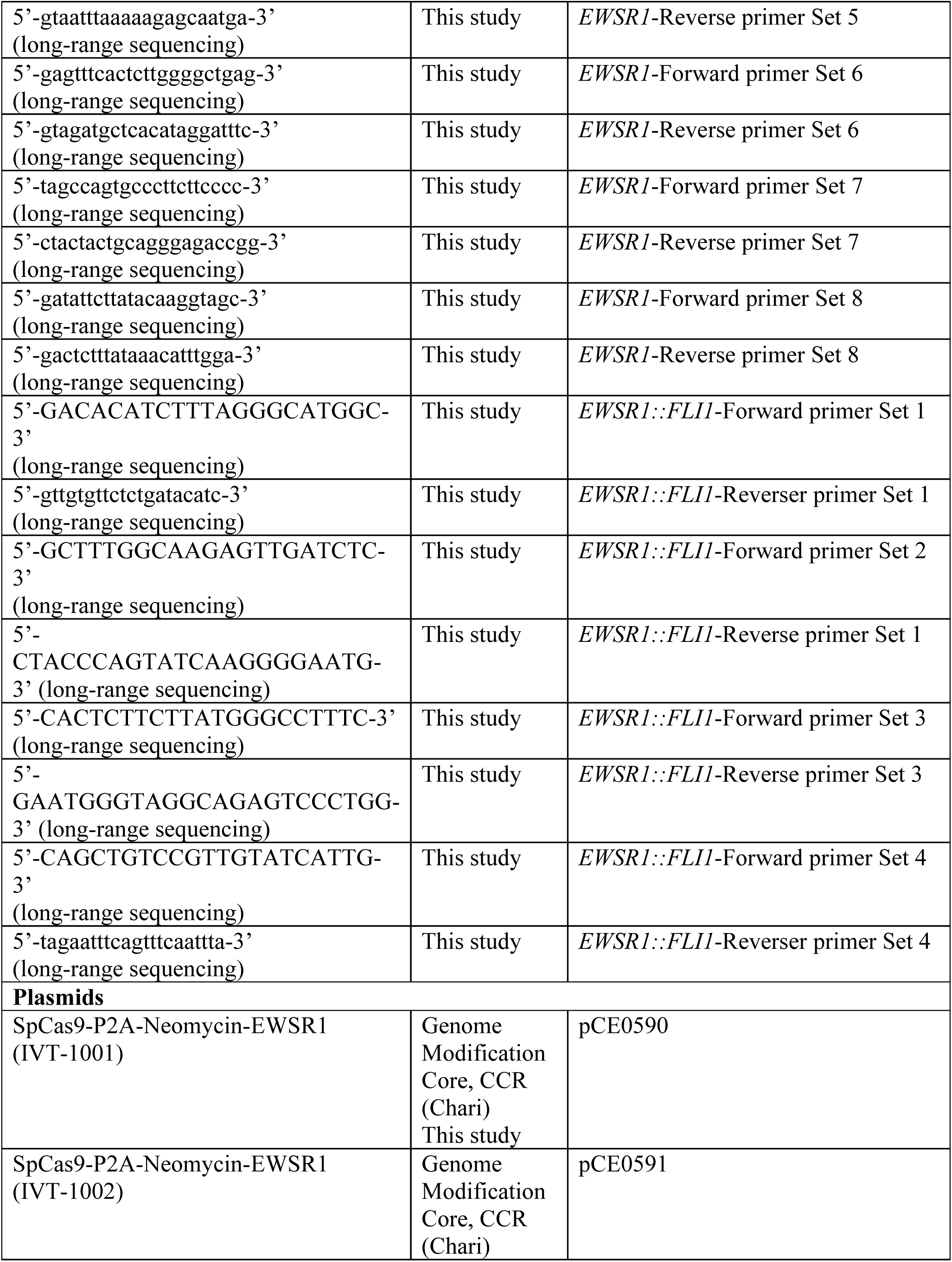
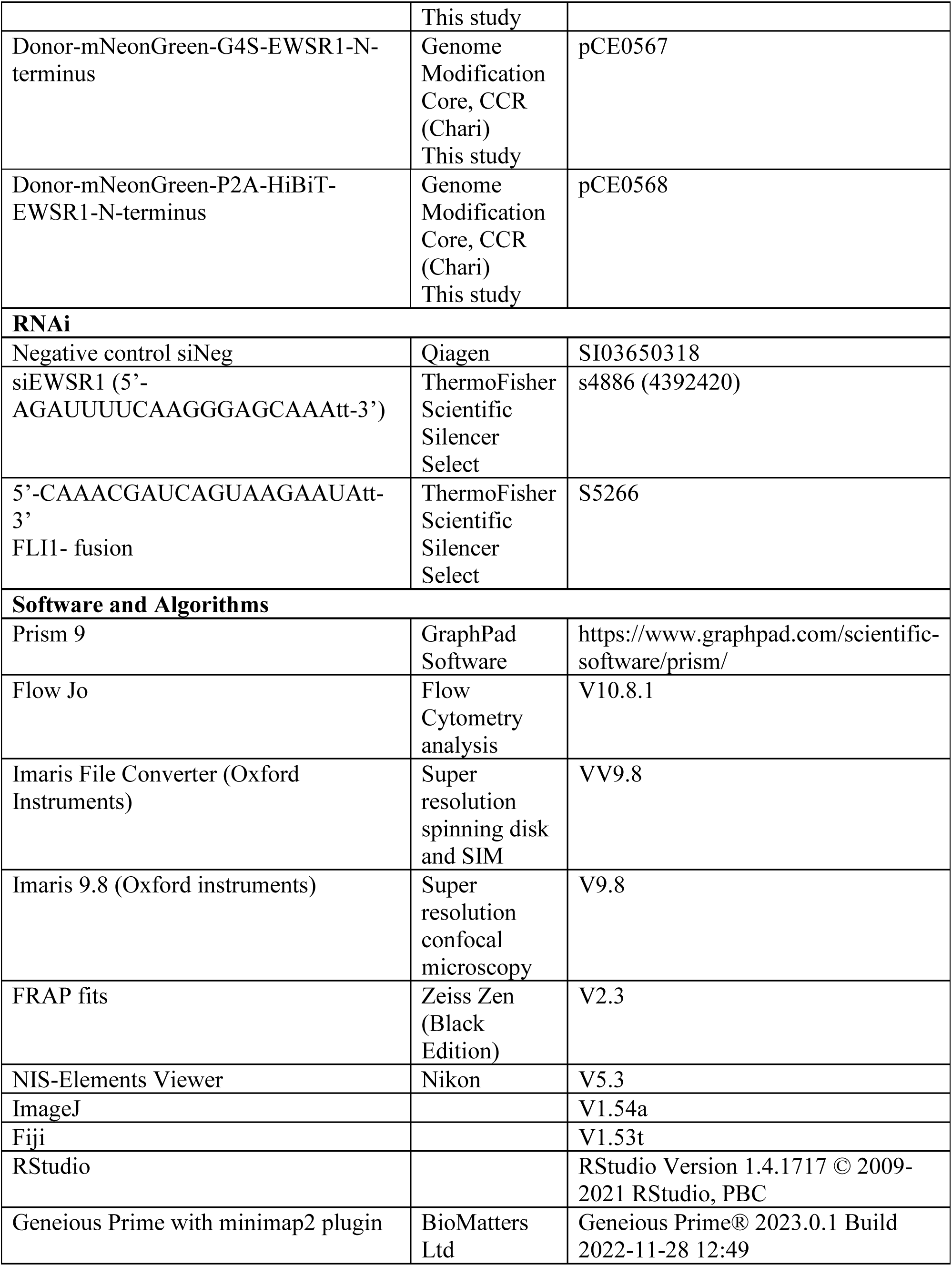
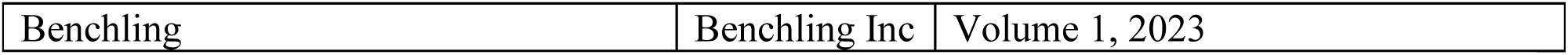
Reagents and resources.

## Contact for Reagent and Resource Sharing

Further information and requests for reagents should be directed to lead contact Natasha Caplen.

## Experimental Model and Subject Details

A673 (ATCC, Manassas, Virginia) cells were cultured in DMEM media, and TC-32 cells (a gift from the Pediatric Oncology Branch, CCR) in RPM1 1640 media. Media was supplemented with 10% FBS (ThermoFisher Scientific) and Plasmocin Prophylactic (Invivogen, San Diego, CA), and cells were grown at 37°C, 5% CO_2_. We confirmed the identity of cell lines using short tandem repeat (STR) analysis (ATCC) at intervals throughout experimentation, and we monitored for mycoplasma contamination using the MycoAlert Plus system (Lonza, Walkersville, MD). **Table S1** summarizes relevant mutation information for each cell line, and **Supplementary File 2** reports the most recent STR fingerprint information for parental and modified cell lines.

## Method Details

### Plasmid construction and the generation of CRISPR-Cas9-modified EWS cells

To select single guide RNA (sgRNA) sequences, we used sgRNAScorer2.0 (Chari et al., 2017) and we selected sgRNAs IVT-1001 and IVT-1002 (**Table S1**) for the generation of the plasmids pCE0590 (Lenti-SpCas9-2A-mCherry-*EWSR1*-IVT-1001) and pCE0591 (Lenti-SpCas9-2A-mCherry-*EWSR1*-IVT-1002), respectively (**Table S1)**. We used 800 bp around the predicted cut site as homology arm templates that included minor sequence changes at the 5’ end of *EWSR1* exon 1 within the right homology arm (noted in **Figure 1A**). We inserted into EWS cell lines a DNA cassette consisting of the mNeonGreen (mNG) fluorescent reporter protein and a 45-nucleotide sequence encoding a G4S3 linker (GGGGS x 3) to form a flexible linker adjacent to the N-terminal of the tagged protein (pCE0567: NeonGreen-Linker-N-term-*EWSR1*). We included this sequence to mitigate the potential effects of the reporter on the structure, and, thus, the function of the modified protein. We electroporated EWS cells using the Lonza kit T and the Lonza kit R reagents and 2 µg of donor plasmid and 0.5 µg of each sgRNA plasmid using the following conditions: TC32: Program D-032 and 0.5x 10^6 cells; A673: program A-028, 1 x 10^6 cells. Following electroporation, we grew cells for several days and used mNG fluorescence and flow cytometry to sort single cells. We maintained clones for three to four weeks until we generated enough cells for confirmation of the inserted DNA. We used the same strategy to insert into EWS cell lines a DNA cassette consisting of the mNG fluorescent reporter, a cleavable P2A peptide, and the HiBiT peptide (part of split luciferase reporter) to generate cells expressing unfused mNG and a fusion of the HiBiT peptide (**Figure S1H**) and EWSR1 (pCE0568: NeonGreen-P2A-HiBiT-*EWSR1*). **Supplementary File 3** details the maps of each plasmid.

### Flow cytometry

For flow cytometry, we pelleted cells by centrifugation and washed each sample once with ice-cold PBS. Unstained control samples were resuspended in FACS buffer (Phenol-red free media with 2% FBS) and other samples were incubated with DAPI live-dead stain for ten minutes before being pelleted and resuspended in FACS buffer. All samples were passed through a cell strainer to obtain a single-cell suspension. At least 10,000 events were assessed per sample on a SONY4800 spectral flow analyzer and gated for single cells using side scatter and forward scatter. mNeonGreen and DAPI fluorescence was set using single-stained controls, and mNeonGreen positive percentages were gated on single cells that were DAPI negative. The output from the Sony4800 analyzer was obtained as .FCS files and replotted in FlowJo V10.8.1 to generate the histograms shown in this study. For cell cycle analysis, 150,000 cells were plated and treated with DMSO or 2 µM PHA-767941. Cells were fixed at the appropriate time points post-treatment using 70% ice-cold ethanol for at least 2 hours at 4°C. Cells were treated with 50 µg/ml of Ribonuclease A and stained with 40 µg/ml propidium iodide for 30 minutes. Samples were run on a spectral analyzer SONYID7000 and 20,000 gated events were collected per sample. Data was imported into FlowJo and the cell cycle module was used for analysis. Obtained percentages were inputted into GraphPad and plotted.

### RNAi

**Table S1** lists the siRNAs used in this study, and **Figure S1B** depicts the relative locations of the sequences targeted within either the *EWSR1* or *EWSR1::FLI1* transcripts. Cells were transfected using 20 nM siRNA complexed with RNAi-Max (ThermoFisher Scientific) at 4.5 µl/well using 135,000 cells per well of a 6-well plate. Cells were harvested 48 hours post-siRNA transfection unless otherwise noted. For confirmation of the specificity of the siRNAs targeting *EWSR1* and *EWSR1::FLI1,* see (Grohar et al., 2016; Neckles et al., 2019; Vo et al., 2022).

### Proliferation assay

Cells (10,000) of either parental or mNG-modified cells were plated per well in a 6-well plate in triplicate and placed in an Incucyte system (Sartorius, Essen BioScience, Inc. Ann Arbor, MI). Four images per well (total =12 per sample) were taken every 6 hours for 7 days. Raw data from the Incucyte system was obtained and analyzed using the Incucyte 2020B GUI Analysis Software and the analyzed data (mean and error) per well was exported to an Excel file and transferred into GraphPad Prism to plot the results shown.

### Genomic DNA analysis

Primers used for sequencing are listed in **Table S1**. For PCR products 600 bp and less than 1000 bp, Phusion polymerase (NEB, Ipswich, MA) was used. For PCR products greater than 1000 bp, Q5 Hot-Start (NEB, Ipswich, MA) master-mix was used. PCR products were first run on a 1% agarose gel to ensure amplification of the expected product. The amplified product was then gel extracted and PCR purified prior to sequencing. To confirm the insertion of each tag into the 5’ end of *EWSR1* exon 1, we first used PCR to amplify across either junction of the insertion or the entire left and right homology arms into the insertion sites and subjected each amplified product to Sanger sequencing. To further validate the correct insertion of the mNG reporter gene cassette and associated sequences, a total of 14 primer pairs were designed to amplify the entire *EWSR1* genes in parental and mNG-modified cells. Each primer pair was designed to produce a 5 kb product, and all primer pairs overlapped to ensure complete sequence coverage. PCR conditions were optimized, using the Q5 Hot Start 2x master-mix for the amplification of a single 5 kb product per primer pair. We obtained single amplified products for12 out of the 14 PCR primer pairs and these amplified products were purified using the QIAquick PCR Purification Kit (28104, Qiagen, Germantown, MD) before sequencing. For the remaining two PCR reactions, we subjected the amplified products to gel electrophoresis and DNA at the expected product size was excised from the gel and extracted using the QIAquick Gel Extraction Kit (28706X4 Qiagen). Purified DNA was subjected to long-read sequencing (PacBio) and post-sequencing .ccs files for each sample were obtained and assembled for comparison to reference sequences. Reference sequences were built in Benchling using the *EWSR1* (ENSG0000018294) genomic sequence and *de-novo* sequences for mNG-EWSR1 were generated. Minimap2 was used to annotate and align the output from PacBio sequencing to these reference sequences using default trimming and mismatch calling parameters.

### Immunoprecipitation

Cells were grown to 80% confluence prior to IP. Cells were harvested, washed with ice-cold PBS, and, in RIPA lysis buffer (10mM Tris-HCl, pH 8.0, 1mM EDTA, 0.5mM EGTA, 1% Triton X-100, 0.1% Sodium Deoxycholate, 0.1% SDS, 140mM NaCl) supplemented with protease/phosphatase inhibitor cocktail (CST #5872S). For TC-32 cells, 2 µg, and for A673 cells, 4 µg of EWSR1 antibody were conjugated in 50 µl of magnetic beads. Protein lysates were quantified and resuspended at a concentration of 2 mg/ml in RIPA per immunoprecipitation. Resuspended lysates were incubated with antibody-coupled magnetic beads at 4°C overnight with gentle rotation, washed and eluted, and samples were examined by immunoblot analysis along with inputs (20 µg and 10 µg for TC32 and A673, respectively).

### Immunoblotting

Whole-cell lysates were prepared using cell extraction lysis buffer (ThermoFisher, Waltham, MA. Catalog: FNN0011) and sonication. Protein concentrations were calculated using a standard BCA assay (ThermoFisher, Waltham, MA, Catalog number: J63283.QA), and 15 µg of each protein sample was analyzed following standard immunoblotting methods using the antibodies listed in **Table S1** and detection using ECL. For the analysis of mNG-P2A-HiBiT-EWSR1 A673 and TC-32 cells, lysates were collected and suspended in ice in hypotonic buffer (20 mM Tris-HCl, pH 7.4, 10 mM NaCl, 3 mM MgCl_2_) for 15 minutes. Following centrifugation at 5000 rpm for 10 minutes, supernatants were collected as the cytoplasmic fractions. The remaining pellets were resuspended in cell extraction lysis buffer and sonicated. The sonicated samples were centrifuged at 10,000 rpm for 10 minutes, and supernatants were collected as the nuclear fraction. The protein concentrations of the cytoplasmic and nuclei fractions were quantified using a BCA assay, and 5 µg of each fraction was assayed using standard immunoblotting methods using the anti-HiBiT and loading control antibodies listed in **Table S1**. For quantification of relative intensity, the same region of interest (ROI) was drawn across all protein signals from gray-scale raw images of the immunoblots, along with five measurements of background. Mean intensity measurements were exported, and values were inverted. The mean background was subtracted, and intensity relative to the appropriate loading control protein signal was calculated.

### Confocal microscopy

Cells (50,000) were plated on coverslips and allowed to attach overnight. Coverslips were washed twice with PBS and fixed with 3.7% formaldehyde solution for 5 minutes at room temperature (RT). After three washes with PBS, cells were permeabilized with a 0.01% Triton-X-100 solution at RT for 5 minutes. Coverslips were washed three times in PBS and allowed to block for 2 hours at RT in blocking solution (1X PBS / 5% normal serum / 0.3% Triton™ X-100). Coverslips were washed thrice with PBS and mounted onto labeled slides with VECTASHIELD® Antifade Mounting Medium with DAPI (H-1200-10, Newark, CA) and allowed to dry. Coverslips were sealed with clear nail polish. Images were captured on the Zeiss LSM780 Inverted microscope at the indicated magnifications. All images were obtained as. czi files from the scope. Unmodified parental cells were used to obtain background values for signal in the 488 channel which was subtracted from the samples. DAPI was used as a nuclear mask and applied to all samples post background subtraction. Integrated density was calculated on the masked ROIs using the function in ImageJ. Average integrated density across samples were plotted. The average signal density with SEM was plotted as shown across the indicated number of replicates.

### Fluorescence recovery after photobleaching

Cells (50,000) were plated in a 35 mm glass bottom dish (Ibidi USA Inc. Fitchburg, Wisconsin, Catalog No: 81218-200). Before imaging, the media was changed to either phenol-red free RPMI supplemented with 10% FBS for TC-32 modified cells or Fluorobrite DMEM media supplemented with 10% FBS for A673 modified cells. Regions 3 µm in diameter were drawn for reference, background, and ROI (region of interest). 488nm laser intensity was set to 72%, and cells were imaged every second for 3 minutes using the Zeiss LSM880 Airyscan microscope. Signal intensity was subtracted from mean background intensity and normalized to the background subtracted reference signal intensity and plotted over time. A double exponential curve was used to fit the data for mNG-EWSR1 cells, and a single exponential curve was used to fit the data for the unfused mNG control. Unfused mNG cells were generated by using CRISPR-Cas9 based editing to introduce a mNeonGreen-P2A-HiBiT tag at the 5’ end of EWSR1 exon 1 as for the generation of mNG-EWSR1 reporter cells. The presence of the high efficiency P2A cleavage peptide cleaves off the mNG protein post translation resulting in cells that express unfused mNG and HiBiT tagged EWSR1. **Supplementary File 3** details the maps of each plasmid. Dissociation constants were calculated after normalizing data to the control sample. All statistical measures and curve fitting were conducted in the Zeiss Zen (black edition) software.

### Super resolution microscopy, 3D-structured illumination microscopy (SIM) and 2DSTED microscopy

Cells (130,000) were either plated or used to set up transfections as described earlier up and plated in wells containing coverslips. At the indicated time point, cells were washed twice with PBS and fixed using 3.7% formaldehyde solution, prepared fresh. Cells were permeabilized using PBS with 0.5% Triton X-100 for 5 minutes. After three washes with PBS, cells were incubated for two hours at RT in blocking solution (1X PBS, 5% Normal Goat Serum, 0.01% Triton X-100). Primary antibody was prepared in PBS, and coverslips were incubated in primary antibody overnight at 4°C. Post-incubation, coverslips were first washed with 0.1% Triton X-100 in PBS and three times with PBS. Coverslips were incubated in the appropriate fluorochrome-conjugated antibody and protected from light at RT for 4 hours or less as needed. Coverslips were washed once with 0.1% Triton X-100 in PBS and three times with PBS. Coverslips were incubated for 10 minutes in 1 µg/ml DAPI (Premade DAPI solution from ThermoFisher (62248) and washed three times with PBS. Coverslips were mounted onto labeled slides and mounted with 15 µl of ProLong™ Glass Antifade Mountant (ThermoFisher: P36984). Coverslips cured for 24 hours before imaging. Unless otherwise indicated, all primary antibodies were used at a dilution of 1:1,000 and secondary antibodies at a dilution of 1:5,000.

For super-resolution microscopy, images were captured using the Nikon SoRa spinning disk microscope equipped with a 60x Apo TIRF oil immersion objective lens (N.A. 1.49) and Photometrics BSI sCMOS camera. Z-stacks were collected with 0.15 um step size and 0.03 um x-y pixel size. Within a single experiment or imaging time frame all imaging conditions were kept the same, and conditions were chosen to avoid saturation of the signal. Batch processing was performed on all collected images for deconvolution and channel alignment (based on TetraSpeck beads, 100 nm) These functions were conducted using standard macros run on the NIS software. Deconvolved, channel-aligned image files were converted to be compatible with the Imaris analysis software (v 9.8). Images were collected in respective channels across at least 30 z-steps across a nuclear size of 10 µm. Post-collection, images were deconvolved and channel aligned relative to appropriate controls (either wild-type cells or secondary-only controls) in channels in which images were obtained. Deconvolved, aligned images were background subtracted in the Imaris software. DAPI was used to generate a nuclear mask, and this mask was applied to the signal obtained in each channel. Overlap in each signal relative to the other was calculated using the Coloc function in the Imaris software. Statistics were exported.

For 3D-SIM and 2DSTED, coverslips with cells were fixed and permeabilized as described previously and mounted in a homemade mounting medium (90% glycerol, 100 mM Tris, pH 8.0, and 0.1 mg/ml *p*-phenylenediamine [695106; Sigma-Aldrich]).

3D-SIM was performed on N-SIM (Nikon), equipped with 405-,488-, 561-, and 640-nm excitation lasers, Apo TIRF 100× NA 1.49 Plan Apo oil objective, and a back-illuminated electron-multiplying charge-coupled device (CCD) camera (DU897; Andor). TetraSpeck beads (100 nm) were imaged in the same plane of view as representative nuclei presented and imaged in all relevant channels **(Figure S3).** Chanel alignment was performed using 100 nm TetraSpeck beads embedded in each sample. Single-channel orthogonal views representing whole nuclei or indicated insets were assembled in Fiji (National Institutes of Health).

2DSTED imaging was performed on the same samples prepared for 3D-SIM using STEDYCON (Abberior Instruments) assembled on an Eclipse Ti2 inverted microscope (Nikon) with a 100×, NA 1.45 Plan Apo objective. Signal was detected using Avalanche photo detectors (650–700, 575– 625, and 505–545 nm; DAPI detection) and Abebrior Instruments based control software was used to generate 2DSTED images. The following imaging conditions were used : excitation/detector wavelength, 561/616 nm and 640/660 nm for STAR ORANGE and STAR RED (Table S1), respectively; gate delay and width, 1 and 7 ns: Objective magn.: 100x; Pinhole: 32 µm; Pixel size:20 nm.

### Line plot profiles

Merged images or single channel images were obtained and the representative line plot drawn as indicated. Using the plot profile function in Fiji, the data was copied for the required channels. Data were exported to GraphPad prism and plotted as shown.

### Threshold setting for identification of lower and higher intensity mNG-EWSR1

Background subtracted images were imported into Fiji. For each image, the threshold cutoff to 3X times the minimum signal. A size exclusion parameter of 20 pixels was applied using the function “Analyze particles” to isolate foci. Number of foci and linear size was then obtained using the “Measure particles” function in Fiji. Data was then imported into GraphPad prism and plotted as shown.

### Mander’s overlap coefficient (MOC) and Pearson Correlation coefficient (PCC)

All deconvolved, channel-aligned, and background-subtracted images were used for the calculation of these metrics. mNG-EWSR1 foci were isolated as described above and signal intensity thresholds for proteins assessed via immunofluorescence were set based primary antibody only controls. The Coloc2 function in Fiji was used to obtain all necessary metrics across technical and independent replicates. Metrics were imported into GraphPad Prism and plotted as shown.

### Statistical analysis

Results shown as mean and standard error of the mean (S.E.M) were calculated using GraphPad Prism version 8.0.0 for Windows, GraphPad Software, San Diego, California USA, www.graphpad.com. Statistical analysis was conducted using an unpaired t-test with Welch’s correction in GraphPad Prism. A p-value of <0.05 was considered significant.

### Number of independent experiments performed in this study

**Figure 1**: Super-resolution confocal microscopy: Experiments were performed over five times with comparable results – data shown from multiple experiments. FRAP: Experiments were performed twice with ten replicates each with comparable results – data shown from the same experiment. **Figure S1**: Flow cytometry: Experiments were performed three times with comparable results – data shown from single experiment. Confocal microscopy: experiments were performed twice with comparable results – data shown from a single experiment. Flow cytometry: Experiments were performed three times – data shown from single experiment. Immunoblotting: Experiment was performed once. Super-resolution microscopy: Experiments were performed three times – data shown across multiple experiments. Figure 2: SIM: Experiments were performed greater than five times with comparable results – data shown from multiple experiments. **Figure S2**: Immunoblot was performed once. SIM microscopy: Experiments were performed two times with two independent replicates – data shown from a single experiment. Figure 3: 3D-SIM microscopy: Experiment were performed three times with comparable results – data shown from multiple experiments. STED microscopy: Experiment performed once across two independent replicates. Data shown from multiple experiments. Figure 4: STED microscopy: Experiments were performed once across two independent replicates with comparable results – data shown from multiple experiments. Figure 5: STED microscopy: Experiments were performed once across three independent replicates with comparable results – data shown from a single experiment. **Figure S3**: 3D-SIM microscopy: Experiments were performed three times across independent replicates with comparable results – data shown from a single experiment. Figure 6: 3D-SIM: Experiments were performed greater than three times with comparable results – data shown from a single experiment. STED microscopy: Experiments were performed once across two independent replicates with comparable results – data shown from a single experiment. **Figure S4:** Immunoprecipitation: Experiment performed twice – data shown from a single experiment. STED microscopy: Experiments were performed once across two independent replicates with comparable results – data shown from a single experiment. Figure 7: 3D**-**SIM microscopy: Experiments were performed two times across independent replicates with comparable findings – data shown from a single experiment. **Figure S5**: Cell cycle analysis: Experiment was performed twice with comparable results, data shown from a single experiment. Immunoblotting: Experiments were performed two times with comparable results – data presented from a single experiment shown. 3D**-**SIM microscopy: Experiments were performed two times across independent replicates with comparable findings – data shown from a single experiment. Figure 8: Immunoblotting: Experiment performed once. STED microscopy: Experiment was performed two times across independent replicates – data shown across a single experiment.

## Acknowledgments

The Intramural Research Program (IRP) of the National Cancer Institute (NCI), Center for Cancer Research (CCR), National Institutes of Health supported this study [ZIA BC 011704 to N.J.C. and ZIA BC 011459 to JL and ZIC BC 010858 supports the CCR Confocal Microscopy Core Facility]. The generation of CRISPR-Cas9 reagents included the support of Federal funds from the National Cancer Institute, National Institutes of Health, under Contract No. HHSN261201500003I. We are grateful to Javed Khan (Genetics Branch), and members of the Caplen laboratory for discussions and comments on the manuscript. We also thank Sanjit Mukherjee and Roshan Shrestha (Genetics Branch) for their technical advice and guidance and the CCR Sequencing Facility (long-read sequencing, Pac Bio), CCR Genomics Core (Sanger sequencing), and CCR Flow Cytometry Core, especially Karen Wolcott, for technical assistance. The content of this publication does not necessarily reflect the views or policies of the U.S. Department of Health and Human Services, nor does mention of trade names, commercial products, or organizations imply endorsement by the U.S. Government.

The authors declare no competing financial interests.

## Author contributions

S. S. Rajan and N.J. Caplen conceived and designed this study. S. S. Rajan conducted most experiments and analyzed the data. J. Loncarek supported the employment of microscopy (SIM and STED), performing experimentation and image and data analysis. V. Ebegboni performed the immunoprecipitation experiments, P. Pichling generated immunoblotting results, and K. Ludwig assisted with generating and characterizing the modified EWS cell lines. T.L. Jones contributed to the long-read sequence analysis of the modified EWS cell lines. R. Chari designed and generated the CRISPR -targeting reagents. M. Kruhlak contributed technical support and advice, particularly the employment of confocal and super-resolution microscopy, including FRAP analysis. A. Tran assisted with image analysis. N.J. Caplen supervised the project and conducted some of the data analysis. S.S. Rajan and N.J. Caplen wrote the manuscript. All the authors read and agreed on the publication of this manuscript.

## Notes

### Competing Interest Statement

The authors have declared no competing interest.

