## Supplementary Files for "EWSR1’s visual modalities are defined by its association with nucleic acids and RNA polymerase II"

Supplementary File 1

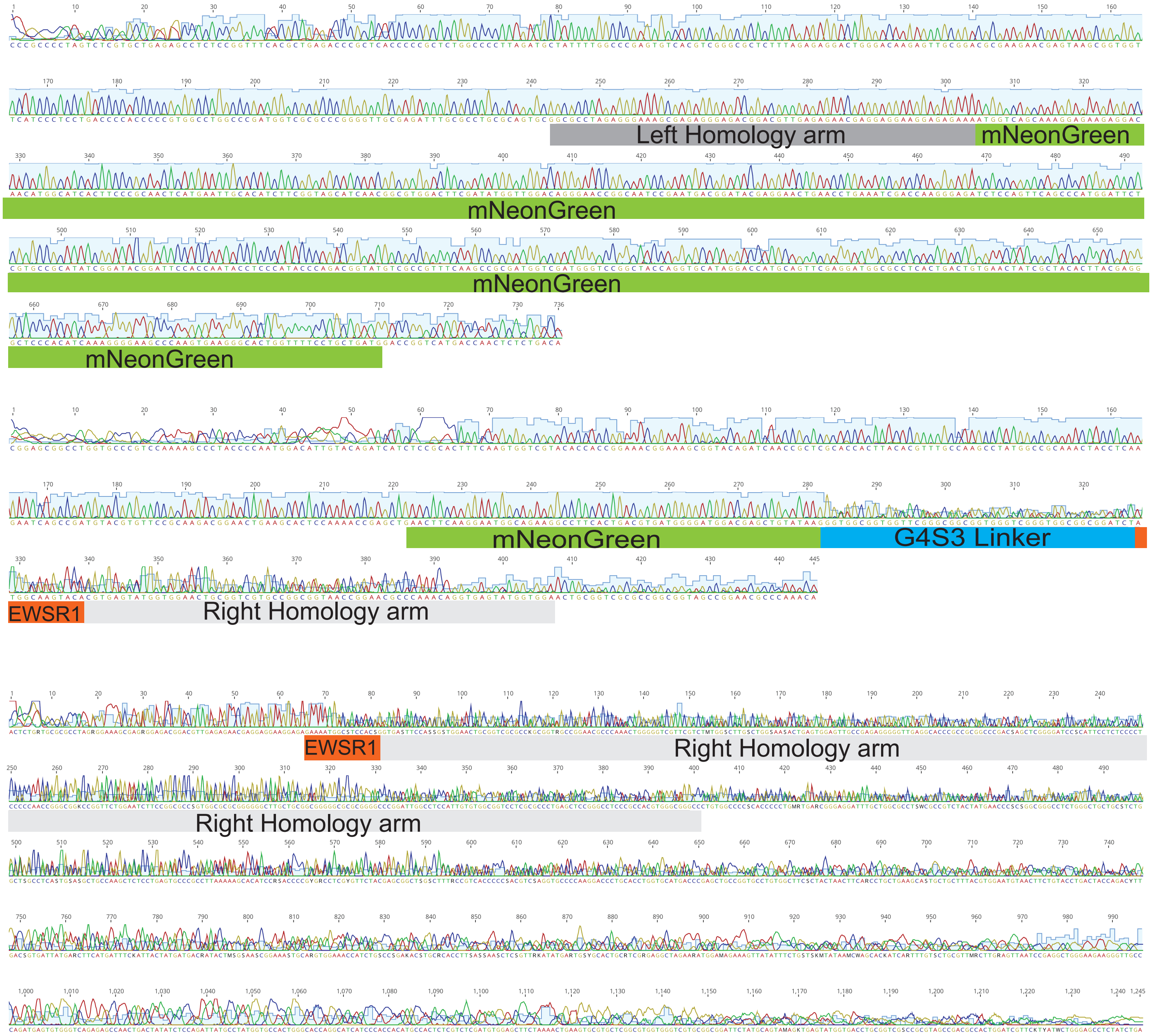

Representative sequence analysis confirming the correct insertion of the mNeonGreen containing DNA cassette in-frame with EWSR1

### Cellosaurus

Home | Contact

Due to scheduled maintenance work, this service and ftp://ftp.expasy.org will not be available from Friday January 15 7.00 p.m. until Saturday January 16 12.00 p.m. CEST.

Search Clear

#### Cellosaurus TC-32 (CVCL_7151)

[Text version]

| | |
|---|---|
| **Cell line name** | TC-32 |
| **Synonyms** | TC32 |
| **Accession** | CVCL_7151 |
| **Resource Identification Initiative** | To cite this cell line use: TC-32 (RRID:CVCL_7151) |
| **Comments** | Part of: Cancer Cell Line Encyclopedia (CCLE) project.<br>Part of: MD Anderson Cell Lines Project.<br>Doubling time: 24 hours (PubMed=24312454); 36 hours (PubMed=25984343).<br>Omics: Deep exome analysis.<br>Omics: Deep RNAseq analysis.<br>Omics: Protein expression by reverse-phase protein arrays. |
| **Sequence variations** | Gene fusion EWSR1-FLI1 (EWS-FLI1) (PubMed=8040301).<br>Homozygous for CDKN2A deletion (PubMed=25010205).<br>STAG2 p.Tyr636fs (PubMed=25010205; DepMap). |
| **Genome ancestry** | **Source:** PubMed=30894373 |
| **Disease** | Primitive neuroectodermal tumor (NCIt: C3716) |
| **Species of origin** | Homo sapiens (Human) (NCBI Taxonomy: 9606) |
| **Sex of cell** | Female |
| **Age at sampling** | 2Y7M |
| **Category** | Cancer cell line |
| **STR profile** | **Source(s):** PubMed=24312454; PubMed=25010205 |

Genome ancestry:

| Origin | % genome |
|---|---|
| African | 0.13 |
| Native American | 0.3 |
| East Asian, North | 1.12 |
| East Asian, South | 0 |
| South Asian | 0.2 |
| European, North | 67.6 |
| European, South | 30.65 |

**Markers:**

| | |
|---|---|
| Amelogenin | X |
| CSF1PO | 11,13 |
| D2S1338 | 17,19 |
| D3S1358 | 15,16 |
| D5S818 | 12,13 |
| D7S820 | 8,11 |
| D8S1179 | 12 |
| D13S317 | 10,12 |
| D16S539 | 13,14 |
| D18S51 | 11,19 |

| | |
|---|---|
| D19S433 | 12,14 |
| D21S11 | 27,28 |
| FGA | 23,24 |
| TH01 | 6,9.3 |
| TPOX | 9,11 |
| vWA | 15,18 |

**Run an STR similarity search on this cell line**

**Web pages**

http://www.cogcell.org/dl/EFT_Lines_DataSheets/TC-32_Cell_Line_Data_Sheet_COGcell_org.pdf
http://tcpaportal.org/mclp/

**Publications**

PubMed=3004699; DOI=10.1016/0165-4608(86)90001-4
Whang-Peng J., Triche T.J., Knutsen T., Miser J.S., Kao-Shan S., Tsai S., Israel M.A.
Cytogenetic characterization of selected small round cell tumors of childhood.
Cancer Genet. Cytogenet. 21:185-208(1986)

PubMed=3390826
McKeon C., Thiele C.J., Ross R.A., Kwan M., Triche T.J., Miser J.S., Israel M.A.
Indistinguishable patterns of protooncogene expression in two distinct but closely related tumors: Ewing's sarcoma and neuroepithelioma.
Cancer Res. 48:4307-4311(1988)

DOI=10.1016/B978-0-12-333530-2.50006-X
Israel M.A., Thiele C.J.
Tumor cell lines of the peripheral nervous system.
(In) Atlas of human tumor cell lines; Hay R.J., Park J.-G., Gazdar A.F. (eds.); pp.43-78; Academic Press; New York (1994)

PubMed=8040301; DOI=10.1172/JCI117360
Giovannini M., Biegel J.A., Serra M., Wang J.-Y., Wei Y.H., Nycum L., Emanuel B.S., Evans G.A.
EWS-erg and EWS-Fli1 fusion transcripts in Ewing's sarcoma and primitive neuroectodermal tumors with variant translocations.
J. Clin. Invest. 94:489-496(1994)

PubMed=16631476; DOI=10.1016/j.cancergencyto.2005.11.006
Szuhai K., Ijszenga M., Tanke H.J., Rosenberg C., Hogendoorn P.C.W.
Molecular cytogenetic characterization of four previously established and two newly established Ewing sarcoma cell lines.
Cancer Genet. Cytogenet. 166:173-179(2006)

PubMed=22142829; DOI=10.1158/1078-0432.CCR-11-2056
Shukla N., Ameur N., Yilmaz I., Nafa K., Lau C.-Y., Marchetti A., Borsu L., Barr F.G., Ladanyi M.
Oncogene mutation profiling of pediatric solid tumors reveals significant subsets of embryonal rhabdomyosarcoma and neuroblastoma with mutated genes in growth signaling pathways.
Clin. Cancer Res. 18:748-757(2012)

PubMed=24312454; DOI=10.1371/journal.pone.0080060
May W.A., Grigoryan R.S., Keshelava N., Cabral D.J., Christensen L.L., Jenabi J., Ji L., Triche T.J., Lawlor E.R., Reynolds C.P.
Characterization and drug resistance patterns of Ewing's sarcoma family tumor cell lines.
PLoS ONE 8:E80060-E80060(2013)

PubMed=25010205; DOI=10.1371/journal.pgen.1004475
Brohl A.S., Solomon D.A., Chang W., Wang J., Song Y., Sindiri S., Patidar R., Hurd L., Chen L., Shern J.F., Liao H., Wen X., Gerard J., Kim J.-S., Lopez Guerrero J.A., Machado I., Wai D.H., Picci P., Triche T.J., Horvai A.E., Miettinen M., Wei J.S., Catchpoole D., Llombart-Bosch A., Waldman T., Khan J.
The genomic landscape of the Ewing sarcoma family of tumors reveals recurrent STAG2 mutation.
PLoS Genet. 10:E1004475-E1004475(2014)

PubMed=25984343; DOI=10.1038/sdata.2014.35
Cowley G.S., Weir B.A., Vazquez F., Tamayo P., Scott J.A., Rusin S., East-Seletsky A., Ali L.D., Gerath W.F.J., Pantel S.E., Lizotte P.H., Jiang G., Hsiao J., Tsherniak A., Dwinell E., Aoyama S., Okamoto M., Harrington W., Gelfand E., Green T.M., Tomko M.J., Gopal S., Wong T.C., Li H., Howell S., Stransky N., Liefeld T., Jang D., Bistline J., Hill Meyers B., Armstrong S.A., Anderson K.C., Stegmaier K., Reich M., Pellman D., Boehm J.S., Mesirov J.P., Golub T.R., Root D.E., Hahn W.C.
Parallel genome-scale loss of function screens in 216 cancer cell lines for the identification of context-specific genetic dependencies.
Sci. Data 1:140035-140035(2014)

PubMed=26351324; DOI=10.1158/1535-7163.MCT-15-0074
Teicher B.A., Polley E.C., Kunkel M., Evans D., Silvers T.E., Delosh R.M., Laudeman J., Ogle C., Reinhart R., Selby M., Connelly J., Harris E., Monks A., Morris J.
Sarcoma cell line screen of oncology drugs and investigational agents identifies patterns associated with gene

| | |
|---|---|
| | and microRNA expression.<br>Mol. Cancer Ther. 14:2452-2462(2015)<br><br>PubMed=26979953; DOI=10.1073/pnas.1521251113<br>Town J., Pais H., Harrison S., Stead L.F., Bataille C., Bunjobpol W., Zhang J., Rabbitts T.H.<br>Exploring the surfaceome of Ewing sarcoma identifies a new and unique therapeutic target.<br>Proc. Natl. Acad. Sci. U.S.A. 113:3603-3608(2016)<br><br>PubMed=28196595; DOI=10.1016/j.ccell.2017.01.005<br>Li J., Zhao W., Akbani R., Liu W., Ju Z., Ling S., Vellano C.P., Roebuck P., Yu Q., Eterovic A.K., Byers L.A., Davies M.A., Deng W., Gopal Y.N.V., Chen G., von Euw E.M., Slamon D.J., Conklin D., Heymach J.V., Gazdar A.F., Minna J.D., Myers J.N., Lu Y., Mills G.B., Liang H.<br>Characterization of human cancer cell lines by reverse-phase protein arrays.<br>Cancer Cell 31:225-239(2017)<br><br>PubMed=30894373; DOI=10.1158/0008-5472.CAN-18-2747<br>Dutil J., Chen Z., Monteiro A.N., Teer J.K., Eschrich S.A.<br>An interactive resource to probe genetic diversity and estimated ancestry in cancer cell lines.<br>Cancer Res. 79:1263-1273(2019)<br><br>PubMed=31068700; DOI=10.1038/s41586-019-1186-3<br>Ghandi M., Huang F.W., Jane-Valbuena J., Kryukov G.V., Lo C.C., McDonald E.R. III, Barretina J., Gelfand E.T., Bielski C.M., Li H., Hu K., Andreev-Drakhlin A.Y., Kim J., Hess J.M., Haas B.J., Aguet F., Weir B.A., Rothberg M.V., Paolella B.R., Lawrence M.S., Akbani R., Lu Y., Tiv H.L., Gokhale P.C., de Weck A., Mansour A.A., Oh C., Shih J., Hadi K., Rosen Y., Bistline J., Venkatesan K., Reddy A., Sonkin D., Liu M., Lehar J., Korn J.M., Porter D.A., Jones M.D., Golji J., Caponigro G., Taylor J.E., Dunning C.M., Creech A.L., Warren A.C., McFarland J.M., Zamanighomi M., Kauffmann A., Stransky N., Imielinski M., Maruvka Y.E., Cherniack A.D., Tsherniak A., Vazquez F., Jaffe J.D., Lane A.A., Weinstock D.M., Johannessen C.M., Morrissey M.P., Stegmeier F., Schlegel R., Hahn W.C., Getz G., Mills G.B., Boehm J.S., Golub T.R., Garraway L.A., Sellers W.R.<br>Next-generation characterization of the Cancer Cell Line Encyclopedia.<br>Nature 569:503-508(2019) |
| **Cross-references** | |
| **Cell line databases/resources** | CCLE; TC32_BONE<br>Cell_Model_Passport; SIDM01465<br>DepMap; ACH-001205 |
| **Ontologies** | BTO; BTO:0004227 |
| **Chemistry resources** | PharmacoDB; TC32_1566_2019 |
| **Gene expression databases** | GEO; GSM1676322<br>GEO; GSM1701655<br>GEO; GSM1899423 |
| **Other** | Wikidata; Q54971813 |
| **Polymorphism and mutation databases** | Cosmic; 759891<br>Cosmic; 1078387<br>Cosmic; 1084513<br>Cosmic; 1718432<br>Cosmic; 2060803<br>Cosmic; 2228244<br>Cosmic; 2294588<br>Progenetix; CVCL_7151 |
| **Entry history** | |
| **Entry creation** | 04-Apr-2012 |
| **Last entry update** | 29-Oct-2020 |
| **Version number** | 20 |

Expasy is operated by the SIB Swiss Institute of Bioinformatics | Terms of Use

Back to the top

### Cell Line Authentication Service
#### STR Profile Report

| | |
|---|---|
| **Sample Submitted By:** | National Institutes of Health<br>Allison Cross |
| **Email Address:** | |
| **ATCC Sales Order:** | SO0312848 |
| **FTA Barcode:** | STRA9963 |
| **Cell Line Designation:** | TC-32 |
| **Date Sample Received:** | Tuesday, October 15, 2019 |
| **Report Date:** | Thursday, October 17, 2019 |

**Methodology:** Seventeen short tandem repeat (STR) loci plus the gender determining locus, Amelogenin, were amplified using the commercially available PowerPlex® 18D Kit from Promega. The cell line sample was processed using the ABI Prism® 3500xl Genetic Analyzer. Data were analyzed using GeneMapper® ID-X v1.2 software (Applied Biosystems). Appropriate positive and negative controls were run and confirmed for each sample submitted.

**Data Interpretation:** Cell lines were authenticated using Short Tandem Repeat (STR) analysis as described in 2012 in ANSI Standard (ASN-0002) Authentication of Human Cell Lines: Standardization of STR Profiling by the ATCC Standards Development Organization (SDO) and in Capes-Davis et al., Match criteria for human cell line authentication: Where do we draw the line? Int. J. Cancer. 2012 Nov 8. doi: 10.1002/ijc.27931

**ATCC performs STR Profiling following ISO 9001:2008 and ISO/IEC 17025:2005 quality standards.**

There are no warranties with respect to the services or results supplied, express or implied, including, without limitation, any implied warranty of merchantability or fitness for a particular purpose. Neither ATCC nor Promega is liable for any damages or injuries resulting from receipt and/or improper, inappropriate, negligent or other wrongful use of the test results supplied, and/or from misidentification, misrepresentation, or lack of accuracy of those results. Your exclusive remedy against ATCC, Promega and those supplying materials used in the services for any losses or damage of any kind whatsoever, whether in contract, tort, or otherwise, shall be, at Promega's option, refund of the fee paid for such service or repeat of the service.

The ATCC trademark and trade name, any and all ATCC catalog numbers are trademarks of the American Type Culture Collection. PowerPlex is a registered trademark of Promega Corporation. Applied Biosystems, ABI Prism and GeneMapper are registered trademarks of Life Technologies Corporation.

**Technical questions?**

ATCC Technical Support
(800) 638-6597 / +1 703-365-2700


**Ordering questions?**

800-638-6597 or 703-365-2700


### Cell Line Authentication Service
#### STR Profile Report

**FTA Barcode:** STRA9963

**ATCC Sales Order:** SO0312848

| Test Results for Submitted Sample | | | | | ATCC Reference Database Profile | | | |
|---|---|---|---|---|---|---|---|---|
| **Locus** | **Query Profile: TC-32** | | | | **Database Profile:** | | | |
| **D3S1358** | 15 | 16 | | | | | | |
| **TH01** | 6 | 9.3 | | | | | | |
| **D21S11** | 27 | 28 | | | | | | |
| **D18S51** | 11 | 19 | | | | | | |
| **Penta_E** | 12 | 13 | | | | | | |
| **D5S818** | 12 | 13 | | | | | | |
| **D13S317** | 10 | 12 | | | | | | |
| **D7S820** | 8 | 11 | | | | | | |
| **D16S539** | 14 | | | | | | | |
| **CSF1PO** | 11 | 13 | | | | | | |
| **Penta_D** | 10 | 12 | | | | | | |
| **Amelogenin** | X | | | | | | | |
| **vWA** | 15 | 18 | | | | | | |
| **D8S1179** | 12 | | | | | | | |
| **TPOX** | 9 | 11 | | | | | | |
| **FGA** | 23 | 24 | | | | | | |
| **D19S433** | 12 | 14 | | | | | | |
| **D2S1338** | 17 | 19 | | | | | | |
| Number of shared alleles between query sample and database profile: | | | | | | | | NA |
| Total number of alleles in the database profile: | | | | | | | | NA |
| Percent match between the submitted sample and the database profile: | | | | | | | | NA |
| *The allele match algorithm compares the 8 core loci plus amelogenin only, even though alleles from all loci will be reported when available.* | | | | | | | | |
| **NOTE:** *Loci highlighted in grey (8 core STR loci plus Amelogenin) can be made public to verify cell identity. In order to protect the identity of the donor,* ***please do not publish*** *the allele calls from all the STR loci tested.* *Electropherograms showing raw data are attached.* | | | | | | | | |

**Explanation of Test Results**
Cell lines with 80% match are considered to be related; i.e., derived from a common ancestry. Cell lines with between a 55% to 80% match require further profiling for authentication of relatedness.

- [X] The submitted sample profile is human, but not a match for any profile in the ATCC STR database.
- [ ] The submitted profile is an exact match for the following ATCC human cell line(s) in the ATCC STR database (8 core loci plus Amelogenin):
- [ ] The submitted profile is similar to the following ATCC human cell line(s):
- [ ] An STR profile could not be generated.

**Additional Comments:**

Submitted sample, STRA9963 (TC-32), is not a match to any cell line in either the ATCC or DSMZ STR database. However, the profile for the submitted sample is a similar match to the STR profile for this cell line that is listed on the ExPASy website.
https://web.expasy.org/cellosaurus

| | |
|---|---|
| e-Signature, Technician: | wklein 10/17/2019 |
| e-Signature, Reviewer: | Bchase 10/17/2019 |

CELL LINE AUTHENTICATION SERVICE ATCC VERIFIED EST. 1925

**Cell Line Authentication Service**
STR Profile Report

**FTA Barcode:** STRA9963

**ATCC Sales Order:** SO0312848

**Addendum: Comparative Output from the ATCC STR Profile Database**

| % Match | ATCC® Cat. No. | Designation | D5S818 | D13S317 | D7S820 | D16S539 | vWA | TH01 | AMEL | TPOX | CSF1PO |
|---|---|---|---|---|---|---|---|---|---|---|---|
| 100 | STRA9963 | TC-32 | 12,13 | 10,12 | 8,11 | 14 | 15,18 | 6,9.3 | X | 9,11 | 11,13 |

**Definitions of terms used in this report:**

**Peak Area Difference (PAD):**
Refers to a heterozygous peak imbalance.
Two alleles at a single locus should amplify in a similar manner; and therefore produce peaks of similar height and area. Peaks which are above threshold (50 rfu) but are not of similar area, within 50% of each other, are referred to as a PAD. Due to their nature cell lines do not amplify in the same manner as a sample taken from a fresh buccal swab. PAD is far more common in cell line samples.

**Stutter:**
A stutter peak is a small peak which occurs immediately before the true peak. It is defined as being a single repeat unit smaller than the true peak. The stutter peak should be less than 15% of the true peak. The stutter is caused by the polymerase.

**+4 Peak:**
A +4 is similar to a stutter but occurs immediately after the true peak. A stutter peak should be less than 5% for a homozygous and 10% for a heterozygous.

**Below Threshold Peak(s):**
Cell lines can produce unusual profiles and occasionally a peak will amplify poorly and be below threshold. Where we find a below threshold peak which we believe is valid we indicate it as a below threshold peak. Our cell line analysis criteria, Homozygous and Heterozygous peaks must be equal to or above the set height threshold for it to be considered a true peak.

**Ladder/ Off Ladder Peak(s):**
The allelic ladder consists of most or all known alleles in the population and allows for precise assignment of alleles. Those which do not align are termed ‘off ladder.

**Artifact:**
A non-allelic product of the amplification process, an anomaly of the detection process, or a by-product of primer synthesis

**Pull-up:**
A term used to describe when signal from one dye color channel produces artificial peaks in another, usually adjacent, color.

**Spike:**
An extraneous peak resulting from dust, dried polymer, an air bubble, or an electrical surge.
**Dye blob:**
Free dye not coupled to primer that can be injected into the capillary (A known and documented dye blob is often found at the D3S1358 locus.)

### Cell Line Authentication Service
#### STR Profile Report

| | |
|---|---|
| **Sample Submitted By:** | National Institutes of Health<br>Soumya Saundara Rajan |
| **Email Address:** | |
| **ATCC Sales Order:** | SO0312848 |
| **FTA Barcode:** | STRA9978 |
| **Cell Line Designation:** | A-673 |
| **Date Sample Received:** | Tuesday, November 05, 2019 |
| **Report Date:** | Thursday, November 07, 2019 |

**Methodology:** Seventeen short tandem repeat (STR) loci plus the gender determining locus, Amelogenin, were amplified using the commercially available PowerPlex® 18D Kit from Promega. The cell line sample was processed using the ABI Prism® 3500xl Genetic Analyzer. Data were analyzed using GeneMapper® ID-X v1.2 software (Applied Biosystems). Appropriate positive and negative controls were run and confirmed for each sample submitted.

**Data Interpretation:** Cell lines were authenticated using Short Tandem Repeat (STR) analysis as described in 2012 in ANSI Standard (ASN-0002) Authentication of Human Cell Lines: Standardization of STR Profiling by the ATCC Standards Development Organization (SDO) and in Capes-Davis et al., Match criteria for human cell line authentication: Where do we draw the line? Int. J. Cancer. 2012 Nov 8. doi: 10.1002/ijc.27931

**ATCC performs STR Profiling following ISO 9001:2008 and ISO/IEC 17025:2005 quality standards.**

There are no warranties with respect to the services or results supplied, express or implied, including, without limitation, any implied warranty of merchantability or fitness for a particular purpose. Neither ATCC nor Promega is liable for any damages or injuries resulting from receipt and/or improper, inappropriate, negligent or other wrongful use of the test results supplied, and/or from misidentification, misrepresentation, or lack of accuracy of those results. Your exclusive remedy against ATCC, Promega and those supplying materials used in the services for any losses or damage of any kind whatsoever, whether in contract, tort, or otherwise, shall be, at Promega's option, refund of the fee paid for such service or repeat of the service.

The ATCC trademark and trade name, any and all ATCC catalog numbers are trademarks of the American Type Culture Collection. PowerPlex is a registered trademark of Promega Corporation. Applied Biosystems, ABI Prism and GeneMapper are registered trademarks of Life Technologies Corporation.

**Technical questions?**

ATCC Technical Support
(800) 638-6597 / +1 703-365-2700


**Ordering questions?**

800-638-6597 or 703-365-2700


### Cell Line Authentication Service
#### STR Profile Report

**FTA Barcode:** STRA9978

**ATCC Sales Order:** SO0312848

| Test Results for Submitted Sample | | | | | ATCC Reference Database Profile | |
|---|---|---|---|---|---|---|
| **Locus** | **Query Profile: A-673** | | | | **Database Profile: A-673; Ewing's Sarcoma; Human (Homo sapiens)** | |
| **D3S1358** | 14 | | | | | |
| **TH01** | 9.3 | | | | 9.3 | |
| **D21S11** | 29 | 30.2 | | | | |
| **D18S51** | 13 | 16 | | | | |
| **Penta_E** | 10 | 13 | | | | |
| **D5S818** | 11 | 12 | | | 11 | 12 |
| **D13S317** | 8 | 13 | | | 8 | 13 |
| **D7S820** | 10 | 12 | | | 10 | 12 |
| **D16S539** | 11 | | | | 11 | |
| **CSF1PO** | 11 | 12 | | | 11 | 12 |
| **Penta_D** | 12 | 13 | | | | |
| **Amelogenin** | X | | | | X | |
| **vWA** | 15 | 18 | | | 15 | 18 |
| **D8S1179** | 11 | 13 | | | | |
| **TPOX** | 8 | | | | 8 | |
| **FGA** | 19 | 20 | | | | |
| **D19S433** | 13 | 14 | | | | |
| **D2S1338** | 16 | 21 | | | | |

| | |
|---|---|
| Number of shared alleles between query sample and database profile: | 14 |
| Total number of alleles in the database profile: | 14 |
| Percent match between the submitted sample and the database profile: | 100 |

*The allele match algorithm compares the 8 core loci plus amelogenin only, even though alleles from all loci will be reported when available.*

**NOTE:** *Loci highlighted in grey (8 core STR loci plus Amelogenin) can be made public to verify cell identity. In order to protect the identity of the donor,* ***please do not publish*** *the allele calls from all the STR loci tested.*
*Electropherograms showing raw data are attached.*

**Explanation of Test Results**
Cell lines with 80% match are considered to be related; i.e., derived from a common ancestry. Cell lines with between a 55% to 80% match require further profiling for authentication of relatedness.

- [ ] The submitted sample profile is human, but not a match for any profile in the ATCC STR database.
- [X] The submitted profile is an exact match for the following ATCC human cell line(s) in the ATCC STR database (8 core loci plus Amelogenin): CRL-1598
- [ ] The submitted profile is similar to the following ATCC human cell line(s):
- [ ] An STR profile could not be generated.

**Additional Comments:**

Submitted sample, STRA9978 (A-673), is an exact match to ATCC cell line CRL-1598 (A-673).

| | |
|---|---|
| e-Signature, Technician: | snicholson 11/7/2019 |
| e-Signature, Reviewer: | Bchase 11/7/2019 |

**FTA Barcode:** STRA9978

**ATCC Sales Order:** SO0312848

##### Addendum: Comparative Output from the ATCC STR Profile Database

| % Match | ATCC® Cat. No. | Designation | D5S818 | D13S317 | D7S820 | D16S539 | vWA | TH01 | AMEL | TPOX | CSF1PO |
|---|---|---|---|---|---|---|---|---|---|---|---|
| 100 | STRA9978 | A-673 | 11,12 | 8,13 | 10,12 | 11 | 15,18 | 9.3 | X | 8 | 11,12 |
| 100 | CRL-1598 | A-673; Ewing's Sarcoma; Human (Homo sapiens) | 11,12 | 8,13 | 10,12 | 11 | 15,18 | 9.3 | X | 8 | 11,12 |

##### Definitions of terms used in this report:

**Peak Area Difference (PAD):**
Refers to a heterozygous peak imbalance.
Two alleles at a single locus should amplify in a similar manner; and therefore produce peaks of similar height and area. Peaks which are above threshold (50 rfu) but are not of similar area, within 50% of each other, are referred to as a PAD. Due to their nature cell lines do not amplify in the same manner as a sample taken from a fresh buccal swab. PAD is far more common in cell line samples.

**Stutter:**
A stutter peak is a small peak which occurs immediately before the true peak. It is defined as being a single repeat unit smaller than the true peak. The stutter peak should be less than 15% of the true peak. The stutter is caused by the polymerase.

**+4 Peak:**
A +4 is similar to a stutter but occurs immediately after the true peak. A stutter peak should be less than 5% for a homozygous and 10% for a heterozygous.

**Below Threshold Peak(s):**
Cell lines can produce unusual profiles and occasionally a peak will amplify poorly and be below threshold. Where we find a below threshold peak which we believe is valid we indicate it as a below threshold peak. Our cell line analysis criteria, Homozygous and Heterozygous peaks must be equal to or above the set height threshold for it to be considered a true peak.

**Ladder/ Off Ladder Peak(s):**
The allelic ladder consists of most or all known alleles in the population and allows for precise assignment of alleles. Those which do not align are termed ‘off ladder.

**Artifact:**
A non-allelic product of the amplification process, an anomaly of the detection process, or a by-product of primer synthesis

**Pull-up:**
A term used to describe when signal from one dye color channel produces artificial peaks in another, usually adjacent, color.

**Spike:**
An extraneous peak resulting from dust, dried polymer, an air bubble, or an electrical surge.
**Dye blob:**
Free dye not coupled to primer that can be injected into the capillary (A known and documented dye blob is often found at the D3S1358 locus.)

Applied Biosystems
GeneMapper® ID-X 1.2

Project: 20191105-WK-STR

| Sample Name | SQ | CGQ |
|---|---|---|
| STRA9978_A-673 | | |

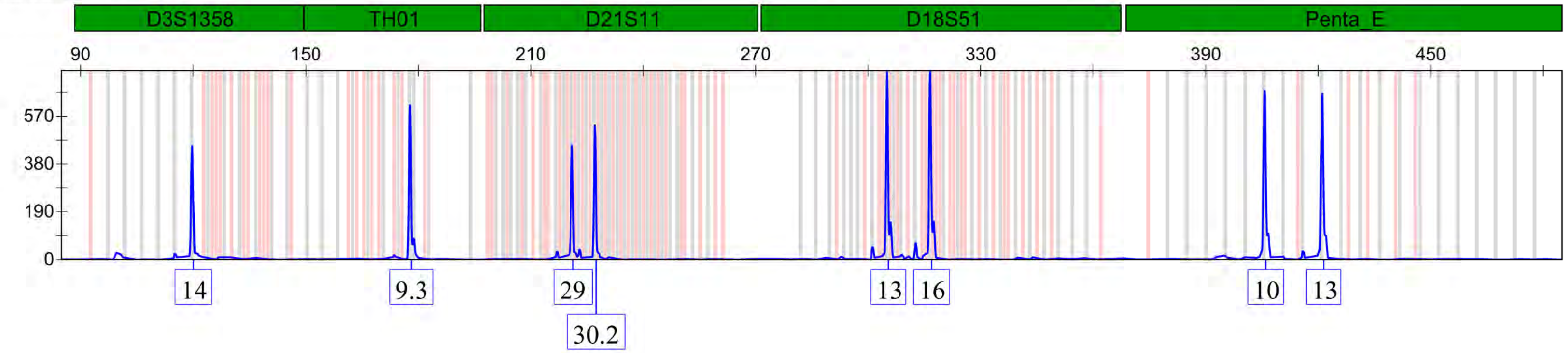

STRA9978_A-673

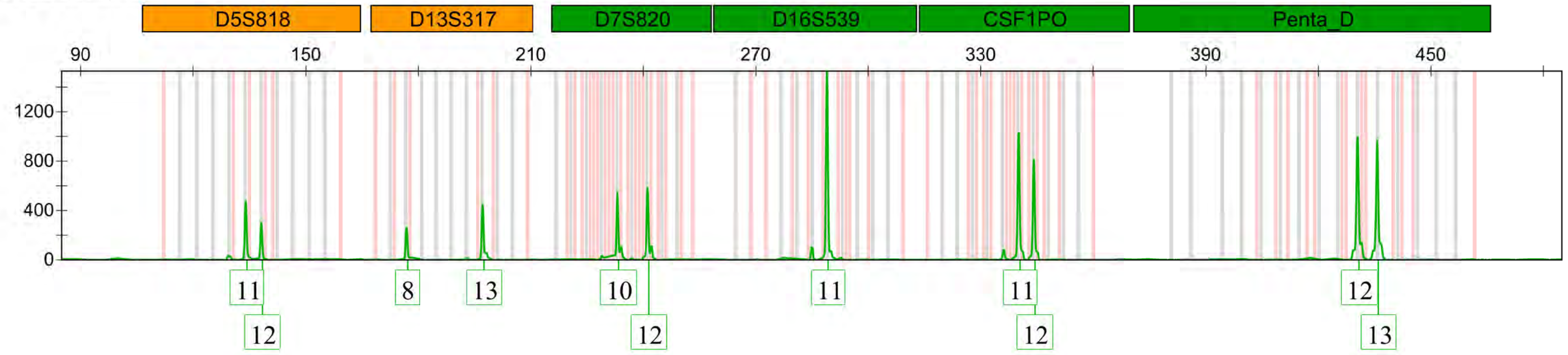

Project: 20191105-WK-STR

| Sample Name | SQ | CGQ |
|---|---|---|
| STRA9978_A-673 | | |

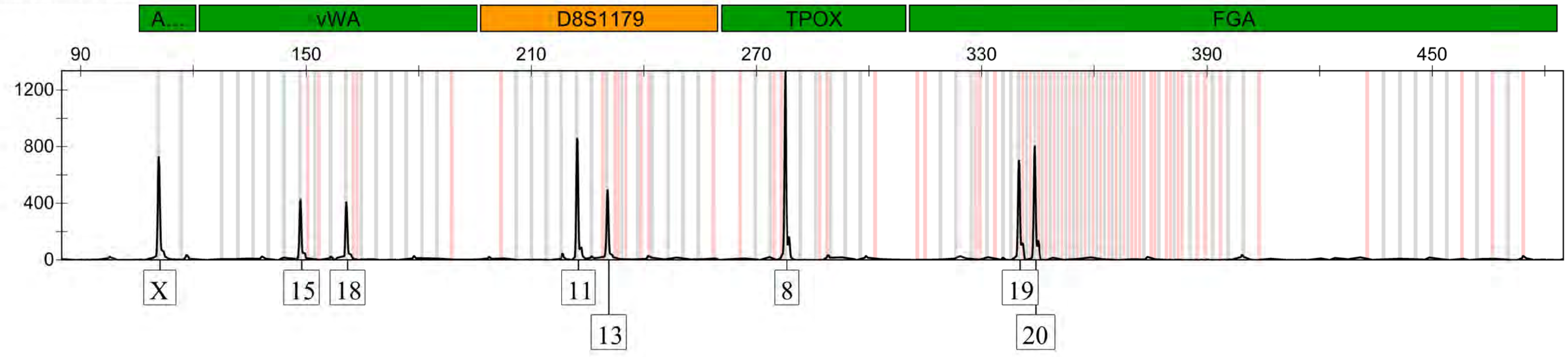

STRA9978_A-673

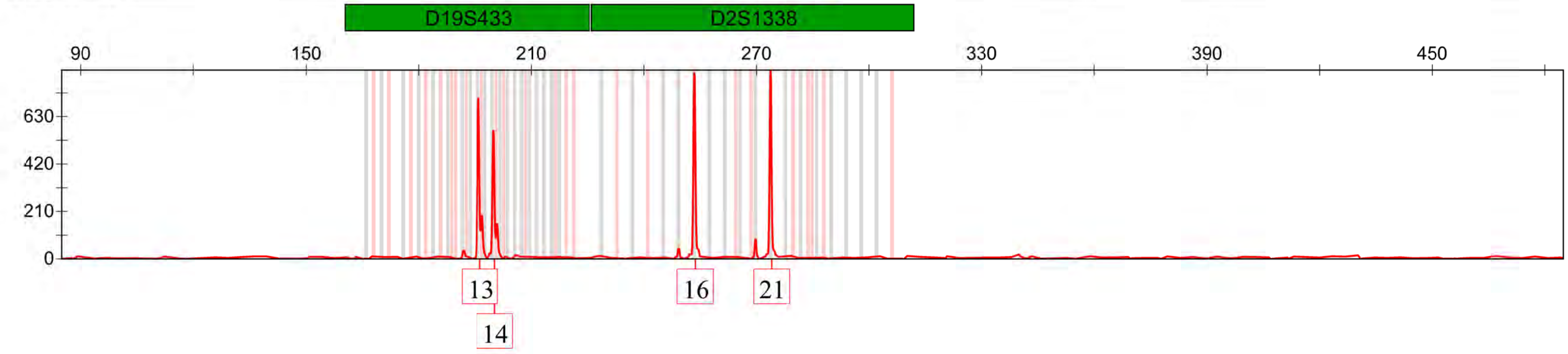

### Cell Line Authentication Service

#### STR Profile Report

| | |
|---|---|
| **Sample Submitted By:** | NIH<br>Soumya Sundara Rajan |
| **Email Address:** | |
| **ATCC Sales Order:** | SO0533550 |
| **FTA Barcode:** | STRB2775 |
| **Cell Line Designation:** | A673-pLE0567_#4 |
| **Date Sample Received:** | Wednesday, November 18, 2020 |
| **Report Date:** | Tuesday, November 24, 2020 |

**Methodology:** Seventeen short tandem repeat (STR) loci plus the gender determining locus, Amelogenin, were amplified using the commercially available PowerPlex® 18D Kit from Promega. The cell line sample was processed using the ABI Prism® 3500xl Genetic Analyzer. Data were analyzed using GeneMapper® ID-X v1.2 software (Applied Biosystems). Appropriate positive and negative controls were run and confirmed for each sample submitted.

**Data Interpretation:** Cell lines were authenticated using Short Tandem Repeat (STR) analysis as described in 2012 in ANSI Standard (ASN-0002) Authentication of Human Cell Lines: Standardization of STR Profiling by the ATCC Standards Development Organization (SDO) and in Capes-Davis et al., Match criteria for human cell line authentication: Where do we draw the line? Int. J. Cancer. 2012 Nov 8. doi: 10.1002/ijc.27931

**ATCC performs STR Profiling following ISO 9001:2008 and ISO/IEC 17025:2005 quality standards.**
There are no warranties with respect to the services or results supplied, express or implied, including, without limitation, any implied warranty of merchantability or fitness for a particular purpose. Neither ATCC nor Promega is liable for any damages or injuries resulting from receipt and/or improper, inappropriate, negligent or other wrongful use of the test results supplied, and/or from misidentification, misrepresentation, or lack of accuracy of those results. Your exclusive remedy against ATCC, Promega and those supplying materials used in the services for any losses or damage of any kind whatsoever, whether in contract, tort, or otherwise, shall be, at Promega's option, refund of the fee paid for such service or repeat of the service.

The ATCC trademark and trade name, any and all ATCC catalog numbers are trademarks of the American Type Culture Collection. PowerPlex is a registered trademark of Promega Corporation. Applied Biosystems, ABI Prism and GeneMapper are registered trademarks of Life Technologies Corporation.

**Technical questions?**

ATCC Technical Support
(800) 638-6597 / +1 703-365-2700


**Ordering questions?**

800-638-6597 or 703-365-2700


**Cell Line Authentication Service**
STR Profile Report

**FTA Barcode:** STRB2775

**ATCC Sales Order:** SO0533550

| Test Results for Submitted Sample | | | | | ATCC Reference Database Profile | | | |
|---|---|---|---|---|---|---|---|---|
| **Locus** | **Query Profile: A673-pLE0567_#4** | | | | **Database Profile: A-673 Ewing's Sarcoma Human (Homo sapiens)** | | | |
| **D3S1358** | 14 | | | | | | | |
| **TH01** | 9.3 | | | | 9.3 | | | |
| **D21S11** | 29 | 30.2 | | | | | | |
| **D18S51** | 13 | 16 | | | | | | |
| **Penta_E** | 10 | 13 | | | | | | |
| **D5S818** | 11 | 12 | | | 11 | 12 | | |
| **D13S317** | 8 | 13 | | | 8 | 13 | | |
| **D7S820** | 10 | 12 | | | 10 | 12 | | |
| **D16S539** | 11 | | | | 11 | | | |
| **CSF1PO** | 11 | 12 | | | 11 | 12 | | |
| **Penta_D** | 12 | 13 | | | | | | |
| **Amelogenin** | X | | | | X | | | |
| **vWA** | 15 | 18 | | | 15 | 18 | | |
| **D8S1179** | 11 | 13 | | | | | | |
| **TPOX** | 8 | | | | 8 | | | |
| **FGA** | 19 | 20 | | | | | | |
| **D19S433** | 13 | 14 | | | | | | |
| **D2S1338** | 16 | 21 | | | | | | |
| Number of shared alleles between query sample and database profile: | | | | | | | | 14 |
| Total number of alleles in the database profile: | | | | | | | | 14 |
| Percent match between the submitted sample and the database profile: | | | | | | | | 100 |

*The allele match algorithm compares the 8 core loci plus amelogenin only, even though alleles from all loci will be reported when available.*

**NOTE:** *Loci highlighted in grey (8 core STR loci plus Amelogenin) can be made public to verify cell identity. In order to protect the identity of the donor,* ***please do not publish*** *the allele calls from all the STR loci tested.*
*Electropherograms showing raw data are attached.*

**Explanation of Test Results**
Cell lines with 80% match are considered to be related; i.e., derived from a common ancestry. Cell lines with between a 55% to 80% match require further profiling for authentication of relatedness.

[ ] The submitted sample profile is human, but not a match for any profile in the ATCC STR database.

[X] The submitted profile is an exact match for the following ATCC human cell line(s) in the ATCC STR database (8 core loci plus Amelogenin): CRL-1598

[ ] The submitted profile is similar to the following ATCC human cell line(s):

[ ] An STR profile could not be generated.

**Additional Comments:**

Submitted sample, STRB2775 (A673-pLE0567_#4), is an exact match to ATCC cell line CRL-1598 (A-673).

| | |
|---|---|
| e-Signature, Technician: | snicholson 11/24/2020 |
| e-Signature, Reviewer: | gsykes 11/24/2020 |

**Addendum: Comparative Output from the ATCC STR Profile Database**

| % Match | ATCC® Cat. No. | Designation | D5S818 | D13S317 | D7S820 | D16S539 | vWA | TH01 | AMEL | TPOX | CSF1PO |
|---|---|---|---|---|---|---|---|---|---|---|---|
| 100 | STRB2775 | A673-pLE0567_#4 | 11,12 | 8,13 | 10,12 | 11 | 15,18 | 9.3 | X | 8 | 11,12 |
| 100 | CRL-1598 | A-673 Ewing's Sarcoma Human (Homo sapiens) | 11,12 | 8,13 | 10,12 | 11 | 15,18 | 9.3 | X | 8 | 11,12 |

**Definitions of terms used in this report:**

**Peak Area Difference (PAD):**
Refers to a heterozygous peak imbalance.
Two alleles at a single locus should amplify in a similar manner; and therefore produce peaks of similar height and area. Peaks which are above threshold (50 rfu) but are not of similar area, within 50% of each other, are referred to as a PAD. Due to their nature cell lines do not amplify in the same manner as a sample taken from a fresh buccal swab. PAD is far more common in cell line samples.

**Stutter:**
A stutter peak is a small peak which occurs immediately before the true peak. It is defined as being a single repeat unit smaller than the true peak. The stutter peak should be less than 15% of the true peak. The stutter is caused by the polymerase.

**+4 Peak:**
A +4 is similar to a stutter but occurs immediately after the true peak. A stutter peak should be less than 5% for a homozygous and 10% for a heterozygous.

**Below Threshold Peak(s):**
Cell lines can produce unusual profiles and occasionally a peak will amplify poorly and be below threshold. Where we find a below threshold peak which we believe is valid we indicate it as a below threshold peak. Our cell line analysis criteria, Homozygous and Heterozygous peaks must be equal to or above the set height threshold for it to be considered a true peak.

**Ladder/ Off Ladder Peak(s):**
The allelic ladder consists of most or all known alleles in the population and allows for precise assignment of alleles. Those which do not align are termed ‘off ladder.

**Artifact:**
A non-allelic product of the amplification process, an anomaly of the detection process, or a by-product of primer synthesis

**Pull-up:**
A term used to describe when signal from one dye color channel produces artificial peaks in another, usually adjacent, color.

**Spike:**
An extraneous peak resulting from dust, dried polymer, an air bubble, or an electrical surge.
**Dye blob:**
Free dye not coupled to primer that can be injected into the capillary (A known and documented dye blob is often found at the D3S1358 locus.)

Applied Biosystems
GeneMapper® ID-X 1.2

Project: 20201119-WK

| Sample Name | SQ | CGQ |
|---|---|---|
| STRB2775_A673-pLE0567_#4 | | |

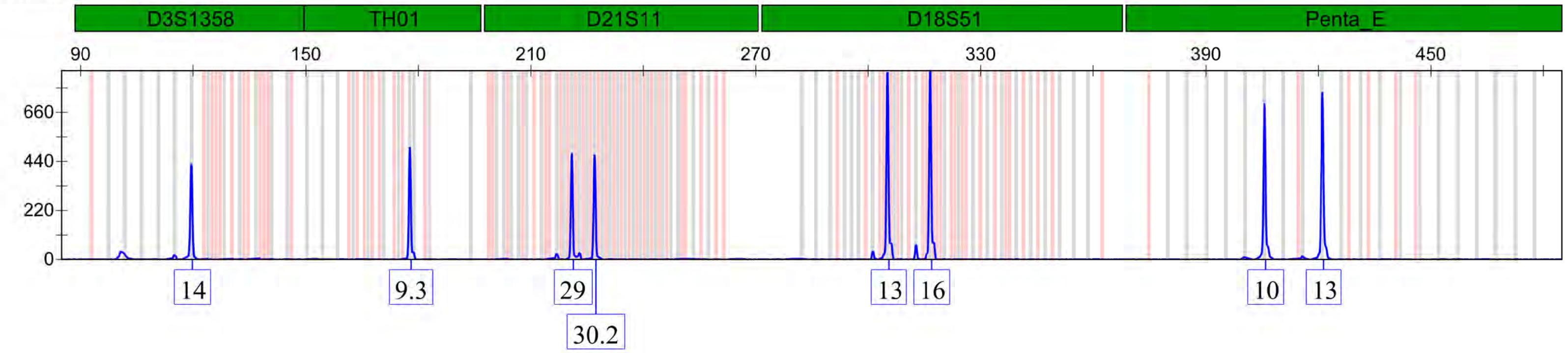

| |
|---|
| STRB2775_A673-pLE0567_#4 |

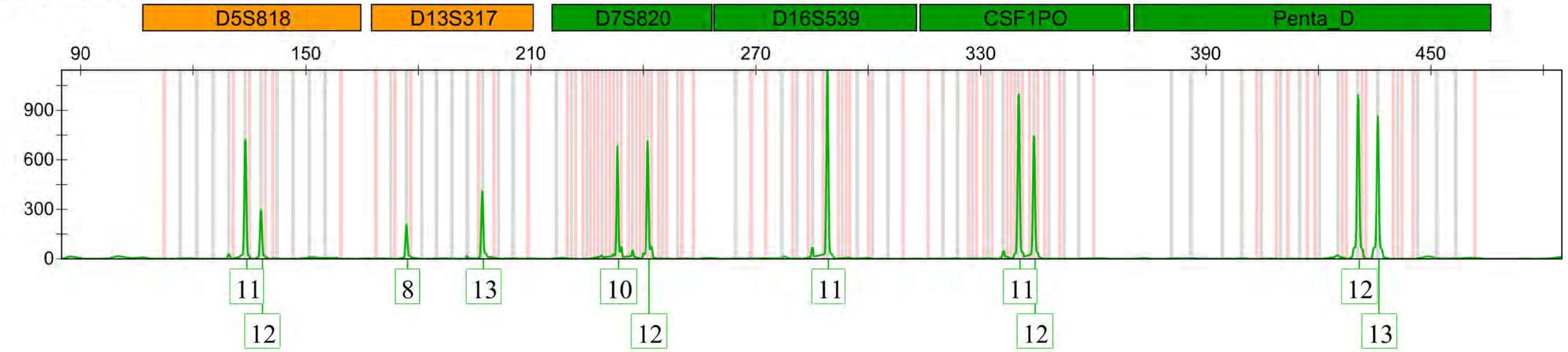

Applied Biosystems
GeneMapper® ID-X 1.2

Project: 20201119-WK

| Sample Name | SQ | CGQ |
|---|---|---|
| STRB2775_A673-pLE0567_#4 | | |

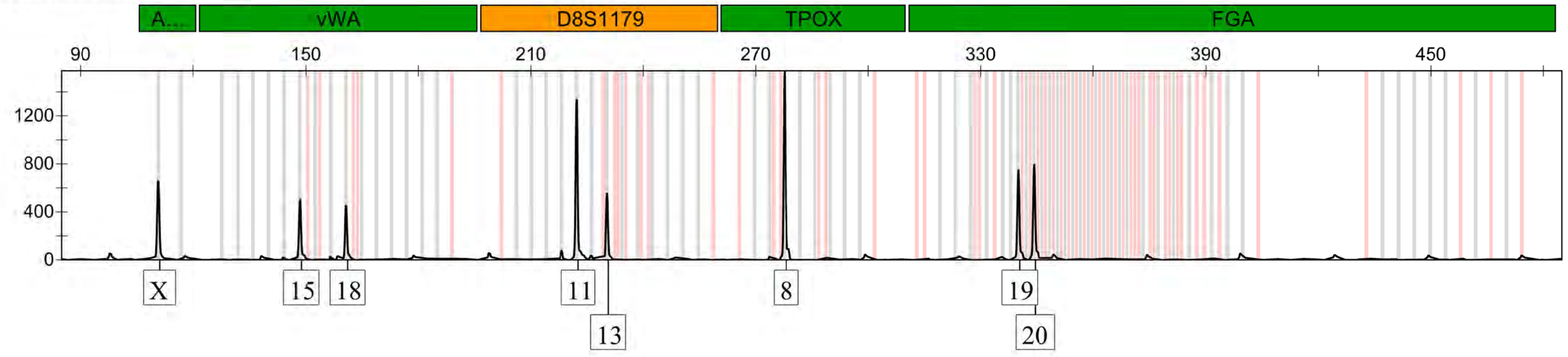

| |
|---|
| STRB2775_A673-pLE0567_#4 |

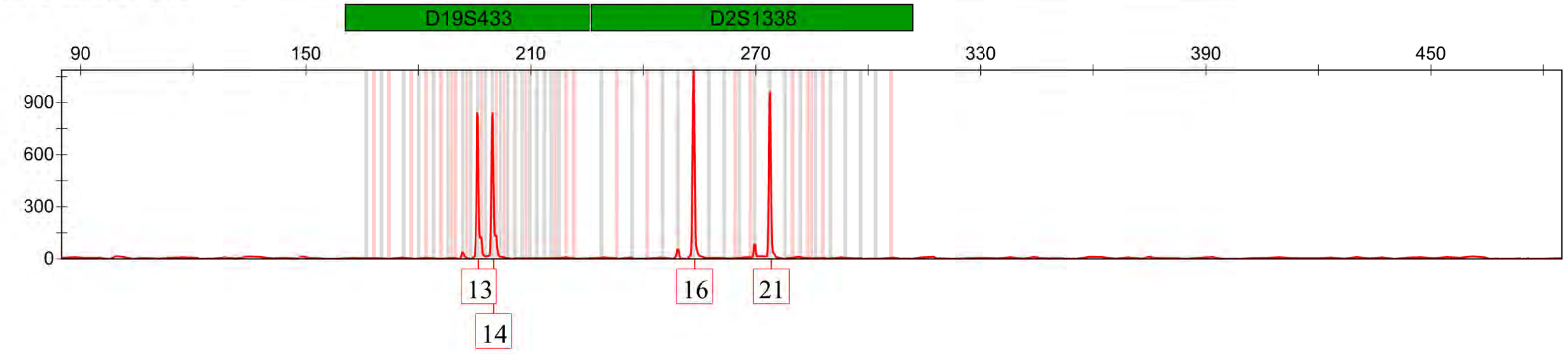

### Cell Line Authentication Service
#### STR Profile Report

| | |
|---|---|
| **Sample Submitted By:** | NIH<br>Soumya Sundara Rajan |
| **Email Address:** | |
| **ATCC Sales Order:** | SO0533550 |
| **FTA Barcode:** | STRB2778 |
| **Cell Line Designation:** | TC32-pLE0567_#6 |
| **Date Sample Received:** | Wednesday, November 18, 2020 |
| **Report Date:** | Tuesday, November 24, 2020 |

**Methodology:** Seventeen short tandem repeat (STR) loci plus the gender determining locus, Amelogenin, were amplified using the commercially available PowerPlex® 18D Kit from Promega. The cell line sample was processed using the ABI Prism® 3500xl Genetic Analyzer. Data were analyzed using GeneMapper® ID-X v1.2 software (Applied Biosystems). Appropriate positive and negative controls were run and confirmed for each sample submitted.

**Data Interpretation:** Cell lines were authenticated using Short Tandem Repeat (STR) analysis as described in 2012 in ANSI Standard (ASN-0002) Authentication of Human Cell Lines: Standardization of STR Profiling by the ATCC Standards Development Organization (SDO) and in Capes-Davis et al., Match criteria for human cell line authentication: Where do we draw the line? Int. J. Cancer. 2012 Nov 8. doi: 10.1002/ijc.27931

**ATCC performs STR Profiling following ISO 9001:2008 and ISO/IEC 17025:2005 quality standards.**

There are no warranties with respect to the services or results supplied, express or implied, including, without limitation, any implied warranty of merchantability or fitness for a particular purpose. Neither ATCC nor Promega is liable for any damages or injuries resulting from receipt and/or improper, inappropriate, negligent or other wrongful use of the test results supplied, and/or from misidentification, misrepresentation, or lack of accuracy of those results. Your exclusive remedy against ATCC, Promega and those supplying materials used in the services for any losses or damage of any kind whatsoever, whether in contract, tort, or otherwise, shall be, at Promega's option, refund of the fee paid for such service or repeat of the service.

The ATCC trademark and trade name, any and all ATCC catalog numbers are trademarks of the American Type Culture Collection. PowerPlex is a registered trademark of Promega Corporation. Applied Biosystems, ABI Prism and GeneMapper are registered trademarks of Life  Technologies Corporation.

**Technical questions?**

ATCC Technical Support
(800) 638-6597 / +1 703-365-2700


**Ordering questions?**

800-638-6597 or 703-365-2700


ATCC® Cell Line Authentication Service STR Profile Report

**FTA Barcode:** STRB2778

**ATCC Sales Order:** SO0533550

| Test Results for Submitted Sample | | | | | ATCC Reference Database Profile | | | |
|---|---|---|---|---|---|---|---|---|
| **Locus** | **Query Profile: TC32-pLE0567_#6** | | | | **Database Profile:** | | | |
| **D3S1358** | 15 | 16 | | | | | | |
| **TH01** | 6 | 9.3 | | | | | | |
| **D21S11** | 27 | 28 | | | | | | |
| **D18S51** | 11 | 19 | | | | | | |
| **Penta_E** | 12 | 13 | | | | | | |
| **D5S818** | 12 | 13 | | | | | | |
| **D13S317** | 10 | 12 | | | | | | |
| **D7S820** | 8 | 11 | | | | | | |
| **D16S539** | 13 | 14 | | | | | | |
| **CSF1PO** | 11 | 13 | | | | | | |
| **Penta_D** | 10 | 12 | | | | | | |
| **Amelogenin** | X | | | | | | | |
| **vWA** | 15 | 18 | | | | | | |
| **D8S1179** | 12 | | | | | | | |
| **TPOX** | 9 | 11 | | | | | | |
| **FGA** | 23 | 24 | | | | | | |
| **D19S433** | 12 | 14 | | | | | | |
| **D2S1338** | 17 | 19 | | | | | | |
| Number of shared alleles between query sample and database profile: | | | | | | | | NA |
| Total number of alleles in the database profile: | | | | | | | | NA |
| Percent match between the submitted sample and the database profile: | | | | | | | | NA |

*The allele match algorithm compares the 8 core loci plus amelogenin only, even though alleles from all loci will be reported when available.*

**NOTE:** *Loci highlighted in grey (8 core STR loci plus Amelogenin) can be made public to verify cell identity. In order to protect the identity of the donor,* ***please do not publish*** *the allele calls from all the STR loci tested.*
*Electropherograms showing raw data are attached.*

**Explanation of Test Results**
Cell lines with 80% match are considered to be related; i.e., derived from a common ancestry. Cell lines with between a 55% to 80% match require further profiling for authentication of relatedness.

[X] The submitted sample profile is human, but not a match for any profile in the ATCC STR database.

[ ] The submitted profile is an exact match for the following ATCC human cell line(s) in the ATCC STR database (8 core loci plus Amelogenin):

[ ] The submitted profile is similar to the following ATCC human cell line(s):

[ ] An STR profile could not be generated.

**Additional Comments:**

Submitted sample, STRB2778 (TC32-pLE0567_#6), shows similarities to ATCC cell line HTB-133 (T-47D) however the cell lines appear to be unrelated, see addendum. The profile for the submitted sample is an exact match to the STR profile for the (TC-32) cell line that is listed on the ExPASy website. https://web.expasy.org/cellosaurus

| | |
|---|---|
| e-Signature, Technician: | snicholson 11/24/2020 |
| e-Signature, Reviewer: | gsykes 11/24/2020 |

CELL LINE AUTHENTICATION SERVICE ATCC VERIFIED EST. 1925

### Cell Line Authentication Service
STR Profile Report

**FTA Barcode:** STRB2778

**ATCC Sales Order:** SO0533550

**Addendum: Comparative Output from the ATCC STR Profile Database**

| % Match | ATCC® Cat. No. | Designation | D5S818 | D13S317 | D7S820 | D16S539 | vWA | TH01 | AMEL | TPOX | CSF1PO |
|---|---|---|---|---|---|---|---|---|---|---|---|
| 100 | STRB2778 | TC32-pLE0567_#6 | 12,13 | 10,12 | 8,11 | 13,14 | 15,18 | 6,9.3 | X | 9,11 | 11,13 |
| 80 | HTB-133 | T-47D; Breast Cancer; Human | 12 | 12 | 11 | 10 | 14 | 6 | X | 11 | 11,13 |

**Definitions of terms used in this report:**

**Peak Area Difference (PAD):**
Refers to a heterozygous peak imbalance.
Two alleles at a single locus should amplify in a similar manner; and therefore produce peaks of similar height and area. Peaks which are above threshold (50 rfu) but are not of similar area, within 50% of each other, are referred to as a PAD. Due to their nature cell lines do not amplify in the same manner as a sample taken from a fresh buccal swab. PAD is far more common in cell line samples.

**Stutter:**
A stutter peak is a small peak which occurs immediately before the true peak. It is defined as being a single repeat unit smaller than the true peak. The stutter peak should be less than 15% of the true peak. The stutter is caused by the polymerase.

**+4 Peak:**
A +4 is similar to a stutter but occurs immediately after the true peak. A stutter peak should be less than 5% for a homozygous and 10% for a heterozygous.

**Below Threshold Peak(s):**
Cell lines can produce unusual profiles and occasionally a peak will amplify poorly and be below threshold. Where we find a below threshold peak which we believe is valid we indicate it as a below threshold peak. Our cell line analysis criteria, Homozygous and Heterozygous peaks must be equal to or above the set height threshold for it to be considered a true peak.

**Ladder/ Off Ladder Peak(s):**
The allelic ladder consists of most or all known alleles in the population and allows for precise assignment of alleles. Those which do not align are termed ‘off ladder.

**Artifact:**
A non-allelic product of the amplification process, an anomaly of the detection process, or a by-product of primer synthesis

**Pull-up:**
A term used to describe when signal from one dye color channel produces artificial peaks in another, usually adjacent, color.

**Spike:**
An extraneous peak resulting from dust, dried polymer, an air bubble, or an electrical surge.
**Dye blob:**
Free dye not coupled to primer that can be injected into the capillary (A known and documented dye blob is often found at the D3S1358 locus.)

Project: 20201119-WK

| Sample Name | SQ | CGQ |
|---|---|---|
| STRB2778_TC32-pLE0567_#6 | | |

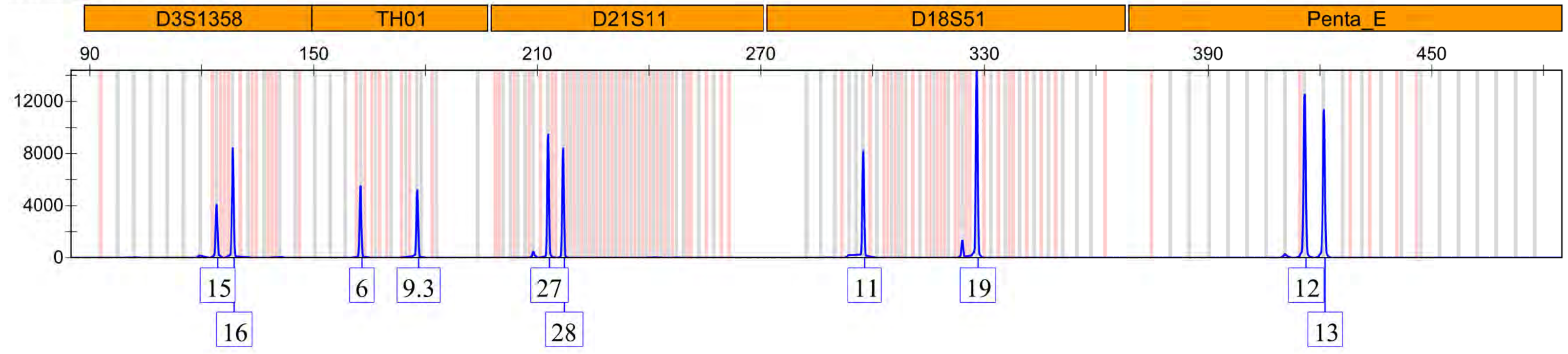

STRB2778_TC32-pLE0567_#6

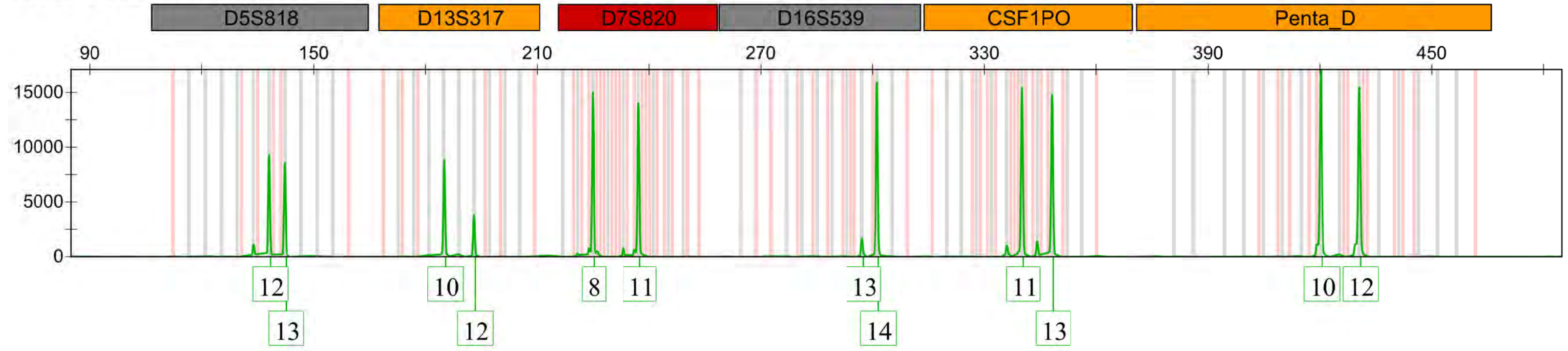

| Sample Name | SQ | CGQ |
|---|---|---|
| STRB2778_TC32-pLE0567_#6 | | |

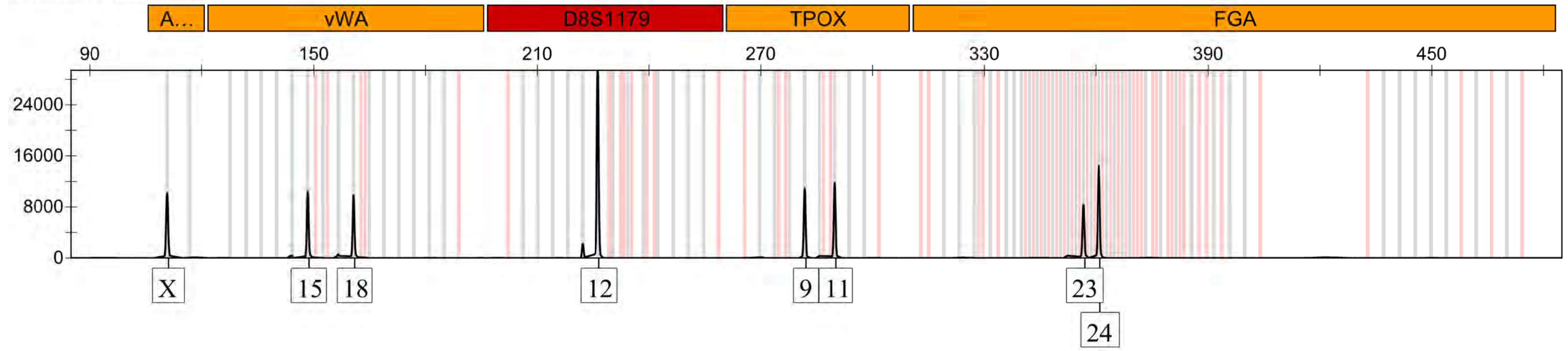

STRB2778_TC32-pLE0567_#6

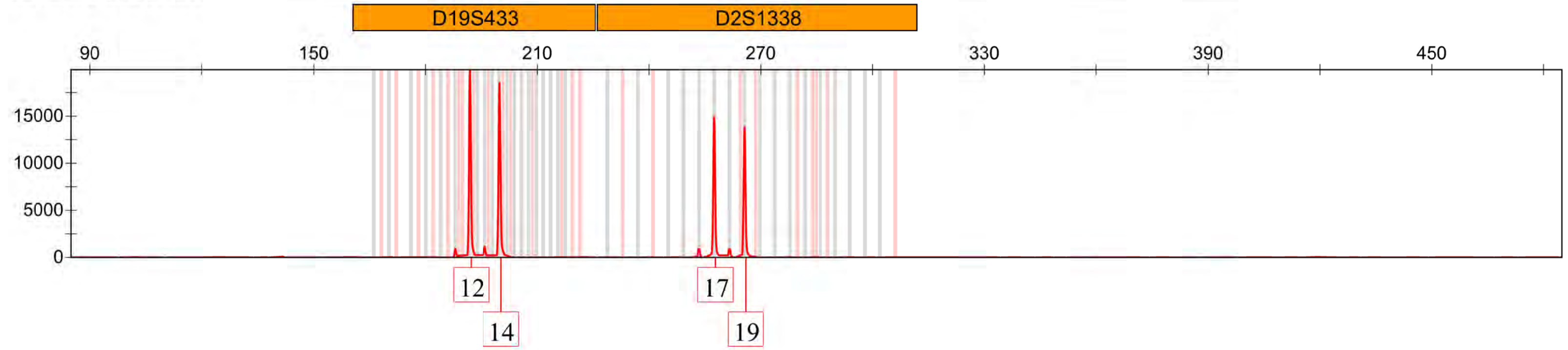

Supplementary Figure 3: Plasmid maps

#### (1) pCDNA5/FRT miniAID-EGFP (Used as a control to optimize electroporation conditions for cell lines of interest):

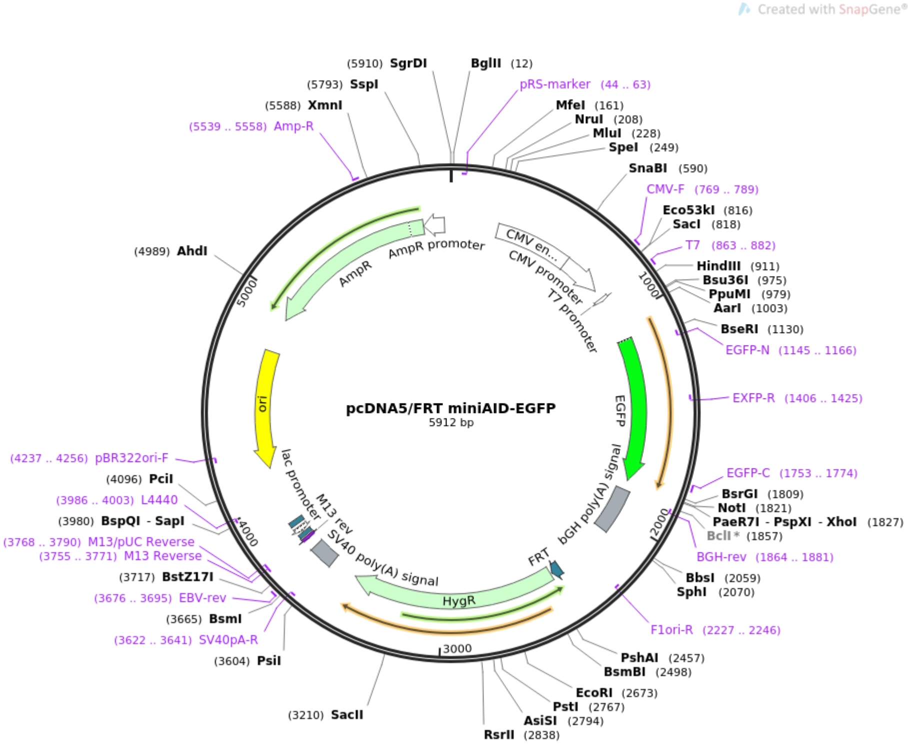

#### (2) Lenti-SpCas9-2A-mCherry_EWSR1-IVT-1001 (sgRNA targeting exon 1 of EWSR1)

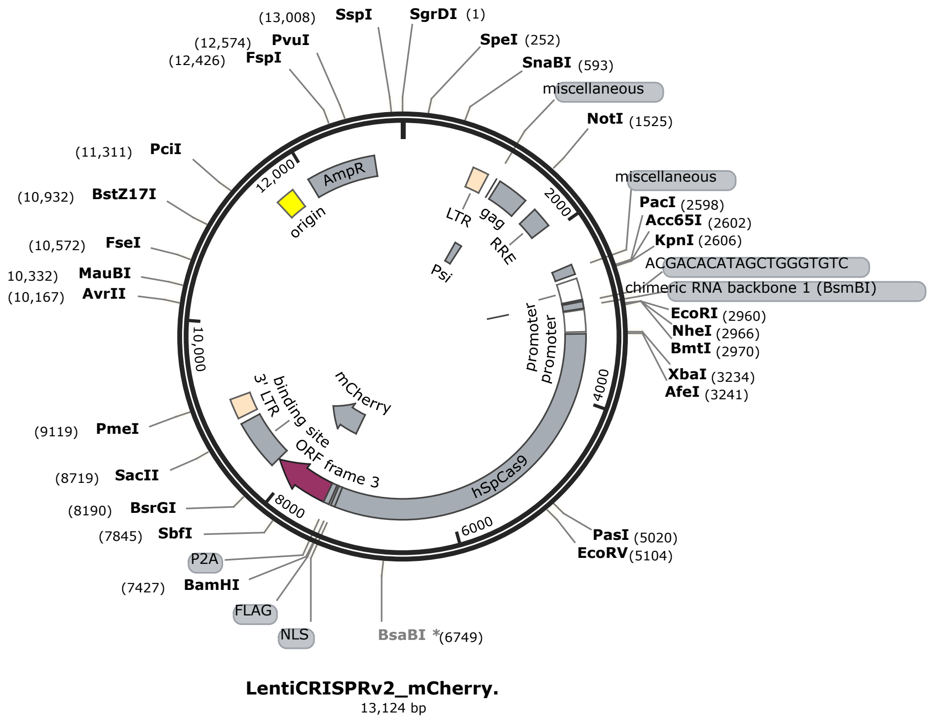

#### (3) Lenti-SpCas9-2A-mCherry_EWSR1-IVT-1002 (Second sgRNA targeting exon 1 of EWSR1)

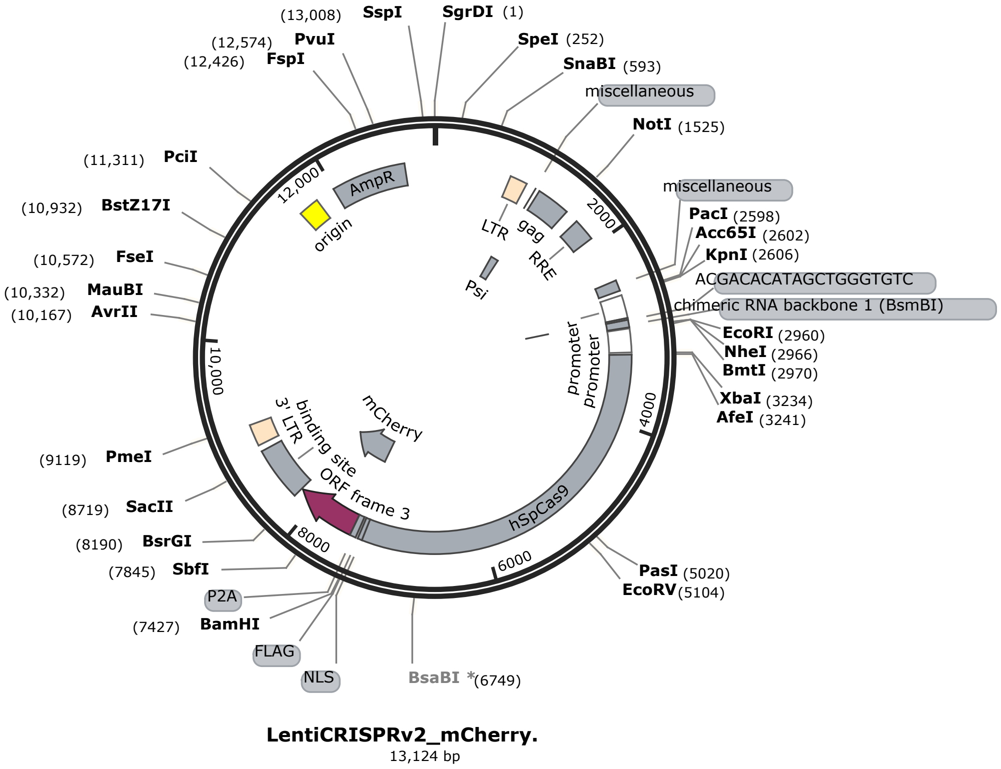

#### (4) Donor-EWSR1-N-term-NeonGreen (pCE0567 – Plasmid carrying the donor template and NeonGreen insertion)

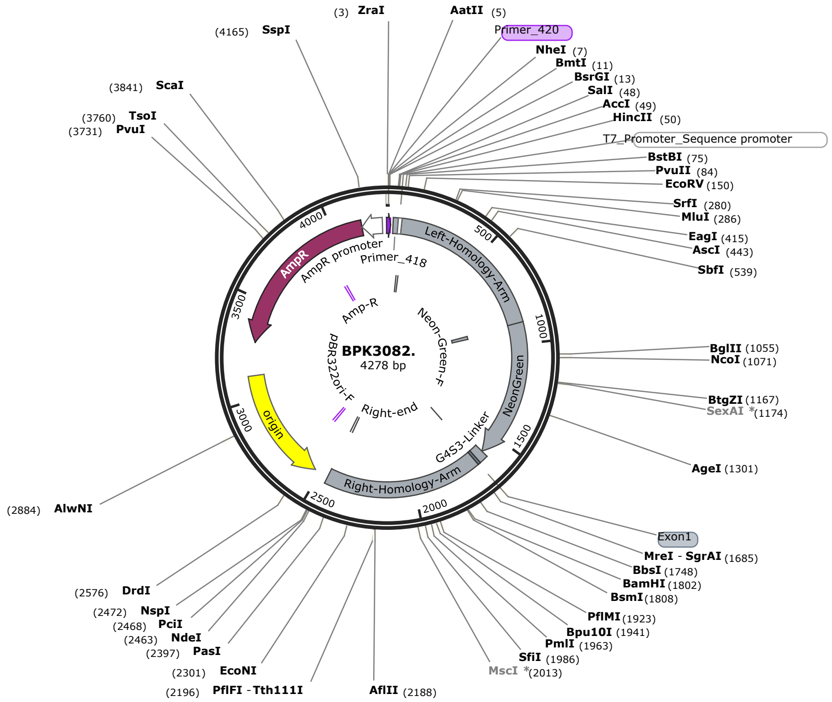

(5) **Donor-EWSR1-N-term-NeonGreen-2A-HiBiT(pCE0568 – Plasmid carrying the donor template and NeonGreen with HiBit tag)**

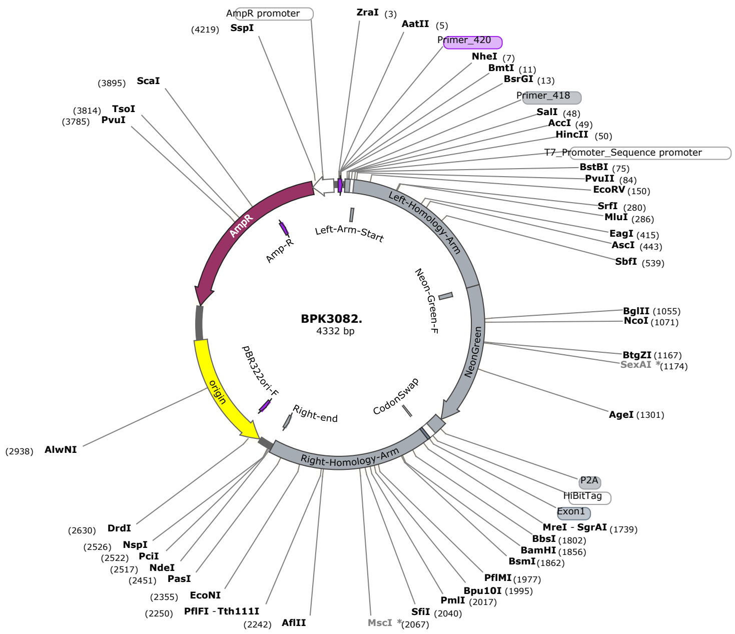
